# The miR-124-AMPAR pathway connects polygenic risks with behavioral changes shared between schizophrenia and bipolar disorder

**DOI:** 10.1101/2021.10.19.465053

**Authors:** Ho Namkung, Hiroshi Yukitake, Daisuke Fukudome, Brian J. Lee, Gianluca Ursini, Shravika Lam, Suvarnambiga Kannan, Atsushi Saito, Minae Niwa, Kamal Sharma, Peter Zandi, Hanna Jaaro-Peled, Koko Ishizuka, Nilanjan Chatterjee, Richard Huganir, Akira Sawa

**Author notes:** To whom correspondence should be addressed: Akira Sawa.

## Abstract

Schizophrenia (SZ) and bipolar disorder (BP) are highly heritable major psychiatric disorders that share a substantial portion of genetic risk as well as their clinical manifestations. This raises a fundamental question of whether, and how, common neurobiological pathways translate their shared polygenic risks into shared clinical manifestations. The present study shows the miR-124-AMPAR pathway as a key common neurobiological mediator that connects polygenic risks with behavioral changes shared between these two psychotic disorders. We discovered upregulation of miR-124 in biopsied neuronal cells and postmortem prefrontal cortex from both SZ and BP patients, implying its role not only as a biomarker, but also as a pathophysiological mediator. Intriguingly, the upregulation is associated with the polygenic risks shared between these two disorders. Seeking mechanistic dissection, we generated a mouse model that upregulates miR-124 in the medial prefrontal cortex, which includes brain regions homologous to sub-regions of the human prefrontal cortex. We demonstrated that upregulation of miR-124 increases GRIA2-lacking calcium permeable-AMPARs and perturbs AMPAR-mediated excitatory synaptic transmission, leading to deficits in the behavioral dimensions shared between SZ and BP.

## INTRODUCTION

Psychiatric conditions are currently a major social and medical burden. Despite considerable efforts for their prevention and treatment in clinical studies, there have been few advances mainly due to our shortage in understanding of their neurobiological mechanisms. Recent advances in psychiatric genomics have illuminated polygenic contributions of common genetic variants to psychiatric conditions, with each having a small effect size (Consortium et al., 2020; Gandal et al., 2016; Geschwind and Flint, 2015; Mullins et al., 2021; O’Donovan and Owen, 2016). Translating these genetic insights into clinical manifestations through intermediate neurobiological pathways may provide a foundation for understanding the pathophysiological mechanisms of these conditions (Namkung et al., 2018). Constructing a novel research framework that realizes this goal may be indispensable to reshaping clinical psychiatry in a neurobiological mechanism-based manner.

Schizophrenia (SZ) and bipolar disorder (BP) are highly heritable major psychiatric disorders that share a substantial number of common genetic risk factors (Geschwind and Flint, 2015; Lee SH, 2013; Ruderfer et al., 2018). Emerging evidence suggests convergence of these shared genetic variants onto molecular pathways which include those underlying synaptic function (Gandal et al., 2018; Ruderfer et al., 2018). SZ and BP also have overlapping deficits in multiple behavioral dimensions from a clinical viewpoint (Ruderfer et al., 2018). However, it has remained elusive whether, and how, their shared polygenic risks converge onto specific molecular pathways that regulate synaptic function, which in turn affects shared behavioral dimensions.

MicroRNAs (miRNAs) are a class of small non-coding RNAs that regulate gene expression at the post-transcriptional level (Bartel, 2004; He and Hannon, 2004). MicroRNA-124 (miR-124) is the most abundant and well-conserved brain-specific miRNA (Krichevsky et al., 2003; Lagos-Quintana et al., 2002; Sanuki et al., 2011). Consistent with its high expression in neurons but low in glial cells (Jovicic et al., 2013), it has been shown to play a key role in neuronal differentiation and neurogenesis (Cheng et al., 2009; Yoo et al., 2011). Emerging evidence also emphasizes the significance of this neuron-enriched miRNA (miR-124) in regulating synaptic function, implying its potential engagement in brain disorders (Gascon et al., 2014; Hou et al., 2015; Rajasethupathy et al., 2009; Sanuki et al., 2011; Wang et al., 2018). In particular, it has been shown to support the regulation of AMPAR-mediated excitatory synaptic transmission (Gascon et al., 2014; Ho et al., 2014; Hou et al., 2015; Wang et al., 2018). However, it is still unknown whether, and how, miR-124-mediated synaptic function drives behavioral dimensions relevant to SZ and BP.

Supported by the discovery on upregulation of miR-124 in biopsied neuronal cells, followed by its validation in postmortem prefrontal cortex, from both SZ and BP patients and its association with their shared polygenic risks, we explored a specific miR-124 pathway that connects polygenic risks with behavioral changes shared between these two psychotic disorders. A mouse model that upregulates miR-124 in the medial prefrontal cortex, which includes brain regions homologous to sub-regions of the human prefrontal cortex, demonstrated that upregulation of miR-124 increases GRIA2-lacking calcium permeable-AMPARs and perturbs AMPAR-mediated excitatory synaptic transmission, leading to deficits in the behavioral dimensions shared between SZ and BP. Taken altogether, this study provides not only a novel neurobiological insight into how miR-124-mediated synaptic function impacts behavioral dimensions of psychiatric relevance, but also a new translational research framework in which human and animal studies are combined to address a disease mechanism that bridges across pathogenesis, pathophysiology, and phenotypic/clinical manifestations.

## RESULTS

### Upregulation of microRNA-124 (miR-124) discovered in biopsied neuronal cells from living patients with SZ and those with BP

The use of biopsied tissues from patients with active symptomatic manifestation has been regarded as indispensable not only for discovering reliable biomarkers, but also for studying pathophysiological mechanisms in many medical conditions, such as several types of cancers, liver disorders such as hepatitis, and lung disorders including idiopathic pulmonary fibrosis (Hunninghake et al., 2001; Iacobellis and Bianco, 2011; Mattie et al., 2006; Rockey et al., 2009; Siravegna et al., 2017). However, due to the difficulty in accessing the brain in living patients, this strategy has not been utilized in psychiatry (Gamo and Sawa, 2014). To overcome this limitation, investigators have recently highlighted the utility of olfactory epithelium (OE)-derived neuronal cells (henceforth, ‘olfactory neuronal cells’), easily accessible through a nasal biopsy, as a promising surrogate that captures neuron-relevant molecular signatures in the course of functional impairment in living patients with major psychiatric disorders (Arnold et al., 1998; Borgmann-Winter et al., 2016; Doostparast Torshizi et al., 2019; Evgrafov et al., 2020; Feron et al., 1998; Hahn et al., 2005; Kano et al., 2013; Lavoie et al., 2017; Mackay-Sim, 2012; Rhie et al., 2018). Thus, we initiated the present study by analyzing gene expression profiles of olfactory neuronal cells from patients with schizophrenia (SZ) compared to those from healthy controls (CON): a microarray analysis identified 854 dysregulated genes in SZ, with 330 genes being upregulated and 524 genes being downregulated (**Figure S1**). We next performed a pathway enrichment analysis with these dysregulated genes in order to identify key pathways that were perturbed in SZ (**Figure 1A**). While most of the top-listed pathways, such as the glutathione metabolic pathway (Do et al., 2000), were reasonably expected in the context of SZ pathophysiology, it was unexpectedly discovered that putative targets of microRNA-124 (miR-124), one of the most abundant microRNAs in the brain (Sanuki et al., 2011), were significantly enriched in genes displaying decreased expression in SZ.

**Figure 1.**
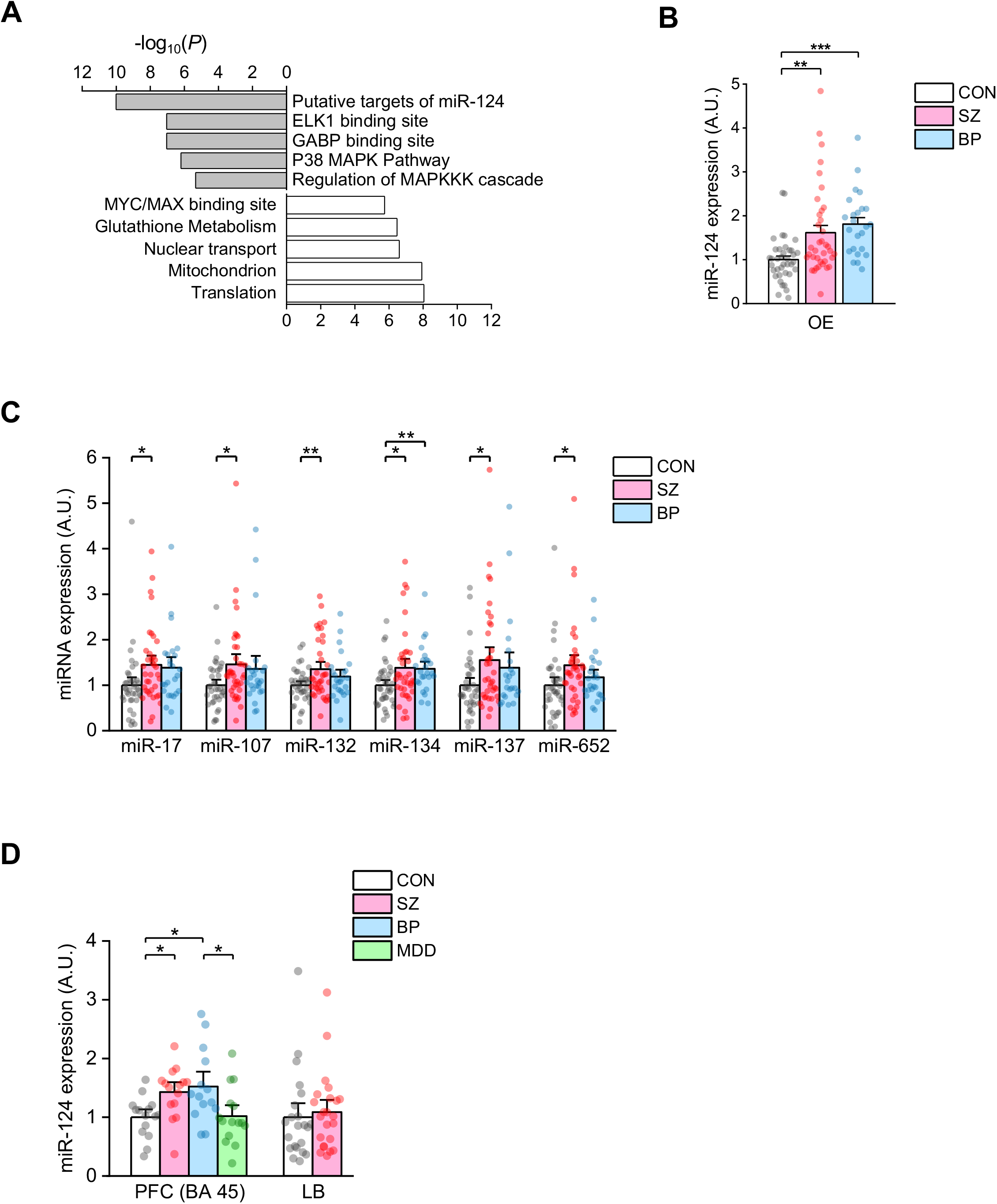
MiR-124 is significantly upregulated in biopsied neuronal cells and the postmortem prefrontal cortex (PFC) from patients with schizophrenia (SZ) and those with bipolar disorder (BP). **(A)** Pathway enrichment analysis found that putative targets of miR-124 were significantly downregulated in olfactory epithelium (OE)-derived neuronal cells (olfactory neuronal cells) from patients with SZ, implying upregulation of miR-124. (**B**) Upregulation of miR-124 was validated by quantitative real-time polymerase chain reaction (qRT-PCR) in olfactory neuronal cells from SZ and BP patients, compared to healthy controls (CON) (*N*_CON_=38, *N*_SZ_=37, *N*_BP_=24; Welch’s ANOVA, *F*_2,52.335_=14.100, *p*=1.3×10^-5^; *post hoc* Games-Howell test). **(C)** Alterations of miR-17, −107, −132, −134, −137, and −652 were detected in olfactory neuronal cells from SZ and/or BP patients. miR-17 (*N*_CON_=39, *N*_SZ_=37, *N*_BP_=25; one-way ANOVA, *F*_2,98_=3.246, *p*=0.043; *post hoc* Tukey’s HSD test). miR-107 (*N*_CON_=38, *N*_SZ_=38, *N*_BP_=25; one-way ANOVA, *F*_2,98_=3.393, *p*=0.038; *post hoc* Tukey’s HSD test). miR-132 (*N*_CON_=39, *N*_SZ_=38, *N*_BP_=25; Welch’s ANOVA, *F*_2,55.228_=4.552, *p*=0.015; *post hoc* Games-Howell test). miR-134 (*N*_CON_=39, *N*_SZ_=38, *N*_BP_=25; Welch’s ANOVA, *F*_2,59.049_=5.428, *p*=0.007; *post hoc* Games-Howell test). miR-137 (*N*_CON_=38, *N*_SZ_=37, *N*_BP_=23; Welch’s ANOVA, *F*_2,49.415_=3.811, *p*=0.029; *post hoc* Games-Howell test). miR-652 (*N*_CON_=39, *N*_SZ_=38, *N*_BP_=25; one-way ANOVA, *F*_2,99_=3.117, *p*=0.049; *post hoc* Tukey’s HSD test). **(D)** MiR-124 was also found to be upregulated in a sub-region (BA 45) of the PFC from both SZ and BP (but not MDD) patients, compared to that from CON (*N*_CON_=15, *N*_SZ_=15, *N*_BP_=14, *N*_MDD_=15; one-way ANOVA, *F*_3,58_=4.695, *p*=0.005; *post hoc* Tukey’s HSD test). However, no alteration was detected in lymphoblasts (LB) from SZ, compared to those from CON (*N*_CON_=22, *N*_SZ_=23; two-tailed Student’s *t*-test, *t*_43_=-0.415, *p*=0.754). Bar graph expressed as mean ± SEM. \**p* < 0.05, \*\**p* < 0.01, *** *p* < 0.001. More detailed statistical information is in **Table S2**.

We therefore hypothesized that miR-124 is significantly upregulated in olfactory neuronal cells from SZ patients. Given that SZ and bipolar disorder (BP) share a substantial portion of molecular pathways underlying their pathophysiology (Gandal et al., 2018; Shao and Vawter, 2008), we further hypothesized that miR-124 is also dysregulated in olfactory neuronal cells from BP patients. To validate these hypotheses, we measured miR-124 expression levels in olfactory neuronal cells from CON, SZ, and BP subjects by performing quantitative real-time polymerase chain reaction (qRT-PCR), and subsequently found that miR-124 is significantly upregulated in olfactory neuronal cells from both SZ and BP patients, compared to those from CON (**Figure 1B**). We also examined, in olfactory neuronal cells, several other microRNAs which have been reportedly dysregulated in SZ and/or BP patients (Beveridge et al., 2010; Kim et al., 2010; Santarelli et al., 2011; Siegert et al., 2015). We detected disease-relevant changes in all microRNAs we tested, including miR-17, −107, −132, −134, and −652 (**Figure 1C**), with their alterations in olfactory neuronal cells being consistent with those previously found in the postmortem prefrontal cortex (PFC) (Beveridge et al., 2010; Kim et al., 2010; Santarelli et al., 2011). In addition, we found an alteration of miR-137 in olfactory neuronal cells from SZ patients (**Figure 1C**), consistent with what was reported in induced human neurons (Siegert et al., 2015). These data indicate the utility of biopsied olfactory neuronal cells to study dynamic changes of microRNAs in living patients with psychotic disorders. Notably, the magnitude and significance of the observed alterations were most robust in miR-124 compared with those of other microRNAs we tested, implying its potential role not only as a reliable biomarker, but also as a pathophysiological mediator (**Figures 1B and 1C**).

The human PFC has been significantly implicated in the pathophysiology of both SZ and BP (Ellison-Wright and Bullmore, 2010; Gandal et al., 2018; Konopaske et al., 2014; Shao and Vawter, 2008). Gene expression profiling has showed that molecular signatures relevant to the PFC are resonably represented in olfactory neuronal cells (Doostparast Torshizi et al., 2019). Together, we hypothesized that olfactory neuronal cells are a promising surrogate that captures neuron-relevant molecular signatures in the PFC, and that our observation in olfactory neuronal cells may be a recapitulation of important molecular changes in PFC. Thus, we first examined published data of RNA sequencing from the postmortem PFC (Fromer et al., 2016). Consistent with the finding from olfactory neuronal cells, pathway enrichment analysis identified that putative targets of miR-124 were significantly downregulated in the SZ postmortem dorsolateral PFC (DLPFC: BA9 and BA46) (**Table S1**), implying a potential upregulation of miR-124. Indeed, the primary miR-124-2 [pri-miR-124-2, also known as miR-124-2 host gene (miR-124-2HG)] and precursor miR-124-2 (pre-miR-124-2) were found to be significantly upregulated in the postmortem DLPFC from SZ and BP, respectively, in an independent cohort (Jaffe et al., 2018). (**Figures S2A and S2B**). Furthermore, recent genome-wide association studies (GWAS) have indicated that single nucleotide polymorphisms (SNPs) near or within the pri-/pre-miR-124-2 and pri-/pre-miR-124-1 gene loci have been shown to confer risks for SZ and BP, respectively (Consortium et al., 2020; Mullins et al., 2021) (**Figures S2C and S2D**). We could access another sub-region of the postmortem PFC (BA45, adjacent to the DLPFC) from patients with SZ, BP, and major depressive disorder (MDD), as well as CON subjects (Torrey et al., 2000). We thus conducted qRT-PCR and detected an upregulation of miR-124 only in SZ and BP, but not in MDD, when compared with CON (**Figure 1D**). Taken together, miR-124 may be dysregulated on a relatively broad range of the PFC specifically in SZ and BP. In contrast, miR-124 expression levels were not significantly different between CON and SZ lymphoblasts (LB), implying that dysregulation of miR-124 may be a molecular signature that is specific to neural tissues (**Figure 1D**).

### MiR-124 serves as a key pathophysiological mediator that connects polygenic risks with clinical manifestations shared between SZ and BP

SZ and BP share substantial polygenic risk factors, which may underlie their common molecular pathways (Gandal et al., 2018; Purcell et al., 2009; Shao and Vawter, 2008). It is also known that SZ and BP share some clinical manifestations and aberrant behavioral dimensions (Pearlson, 2015; Ruderfer et al., 2018). Thus, we hypothesized that miR-124 may serve as a key mediator linking the shared polygenetic risks and common clinical feature(s) that can exist in both SZ and BP. To this end, we calculated polygenic risk scores (PRSs) in our independent samples of European ancestry using four GWAS summary statistics (Ruderfer et al., 2018): (i) combined SZ and BP cases vs. combined SZ and BP controls (SZ+BP PRSs); (ii) SZ cases vs. SZ controls (SZ PRSs); (iii) BP cases vs. BP controls (BP PRSs); and (iv) SZ cases vs. BP cases (SZ vs. BP PRSs) (**Figure 2A**). Applying logistic regression analysis, we found that polygenic risks shared between SZ and BP (SZ+BP PRSs) account for a substantial proportion (14%) of the variance in case-control status (either SZ or BP vs. CON) (**Figure 2B**). We next tested whether the shared polygenic risks are also associated with miR-124 expression. Linear regression analysis showed that SZ+BP PRSs significantly predict miR-124 expression, accounting for 17.4% of its variance, at the *p*-value threshold where the PRSs best account for the diagnosis of either SZ or BP (**Figure 2C**).

**Figure 2.**
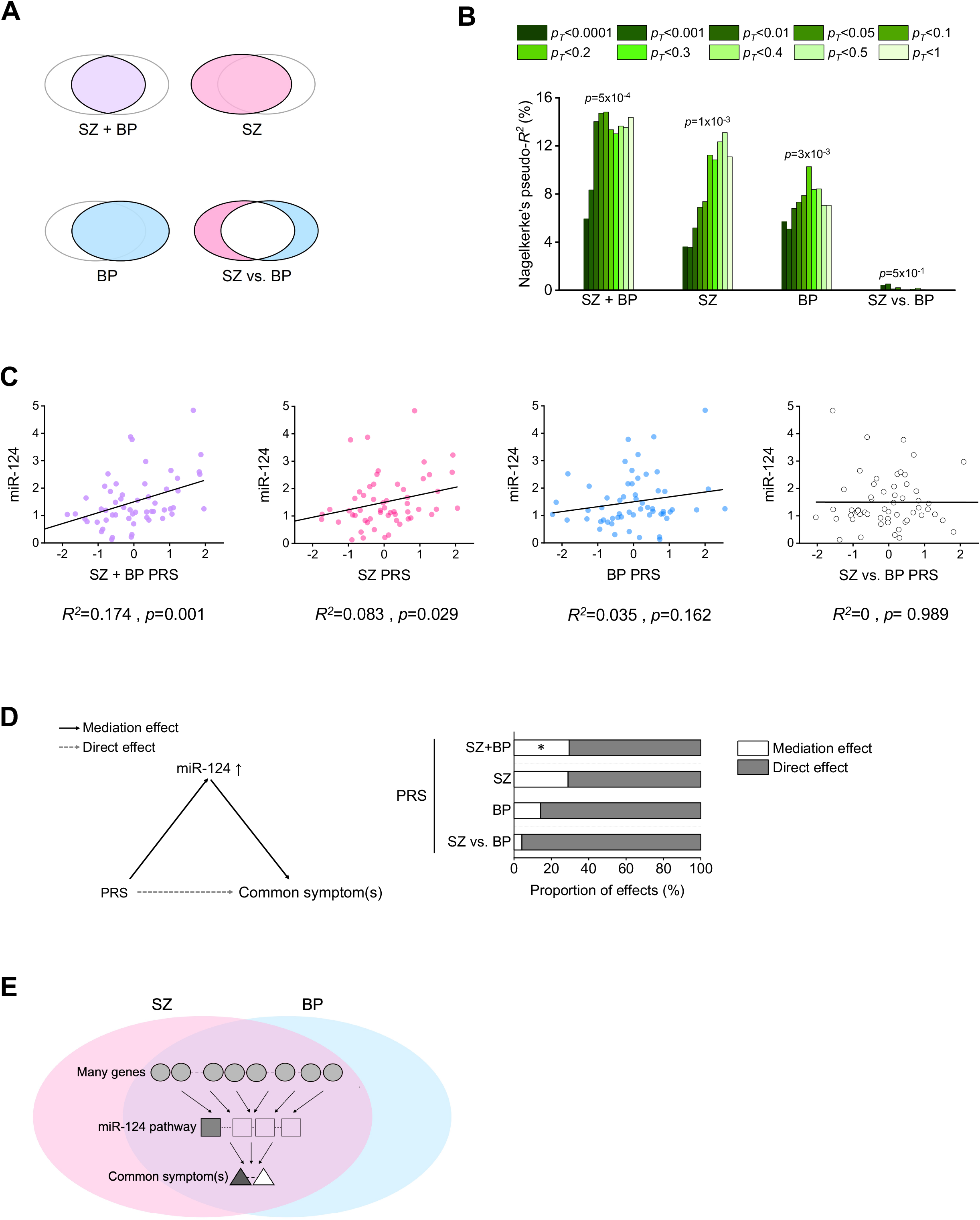
MiR-124 serves as a key mediator linking the polygenetic risks shared between SZ and BP to common clinical feature(s) that can exist in both SZ and BP. **(A)** Genome-wide polygenic risk scores (PRSs) for SZ+BP, SZ, BP, SZ vs. BP were calculated in our independent samples of European ancestry to address shared or disease-specific polygenic contributions to phenotypic feature(s) that can commonly exist in SZ and BP. **(B)** Case-control status (either SZ or BP vs. CON) is best predicted by SZ+BP PRSs, and also well predicted by SZ PRSs or BP PRSs, which is in contrast to poor prediction by SZ vs. BP PRSs (*N*_CON_=45, *N*_SZ+BP_=63; logistic regression analysis). **(C)** Both SZ+BP and SZ PRSs significantly predict miR-124 expression, accounting for 17.4% and 8.3% of the variance of miR-124 expression, respectively, at the *p*-value thresholds where the PRSs best account for the diagnosis of either SZ or BP. However, PRSs for BP and SZ vs. BP do not predict miR-124 expression significantly, accounting only for 3.5% and 0% of the variance of miR-124 expression, respectively (*N*=57; linear regression analysis). **(D)** MiR-124 functions as an important mediator between the shared polygenic contributions and the diagnosis of either SZ or BP. SZ+BP PRSs (*N* 57; mediation analysis using a bootstrapping method with 5,000 iterations, proportion mediated (%) = 30.7, 95% CI [0.0096, 1.44], *p*=0.042). **(E)** This schematic illustrates that polygenic risks shared between SZ and BP converge onto dysregulation of miR-124 in the common pathophysiology that may in turn contribute to clinical feature(s) commonly observed in SZ and BP, such as psychosis. * *p* < 0.05. More detailed statistical information is in **Table S2.**

We then tested whether miR-124 serves as a key mediator between the shared polygenic risks and either diagnosis. To address this question, we applied mediation analysis, a method which is widely used across many disciplines to explore pathways via mediators (MacKinnon et al., 2007). We found that miR-124 is a significant mediator in the effects of shared polygenic risks on either diagnosis, accounting for 30.7% of the total effect (**Figure 2D**). In summary, these results indicate that miR-124 functions as an important mediator between the shared polygenic contributions and two diseases (both SZ and BP) (**Figure 2E**). Complementing these results, we also applied Mendelian randomization, and found a potential causal role of miR-124 in either SZ or BP by using the pre-miR-124-1-associated SNP rs28548961 as an instrument (**Figure S3**). Given that SZ and BP share clinical and behavioral manifestations to some extent, these genetic data may represent the biological scenario that the pathway involving miR124 links shared polygenic risks to neuro-behavioral changes shared in SZ and BP. We decided to address the validty of this notion in animal studies described below.

### Overexpression of miR-124 in the PFC leads to deficits in the behavioral dimensions relevant to both SZ and BP, and perturbs excitatory synaptic transmission

Given that miR-124 is upregulated in olfactory neuronal cells and PFC from the patients, we decided to further explore a potential mechanism by which the upregulation of miR-124 in the PFC causally drives phenotypes relevant to both SZ and BP. Accordingly, we generated a mouse model overexpressing miR-124 in neurons of the medial PFC (mPFC) by bilaterally delivering adeno-associated viruses (AAVs) that express miR-124 under the synapsin promoter. The mPFC was particularly targeted because it includes brain regions homologous to many sub-regions of the human PFC (Heilbronner et al., 2016; Hoover and Vertes, 2007; Laubach et al., 2018; Wise, 2008). While comparative neuroanatomy between humans and mice still involves some debate, we believe that our choice of mPFC is reasonable given the brain regions where we observed the alteration of miR-124 (**Figure S4**) (Heilbronner et al., 2016; Hoover and Vertes, 2007; Laubach et al., 2018; Wise, 2008). We assayed the miR-124-overexpressing mice with a behavioral test battery that addresses dimensions commonly impaired in many cases of SZ and BP (**Figure S5A**). As a result, we identified behavioral deficits in social interaction and psychostimulant sensitivity, both of which are relevant to SZ and BP (**Figures S5B-S5E**) (Lieberman et al., 1987; Mancuso et al., 2011).

We then asked whether these behavioral phenotypes are dependent on neuronal subtypes. Given that the mouse mPFC innervates brain regions implicated in social interaction and psychostimulant sensitivity, such as the nucleus accumbens and the ventral tegmental area (Deng et al., 2010; Murugan et al., 2017), we focused on investigating excitatory neurons in the mPFC, which are the dominant source of cortical outputs (Brown and Hestrin, 2009; Spruston, 2008). We therefore overexpressed miR-124 in excitatory neurons of the mPFC by bilaterally delivering AAVs that express miR-124 fused to EmGFP under the CaMKIIα promoter (**Figure 3A**). As a result, miR-124 was increased approximately 2.4-fold in miR-124-overexpressing excitatory neurons, which is of a comparable degree to the endogenous upregulation of miR-124 in olfactory neuronal cells from SZ and BP patients (**Figure 3B**). The excitatory neuron-specific overexpression of miR-124 reproduced the behavioral phenotypes induced by neuronal overexpression of miR-124 and showed even more significant changes (**Figures 3C and 3D**), implying that excitatory neurons in the mPFC may have a dominant role in driving such behavioral phenotypes. We next sought to address a synaptic mechanism that may underlie the behavioral deficits. We recorded miniature excitatory postsynaptic currents (mEPSCs) in layer V pyramidal neurons of the mPFC, the main output neurons of the mPFC (Brown and Hestrin, 2009). The layer V pyramidal neurons overexpressing miR-124 displayed a significant increase in the amplitude of mEPSCs with no concurrent change in the frequency of mEPSCs, indicating alterations in AMPA receptor (AMPAR)-mediated synaptic transmission (**Figure 3E**).

**Figure 3.**
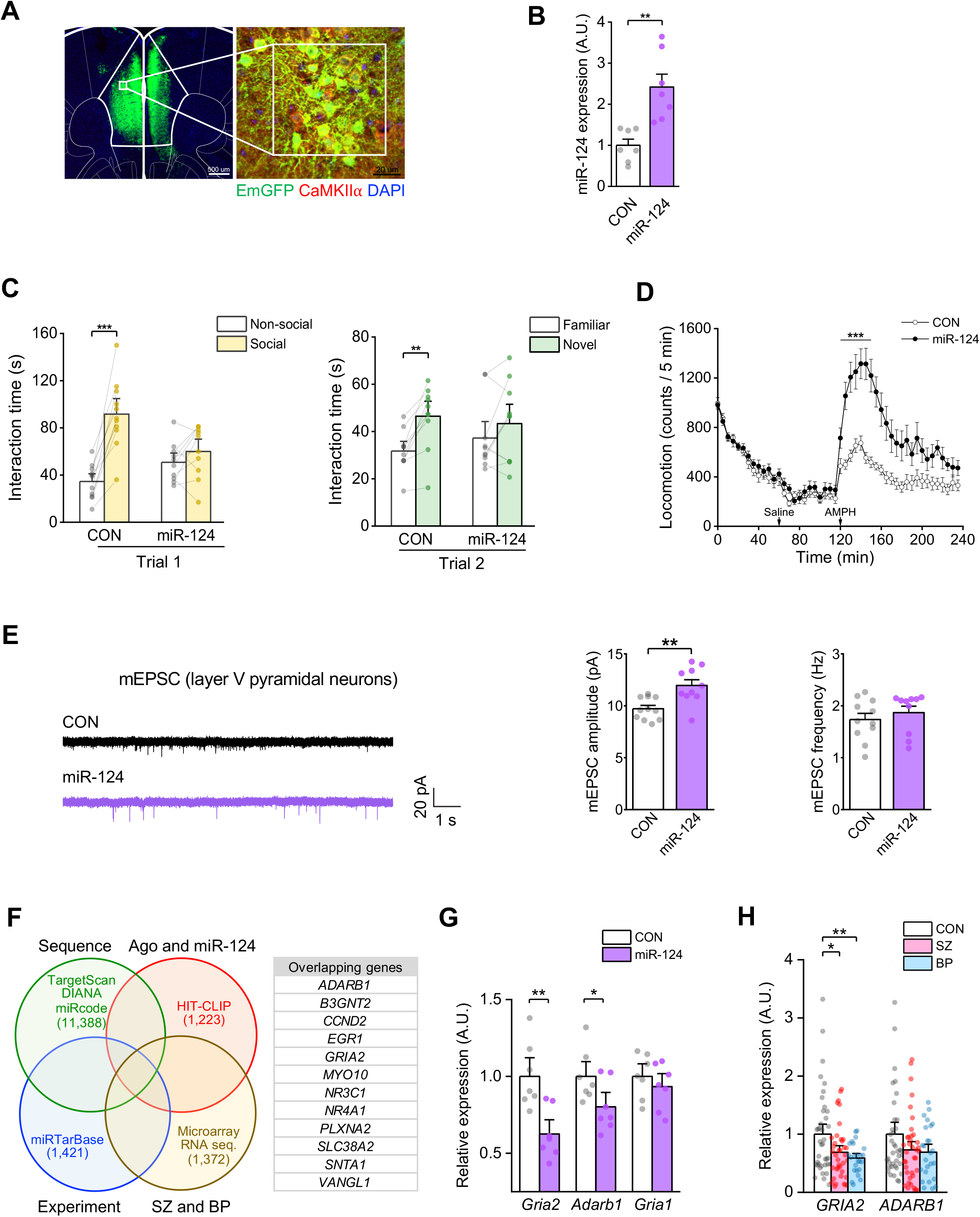
Overexpression of miR-124 in excitatory neurons of the mouse mPFC causes behavioral and synaptic deficits. **(A)** Adeno-associated virus (AAV)-mediated overexpression of miR-124 or miR-control in mPFC excitatory neurons was validated by immunostaining of CaMKIIa and EmGFP that was fused to miR-124 or miR-control. **(B)** The mPFC excitatory neurons infected with miR-124 or miR-control (CON) AAV were isolated by fluorescence-activated cell sorting (FACS), and the subsequent qRT-PCR detected that miR-124 was increased 2.4-fold in miR-124 AAV-infected excitatory neurons compared with CON AAV-infected ones (*N*_CON_=7, *N*_miR-124_=7; two-tailed Student’s *t*-test, *t*_12_=-4.095273, *p*=0.0015). **(C)** Overexpression of miR-124 in mPFC excitatory neurons caused behavioral deficits in both sociability (Trial 1, left) and social novelty recognition (Trial 2, right) in three-chamber social interactions tests. Sociability phase of trial 1 (*N*_CON_=11, *N*_miR-124_=10; two-way mixed ANOVA, AAV x Chamber, *F*_1,19_=21.5188, *p*=0.0001; *post hoc* within-group comparisons using two-tailed paired Student’s *t*-test). Social novelty recognition phase of trial 2 (*N*_CON_=11, *N*_miR-124_=10; two-way mixed ANOVA, AAV x Chamber, *F*_1,42_=6.9918, *p*=0.016; *post hoc* within-group comparisons using two-tailed paired Student’s *t*-test). **(D)** Overexpression of miR-124 in mPFC excitatory neurons caused locomotor hypersensitivity to a psychostimulant, amphetamine (AMPH). The between-group contrast was most evident for the first 30 min after the *i.p.* injection of AMPH (*N*_CON_=14, *N*_miR-124_=14; two-way mixed ANOVA with the Greenhouse-Geisser correction, AAV x Time, *F*_2.226,57.866_=9.065, *p*=2.3410^-4^; *post hoc* two-tailed paired *t*-test for each time point). **(E)** Representative traces of mEPSC recordings in mPFC layer V pyramidal neurons in brain slices from mice infected with miR-124 or CON AAV (left). The miR-124-overexpressing layer V pyramidal neurons showed significant increase in the amplitude of mEPSCs (middle; *N*_CON_=11, *N*_miR-124_=10; two-tailed student’s *t*-test, *t*_19_=-3.590, *p*=0.002). However, there was no change in the frequency of mEPSCs (right; *N*_CON_=11, *N*_miR-124_=10; two-tailed student’s *t*-test, *t*_19_=-0.787, *p*=0.441). **(F)** Downstream targets of miR-124 were predicted by identifying the overlapping genes from the following four types of prediction databases: 1) sequence-matched putative target database merging the datasets from TargetScan, DIANA, and miRcode; 2) Ago-miRNA-mRNA interaction database, for which the HIT-CLIP dataset was used; 3) experimentally-screened target database utilizing the miRTarBase dataset; 4) database consisting of the genes downregulated in the cerebral cortex of both SZ and BP patients, for which we used recently published microarray and RNA sequence datasets. **(G)** *Gria2* and *Adarb1*, but not *Gria1*, were found to be significantly downregulated in FACS-isolated miR-124-overexpressing excitatory neurons. *Gria2* (*N*_CON_=7, *N*_miR-124_=7; two-tailed student’s *t*-test, *t*_12_=3.678, *p*=0.003). *Adarb1* (*N*_CON_=7, *N*_miR-124_=7; two-tailed student’s *t*-test, *t*_12_=2.220, *p*=0.046). *Gria1* (*N*_CON_=7, *N*_miR-124_=7; two-tailed student’s *t*-test, *t*_12_=0.85, *p*=0.412). (**H**) Significant downregulation of *GRIA2* and decreasing trend of *ADARB1* were observed in olfactory neuronal cells from SZ and BP patients. *GRIA2* (*N*_CON_=39, *N*_SZ_=38, *N*_BP_=24; Welch’s ANOVA, *F*_2,64.642_=5.268, *p*=0.008; *post hoc* Games-Howell test). *ADARB1* (*N*_CON_=39, *N*_SZ_=38, *N*_BP_=25; one-way ANOVA, *F*_2,99_=2.188, *p*=0.118; *post hoc* Tukey’s HSD test). Bar graph expressed as mean ± SEM. ** *p* < 0.01, *** *p* < 0.001. More detailed statistical information is in **Table S2.**

### Overexpression of miR-124 increases GRIA2-lacking calcium-permeable AMPARs (CP-AMPARs)

Next, we investigated a potential molecular mechanism by which miR-124 upregulation alters AMPAR-mediated synaptic transmission that in turn drives the previously observed behavioral phenotypes. To address this, we looked for the relevant downstream targets of miR-124 and narrowed them downed to 12 genes by identifying the overlapping genes of the following four databases: (i) sequence matching-based putative target database; (ii) Ago-miRNA-mRNA interaction database; (iii) experimentally-screened target database; and (iv) database for the genes downregulated in the cerebral cortex from both patients with SZ and those with BP (**Figure 3F**). Among these 12 candidates, there were 2 molecules that could directly affect AMPAR-mediated synaptic transmission: *GRIA2* (AMPAR subunit 2) (Studniarczyk et al., 2013) and *ADARB1* (RNA editing enzyme acting on *GRIA2* pre-mRNA at the Q/R site) (Liu et al., 2004). We found that expression of both genes is downregulated upon miR-124 overexpression in mouse mPFC excitatory neurons (**Figure 3G**) and confirmed these molecular changes in human olfactory neuronal cells (**Figure 3H**). AMPARs are calcium-permeable if they lack the GRIA2 subunit or if they contain the unedited GRIA2(Q) subunit (Henley and Wilkinson, 2016). Thus, downregulation of either *Gria2* or *Adarb1* by miR-124 overexpression can increase the abundance of calcium-permeable AMPARs (CP-AMPARs) (Gascon et al., 2014; Hou et al., 2015), which can in turn increase the postsynaptic conductance of AMPARs and mEPSC amplitude. The modest downregulation of *Adarbl* induced by miR-124 overexpression did not affect RNA editing efficiency of *Gria2* (**Figure S6A**). However, the more robust downregulation of *Gria2* mRNA induced by miR-124 overexpression led to a significant reduction in surface GRIA2 protein expression (**Figure S6B**). This implies that downregulation of GRIA2, but not ADARB1, can lead to increase in CP-AMPARs.

### Blockage or reversal of GRIA2-lacking CP-AMPARs ameliorates the synaptic and behavioral deficits relevant to SZ and BP

We then addressed whether the increase in GRIA2-lacking CP-AMPARs induced by miR-124 overexpression causes increase in the amplitude of mEPSCs. Bath application of Naspm, a selective antagonist of CP-AMPARs, on miR-124-overexpressing brain slices normalized the increased amplitude of mEPSCs, proving the causality of the increase in CP-AMPARs on the increase in mEPSC amplitude (**Figure 4A**). As a result, we questioned whether the increase in CP-AMPARs directly causes the behavioral deficits in social interaction and psychostimulant sensitivity. We found that the behavioral deficits in social interaction and psychostimulant sensitivity were ameliorated by local infusion of Naspm, but not saline, into the mPFC, implying the causality of increased CP-AMPARs on the behavioral deficits (**Figures 4B and 4C**). We finally tested whether the restoration of GRIA2 alone could also ameliorate the behavioral deficits. We introduced AAV-mediated expression of GRIA2, along with miR-124, in mouse mPFC excitatory neurons. As a result, the mice with restored GRIA2 in the mPFC showed amelioration of their behavioral deficits, showing the causal effect of GRIA2 downregulation on the behavioral phenotypes (**Figures 4D and 4E**).

**Figure 4.**
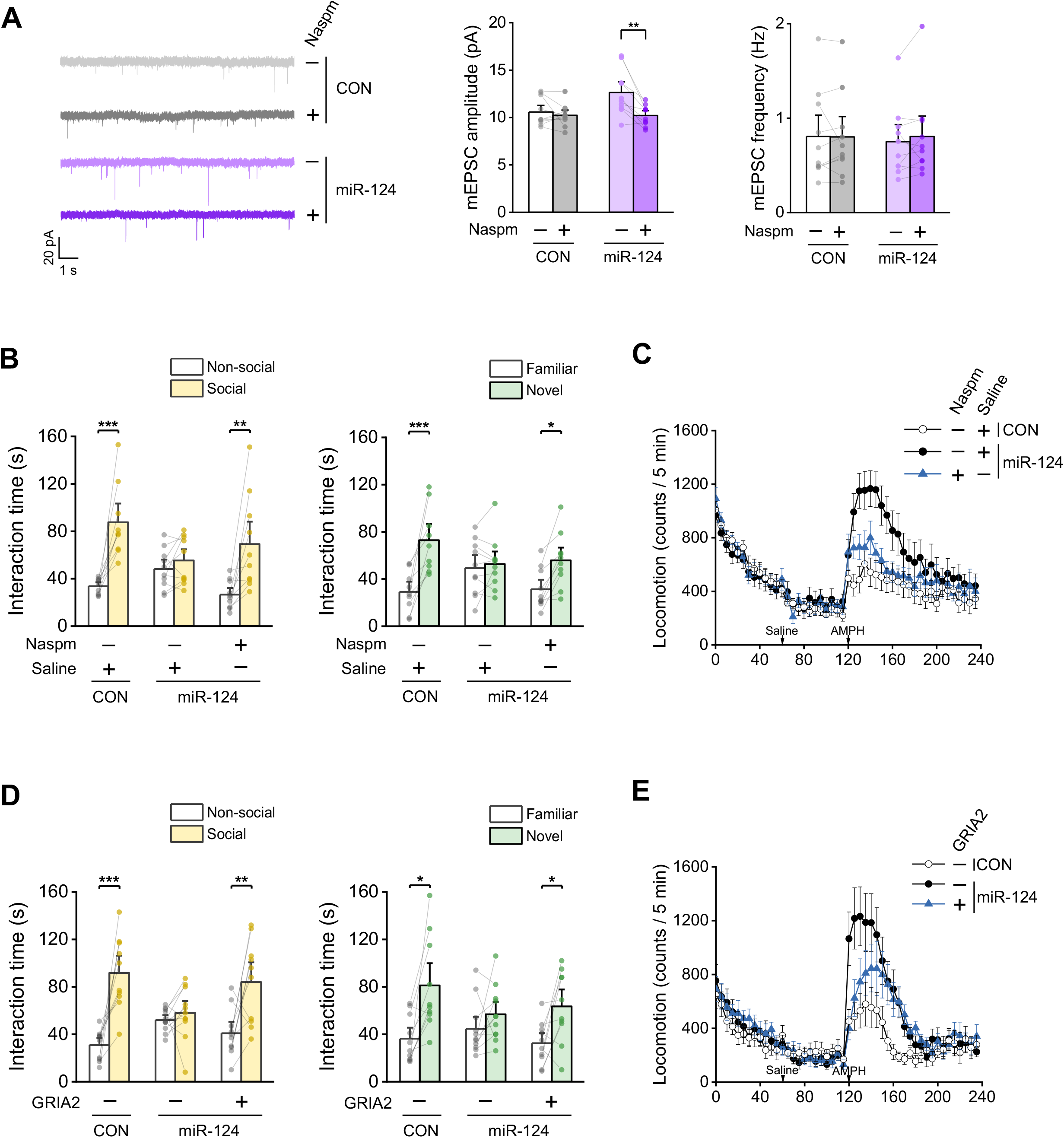
Blocking GRIA2-lacking CP-AMPARs by Naspm or direct restoration of GRIA2 expression ameliorates the previously observed synaptic and behavioral deficits. **(A)** The increase of mEPSC amplitude in mPFC excitatory neurons, induced by AAV-mediated miR-124 overexpression, was normalized by bath application of Naspm, a selective antagonist of CP-AMPARs. mEPSC amplitude (*N*_CON_=10, *N*_miR-124_=10; two-way mixed ANOVA, AAV x Naspm, *F*_1,18_=8.454, *p*=0.009; *post hoc* within-group comparisons using two-tailed paired Student’s *t*-test). mEPSC frequency (*N*_CON_=10, *N*_miR-124_=10; two-way mixed ANOVA, AAV x Naspm, *F*_1,18_=0.593, *p*=0.451; *post hoc* within-group comparisons using two-tailed paired Student’s *t*-test). **(B)** Behavioral deficits in sociability (left) and social novelty recognition (right), which were induced by miR-124 overexpression in mPFC excitatory neurons, were rescued by local infusion of Naspm into the mPFC. Sociability phase of trial 1(*N*_CON AAV + saline_=9, *N*_miR-124 AAV + Saline_=10, *N*_miR-124 AAV + Naspm_=10; miR-124 AAV + Saline vs. miR-124 AAV + Naspm, two-way mixed ANOVA, Intervention x Chamber, *F*_1,18_=6.725, *p*=0.018; *post hoc* within-group comparisons using twotailed paired Student’s *t*-test). Social novelty recognition phase of trial 2 (*N*_CON AAV + Saline_=9, *N*_miR-124 AAV + Saline_=10, *N*_miR-124 AAV + Naspm_=10; miR-124 AAV + Saline vs. miR-124 AAV +Naspm, two-way mixed ANOVA, Intervention x Chamber, *F*_1,18_=9.276, *p*=0.007; *post hoc* within-group comparisons using twotailed paired Student’s *t*-test). **(C)** Locomotor hypersensitivity to AMPH, induced by overexpression of miR-124 in mPFC excitatory neurons, was ameliorated by local infusion of Naspm into the mPFC. The ameliorating effect was most evident for the first 15 min after the *i.p.* injection of AMPH (*N*_CON AAV + Saline_=12, *N*_miR-124 AAV + Saline_=13, *N*_miR-124 AAV + Naspm_=10; miR-124 AAV + Saline vs. miR-124 AAV + NaSpm, two-way mixed ANOVA with the Greenhouse-Geisser correction, Intervention x Time, *F*_1.542,32.391_=9.671, *p*=0.001; *post hoc* two-tailed paired *t*-test for each time point). **(D)** Behavioral deficits in sociability (left) were rescued by AAV-mediated restoration of GRIA2. Behavioral deficits in social novelty recognition (right) were ameliorated, but not robustly rescued, by the restoration of GRIA2. Sociability phase of trial 1 (*N*_CON AAV + No-GRIA2 AAV_=10, *N*_miR-124 AAV + No-GRIA2 AAV_=11, *N*_miR-124 AAV + GRIA2 AAV_=10; miR-124 AAV + No-GRIA2 AAV vs. miR-124 AAV + GRIA2 AAV, two-way mixed ANOVA, Intervention x Chamber, *F*_1,19_=5.741, *p*=0.027; *post hoc* within-group comparisons using two-tailed paired Student’s *t*-test). Social novelty recognition phase of trial 2 (*N*_CON AAV + No-GRIA2 AAV_=10, *N*_miR-124 AAV + No-GRIA2 AAV_=11, *N*_miR-124 AAV + GRIA2 AAV_=10; miR-124 AAV + No-GRIA2 AAV vs. miR-124 AAV + GRIA2 AAV, two-way mixed ANOVA, Intervention x Chamber, *F*_1,19_=1.876, *p*=0.187; *post hoc* within-group comparisons using twotailed paired Student’s *t*-test). **(E)** Locomotor hypersensitivity to AMPH, induced by overexpression of miR-124 in mPFC excitatory neurons, was ameliorated by AAV-mediated restoration of GRIA2. The ameliorating effect was most evident for the first 60 min after the *i.p.* injection of AMPH (*N*_CON AAV + No-GRIA2 AAV_=11, *N*_miR-124 AAV + No-GRIA2 AAV_=10, *N*_miR-124 AAV + GRIA2 AAV_=11; miR-124 AAV + No-GRIA2 AAV vs. miR-124 AAV + GRIA2 AAV, two-way mixed ANOVA with the Greenhouse-Geisser correction, Intervention x Time, *F*_2.593,49.262_=6.301, *p*=0.002; *post hoc* two-tailed paired *t*-test for each time point). Bar graph expressed as mean ± SEM. \**p* < 0.05, \*\**p* < 0.01, \*\*\**p* < 0.001. More detailed statistical information is in **Table S2.**

## DISCUSSION

In the present study, we leveraged a novel research framework in which human and animal studies are combined to address a disease mechanism that bridges across pathogenesis, pathophysiology, and phenotypic/clinical manifestations. This framework enabled us to link together multiple layers from genetics, molecular and cellular mediators, and circuitry, to behavior in an integrative manner. Utilizing this research framework, we demonstrated that upregulation of miR-124, which is associated with the polygenic contribution shared between SZ and BP, increases GRIA2-lacking CP-AMPARs and perturbs AMPAR-mediated excitatory synaptic transmission, eventually leading to behavioral deficits commonly relevant to SZ and BP. Our research framework is also applicable to a wide range of other psychiatric conditions particularly when aiming to translate polygenic insights into clinical manifestations through mechanistically defined neurobiological pathways.

Examining miR-124 in olfactory neuronal cells, which are easily obtainable from living human subjects through a nasal biopsy, holds promise in that this molecule may be used as a biomarker for the shared pathophysiology of SZ and BP. Efforts to minimize the invasiveness of the biopsy have recently been made, including the method of obtaining neuronal cells via soft brush swab that can be applied to children (Lavoie et al., 2017). Given that the detection of pathological molecular signatures from early, pre-symptomatic stages is known to be important in brain disorders, like the case of mild cognitive impairment that precedes Alzheimer’s disease (Hampel et al., 2004; Jack et al., 2010), profiling miR-124 expression of olfactory neuronal cells from subjects at clinically and/or genetically high risk to SZ and BP in their pre-symptomatic stages will have prophylactic significance.

Although the present study supports the utility of olfactory neuronal cells as a surrogate to estimate disease-associated molecular changes in PFC neurons, we acknowledge that disease-associated changes of miR-124 are less prominent in the postmortem PFC than in olfactory neuronal cells from living patients with active symptoms. A major challenge of using postmortem brains in the study of psychotic disorders whose onsets are usually in late adolescence and young adulthood is that they may not fully capture molecular signatures in patients with full symptomatic manifestation (Lavoie et al., 2017; Namkung et al., 2018). In addition, disease-associated molecular signatures may be masked by effects of long-term medications and substance abuse as well as compensatory changes over the lifetime (Yang and Sawa, 2017). It is therefore possible that disease-associated changes of miR-124 may become diluted over the lifetime and eventually less prominent in the postmortem brain than in olfactory neuronal cells that were biopsied when the symptoms were active. Together, olfactory neuronal cells may enable not only to avoid a false negative pitfall arising from confounding factors in the postmortem brain, but also to discover novel molecular pathways captured in biopsied neuronal cells from patients with active psychiatric symptoms. Additionally, while we focused on studying the mouse mPFC based on human postmortem evidence, we cannot rule out the possibility that miR-124 is dysregulated in other brain regions of SZ and BP patients, which may underlie behavioral dimensions other than those observed in our mouse model. Finally, it should be noted that, although GRIA2 has been found to be a crucial downstream target of miR-124 that can mediate the observed synaptic and behavioral phenotypes in the present study, other targets of miR-124 may play crucial roles in different behavioral dimensions.

We also admit that the present study includes additional caveats. Firstly, while a strategy integrating PRS associations with mediation analysis enabled us to identify miR-124 as a potential pathophysiological mediator that connects pathogenic (genetic) and phenotypic feature(s) common to SZ and BP, it remains at the level of causal inference: this does not rigorously demonstrate the causality among these components, in particular because PRSs constructed from many SNPs in general have highly pleiotropic influences (Martin et al., 2019). We therefore sought to complement the causal relationship between miR-124 and clinical diagnoses with Mendelian randomization using several miR-124-associated SNPs as instruments, given that applying a relatively small number of instruments is still useful in drawing causal inference in many fields (Elliott et al., 2009; Haycock et al., 2017; Swerdlow et al., 2012; Swerdlow et al., 2015). However, further validation in a much larger cohort of subjects is awaited, in which more robust causality using a larger number of instruments can be tested. Nevertheless, our animal data sufficiently complement the human data by demonstrating the causal impacts of miR-124 on behavioral dimensions relevant to SZ and BP (Monteggia et al., 2018).

Despite these caveats, however, our preclinical insights can be integrated into large-scale multi-institutional clinical studies whereby behavioral constructs and brain function studied in our animal model can be revisited in the context of miR-124, for which neuropsychology and brain imaging may be key readouts (Drysdale et al., 2017; Krystal and State, 2014). Furthermore, our findings on the miR-124-AMPAR pathway may lead to the development of new drugs that target AMPARs at the level of pathophysiology, which may be more tractable than the level of pathogenesis, in particular for psychiatric conditions with complex genetic architectures and gene-environment interactions (Howe et al., 2018; Namkung et al., 2018). In the bigger picture, our new intellectual framework may open new avenues for the discovery of additional key biological pathways that provide mechanistic bridges between pathogenesis, pathophysiology, and clinical phenotypes for a wide range of neuropsychiatric conditions, hopefully invigorating drug development pipelines (Hall et al., 2015).

## ACKNOWLEDGMENTS

We thank Dr. Ingie Hong for insightful comments on the manuscript, and Dr. Kazuhiro Ishi for helpful advice on AAV generation. We also appreciate Dr. Benjamin Neale and Dr. Dongwon Lee for constructive comments on the genetic study. This work was supported by National Institutes of Mental Health Grants MH-092443 (to AS), MH-094268 (to AS), MH-105660 (to AS), and MH-107730 (to AS); foundation grants from Stanley, RUSK/S-R, and NARSAD/Brain and Behavior Research Foundation (to AS).

## AUTHOR CONTRIBUTIONS

H.N. and A.S. formulated the conceptual framework, designed experimental outlines, and wrote the manuscript. H.N. designed and conducted experiments. H.Y., D.F., B.J.L., S.L., S.K., A.S., K.S., H.J.P., and K.I. conducted experiments. P.Z. assisted genetic study. N.C. designed genetic analysis and helped conceptual framework formulation in genetic study. R.H. assisted conceptual framework formulation and experimental design in neurobiological study. A.S. supervised the overall project.

## DECLARATION OF INTERESTS

The authors declare no competing interests.

## Notes

### Competing Interest Statement

The authors have declared no competing interest.

