## Supplemental materials for "The miR-124-AMPAR pathway connects polygenic risks with behavioral changes shared between schizophrenia and bipolar disorder"

#### EXPERIMENTAL MODEL AND SUBJECT DETAILS

##### Human subjects

Patients with schizophrenia (SZ, n=38) and those with bipolar disorder (BP, n=25), and healthy controls (CON, n=39) were recruited at the Johns Hopkins Schizophrenia Center. All subjects had a clinical interview performed by board-certified psychiatrists with the Structured Clinical Interview for DSM-IV Axis I Disorders-Clinician Version (SCID-IV) or Diagnostic Interview for Genetic Studies (DIGS) to confirm a diagnosis of SZ or BP with or without a lifetime history of psychotic symptoms. Subjects who have history of traumatic brain injury, drug or alcohol abuse, excessive bleeding, or bleeding disorders were excluded. Current psychiatric symptom severity was rated by the Scales for the Assessment of Positive and Negative Symptoms (SAPS and SANS) for SZ, and that for BP was rated by the Montgomery-Asberg Depression Rating Scale (MADRS) and Young Manic Rating Scale (YMRS). The study was approved by the Johns Hopkins Institutional Review Board, and all subjects gave written consent for their participation.

##### Human olfactory neuronal cells

Olfactory epithelium (OE) tissues were obtained by nasal biopsy at the Johns Hopkins Otolaryngology Clinic (Kano et al., 2013). Local anesthesia to the nasal cavity was provided by lidocaine liquid 4% and oxymetazoline HCl 0.05% sprayed in the nose. It was followed by injection of 1% lidocaine with 1/100,000 epinephrine for both anesthesia and vasoconstriction. The biopsy procedure was performed under endoscopic control, and either a small curette or a biting forceps was used to remove tissue. To avoid trauma to the cribriform plate, the biopsies were usually taken from the upper nasal septum. Occasionally, a small piece of superior turbinate (which is usually lined with olfactory tissue) was removed from the lateral nasal wall. Six 5-mm$^3$

tissue pieces were removed from either the front or the back of the olfactory cleft, and sometimes both, from each nostril. After the biopsy, subjects were observed in the clinic for 15 to 30 min, but no packing or treatment was needed. Dissociated olfactory cells were prepared as follows (Kano et al., 2013). First, OE tissue pieces were incubated with 2.4 U/mL Dispase II for 45 min at 37$^{o}$C, and mechanically minced into small pieces. Then, the tissue pieces were further treated with 0.25 mg/mL collagenase A for 10 min at 37$^{o}$C. Cells were gently suspended, and centrifuged to obtain pellets. Cell pellets were resuspended in D- MEM/F12 supplemented with 10% FBS and antibiotics (D-MEM/F12 medium), and tissue debris was removed by transferring only the cell suspension into a new tube. Cells were then plated on 6-well plates in fresh D-MEM/F12 medium. Cells floating or loosely attached to the plate were collected on days 2 and 7, and further incubated on 6-well plates until they reached confluency. Finally, cells were collected by gentle trypsinization and stored in liquid nitrogen tank until further use. After recovery, cells were maintained in D-MEM/F12 medium and supplemented with fresh medium every 2-3 days.

**Human Lymphoblasts**

Lymphocytes were obtained from peripheral blood of patients with SZ (n=23) and CON (n=22) who underwent nasal biopsy, followed by infection with Epstein-Barr virus and incubation at 37.8 °C for 20-25 days to generate lymphoblasts at the Johns Hopkins University Genetic Resources Core Facility.

**Human Brains**

Human postmortem prefrontal cortices (Brodmann's area 45, BA45) from CON (n = 15), SZ (n = 15), BP (n = 15), and MDD (n=15) were obtained from the Stanley Foundation Brain Collection (Bethesda, MD), and utilized in our previous (Sawamura et al., 2005) and current studies. The specimens were collected by medical examiners. The gray matter of the prefrontal cortices (BA45) was dissected with a sharp blade for RNA extraction. The permission of the next of kin was obtained in all cases. The demographic, clinical, and storage characteristics have been already published (Torrey et al., 2000). Each diagnostic group was maximally matched according to the several parameters, including age at death, sex, post-mortem interval (PMI), brain pH, brain weight, and storage days. The expression of pri-miR-124-2 and pre-miR-124-2 in the dorsolateral prefrontal cortex (DLPFC, BAs 9 and 46) was analyzed from an independent sample of CON (N=225), SZ (N=143), and BP (N=50) (Jaffe et al., 2018). The details of tissue acquisition, handling, processing, dissection, clinical characterization, diagnoses, neuropathological examinations, and RNA extraction have been reported previously (Jaffe et al., 2018; Lipska et al., 2006).

**Animals**

Adult wild-type male C57BL/6J mice at 9-12 weeks old (Jackson Laboratory) were used for all experiments in this study. Mice were deeply anesthetized with intraperitoneal injection of Avertin (0.4 mg/g) and placed in a stereotaxic apparatus (David Kopf Instruments). The head was leveled using Bregma and Lambda reference points and a craniotomy was performed. The mice were injected bilaterally with 0.5 ul of AAVs into the medial prefrontal cortex (mPFC) using a syringe (Hamilton, 80383) with the coordinate from Bregma: +1.94 mm AP, ± 0.3 mm ML, -1.5 mm DV. After each injection, the syringe was left in place for 5 minutes and then slowly withdrawn. The skin was sutured and closed. Mice were recovered from anesthesia on a heating pad, and then were returned to their home cage. Animals were allowed about 3 weeks of recovery time after the surgery. All procedures of handling animals were in accordance with the guidelines from NIH, and with approval of the Johns Hopkins University Institutional Animal Care and Use Committee.

**METHOD DETAILS**

**Microarray experiments and data analysis**

Total RNA was extracted from olfactory neuronal cells derived from CON (n=6) and SZ (n=5) using RNeasy kit (Qiagen). The quality of RNA was 10 in RNA integrity number (RIN) score, which was assessed using a Bioanalyzer RNA 6000 Nano Chip (Agilent Technologies). Microarray experiments were performed at the Johns Hopkins University Microarray Core Facility. Biotin-labeled cRNA was produced using the Affymetrix 3' Amplification One-Cycle Target labeling kit according to the manufacturer's protocol. The biotin-labeled cRNA was fragmented and hybridized on Affymetrix U133 Plus 2.0 chip at 45$^{o}$C overnight. Post-hybridization washing was done as indicated in the Affymetrix instructions. The chips were stained with R-Phycoerythrin streptavidin (Invitrogen) and scanned for signal detection with the Affymetrix scanner. The Affymetrix Microarray Analysis Suite version 5 (MAS5) R package was used to process scanned chip images. Raw data were imported into a single AffyBatch object from CEL files. The array quality was assessed using the qc() function in the simpleaffy R package (Wilson and Miller, 2005) to check if any outlier chips exist. Expression values (signal intensities) and detection calls for each probe set were calculated. Each chip was globally scaled with a target value of 100 to make them comparable (MAS5-scaled). We eliminated unreliable signals by filtering out the probe sets whose detection calls were absent in all samples, and log2-transformed the cleaned data. A heatmap with hierarchical clustering was visualized based on z-scores in row direction using the heatmap.2() function in the gplots R package. The genes differentially expressed between CON and SZ were identified by two-tailed student's *t*-test. The differentially expressed genes ($p < 0.05$) were then subject to subsequent pathway enrichment analysis.

**Pathway enrichment analysis**

To identify pathways significantly enriched for the genes dysregulated in SZ olfactory neuronal cells, the differentially expressed genes detected by microarray experiments were applied to the Illumina NextBio Research platform (https://www.nextbio.com) that comprehensively integrates pre-compiled databases of pathways derived from Gene Ontology, TargetScan, InterPro, and Broad MSigDB (canonical pathways, regulatory motifs, and positional gene sets). Furthermore, to study potential dysregulation of miR-124 pathway in the SZ postmortem prefrontal cortex (PFC), the recent CommonMind Consortium (CMC) RNA sequence data of SZ postmortem dorsolateral prefrontal cortex (DLPFC, BAs 9 and 46) was investigated (Fromer et al., 2016). A differential gene expression dataset adjusted for known and hidden variables detected by surrogate variable analysis (SVA) was accessed from the CMC data release 1. Then, the differentially expressed genes identified in the dataset were applied to the NextBio Research platform with the same procedure as that used for the microarray pathway analysis.

**Quantitative real-time polymerase chain reaction (qRT-PCR) for human samples**

Total RNA was extracted with RNeasy Mini kit (Qiagen) following the manufacturer's instruction, and further purified by RNase-Free DNase kit (Qiagen). RNA integrity was evaluated by spectroscopic analysis using the Nanodrop 2000 (Thermo Scientific). Equivalent amounts of RNA (0.2 µg/sample) were used for reverse transcription PCR with miScript II RT Kit (Qiagen) according to the manufacturer's instructions. Mature miRNAs were detected in a QuantStudio 12K Flex Real-Time PCR System (Applied Biosystems), using miScript SYBR Green PCR kit (Qiagen) and the following miScript Primer Assays (Qiagen): hsa-miR-124-3p (MS00006622), hsa-miR-17-5p (MS00029274), hsa-miR-107 (MS00031255), hsa-miR-132-3p (MS00003458), hsa-miR-134-5p (MS00031437), hsa-miR-137 (MS00003486), hsa-miR-652-3p (MS00010451), and RNU6-2 (MS00033740). Primary miRNAs were detected using miScript SYBR Green PCR kit (Qiagen) and the following miScript Precursor Assays (Qiagen): hsa-mir-124-1 (MP00000371), hsa-mir-124-2 (MP00000378), and hsa-mir-124-3 (MP00000392). RNU6-2 served as a reference gene for normalization. Relative expression was analyzed using the $2^{-\Delta\Delta C_t}$ method (Livak and Schmittgen, 2001).

**RNA sequencing of postmortem DLPFC samples (LIBD sample)**

The expression of our target genes was investigated in RNA sequencing data of the postmortem DLPFC from CON, SZ, and BP, which were generated at the Lieber Institute for Brain Development (LIBD). In particular, BrainSeq PhaseI poly(A)+ library data were analyzed (BrainSeq and BrainSeq, 2015; Jaffe et al., 2018). The detailes of RNA extraction and sequencing, and genotyping have been reported previously (BrainSeq and BrainSeq, 2015; Jaffe et al., 2018). In the present work, the data were revisited for aligning reads to the human genome UCSC hg38 build (Eagles et al., 2021). We examined the relationship between diagnosis and gene expression, adjusting for age, sex, race, RNA integrity number (RIN), surrogate variables (SVs), and genetic principal components (top 10 PCs).

**Quality control, imputation, and polygenic risk scoring**

DNA collected from CON (n=45), SZ (n=38), and BP (n=25) subjects was genotyped on Illumina's Psych or global screening arrays at the Broad Institute (Cambridge, MA). Quality control (QC) was performed, according to the standards established by the Psychiatric Genomics Consortium (PGC) (Ruderfer et al., 2018). The quality control parameters for retaining SNPs and subjects were: SNP missingness < 0.05 (before sample removal); subject missingness ($p < 0.02$); autosomal heterozygosity deviation ($|F_{het}|<0.2$); SNP missingness < 0.02 (after sample removal); difference in SNP missingness between cases and controls < 0.02; and SNP Hardy-Weinberg equilibrium ($p > 10^{-6}$ in controls or $p > 10^{-10}$ in cases). Genotype imputation was conducted using the Michigan Imputation Server v1.0.4 (Das et al., 2016) with the 1000 Genomes phase 3 reference panel (Altshuler et al., 2015). The study samples were phased using Eagle (v2.3) (Loh et al., 2016) followed by imputation using minimac3 (v.2.0.1) (Howie et al., 2012). The imputed markers underwent a second stage of QC to exclude SNPs that were missing in >1% of individuals, had imputation information score (INFO) < 0.8, or had minor allele frequency (MAF) <0.05. After repeating the above QC and imputation procedures for both Psych and global screening arrays, we merged our study samples using the only overlapping markers. Using recent GWAS summary statistics (Ruderfer et al., 2018), we calculated genome-wide polygenic risk scores (PRSs) for SZ+BP, SZ, BP, SZ vs. BP in our independent samples of European ancestry as the weighted sum of risk alleles they carried. Using a PRS software (PRSice) (Euesden et al., 2015), a set of SNPs was filtered from a discovery dataset by LD pruning with a cutoff of $r^2 = 0.02$ within a 50-kb window, and PRSs of each individual were calculated for 10 *p*-value thresholds ($p_{T=}$0.0001, 0.001, 0.01, 0.05, 0.1, 0.2, 0.3, 0.4, 0.5, and 1.0, respectively). Population stratification was corrected with principal component analysis (PCA): the top 20 principal components (PCs) were calculated, and a subset of the PCs used to control for population stratification was selected by evaluating the proportion of variance explained by each PC. A logistic regression analysis was used to investigate the association between the PRSs and case-control status (either SZ or BP vs. CON) in our independent samples. Variance explained by the PRSs was calculated as the Nagelkerke's pseudo-$R^2$ of the full model including the PRSs and the top 7 PCs minus the Nagelkerke's pseudo-$R^2$ of the model including the only 7 PCs. The association between the PRSs and miR-124 expression was examined by using linear regression.

**Mediation analysis**

To quantify the extent to which miR-124 pathway can mediate the effects of PRSs (for SZ+BP, SZ, BP, and SZ vs. BP, respectively) on the diagnosis of either SZ or BP, we performed model-based mediation analyses using the mediation package in R (Tingley et al., 2014). The analysis proceeded with two models: an outcome model and a mediator model. We fitted the outcome model in which case-control status (outcome) was modeled as a function of PRS (treatment), miR-124 expression (mediator), and age, sex, PC (covariates). We also fitted the mediator model where miR-124 expression (mediator) was modeled as a function of PRS (treatment) and age, sex, PC, diagnosis (covariates). For a case-control study design in which sampling is associated with the binary outcome, the regression model for the mediator needs to account for the sampling design, although the estimates from the regression model for the outcome can be used in the analysis (VanderWeele, 2016). Therefore, in constructing the mediator model, diagnosis was also included as a covariate to account for the sampling design. The constructed outcome and mediator models were fitted separately, and then their fitted objects comprised the main inputs to the mediate function in the R mediation package. The package evaluated the total effect of PRSs on case-control status and partitioned this effect to the direct effect (not through the miR-124 pathway) and the mediated effect (through the miR-124 pathway). A bootstrapping method with 5,000 iterations was used to test the 95% confidence intervals of the mediated effects. To represent effect sizes for mediation analysis, proportion mediated was presented.

**Mendelian randomization**

We applied Mendelian randomization (MR) to explore the potential causal effect of an exposure (miR-124) on an outcome (SZ or BP), for which SNPs (significantly associated with miR-124) were used as instruments. To identify such SNPs significantly associated with miR-124 expression in olfactory neuronal cells, expression quantitative trait loci (eQTL) analysis was conducted for mature and primary miR-124s (miR-124 and pri-miR-124-1, -3; pri-miR-124-2 below detection limit), using the 88 samples across the 6.1 million genotyped and imputed markers with imputation score (INFO) > 0.8 and estimated minor allele frequency (MAF) > 0.05. eQTL were mapped using the following linear model constructed in MatrixEQTL (Shabalin, 2012): gene expression (quantile normalized) ~ SNP + age + sex + diagnosis + self-reported race + the top 3 PCs for each ethnic group. The false discovery rate (FDR) was estimated separately for cis-eQTL (defined as <1 MB between SNP marker and gene position) and trans-eQTL (>1 MB between marker and gene), with FDR < 5% used as the threshold for significant eQTL. Then, the SNP-exposure association (obtained from the eQTL analysis) was integrated with the SNP-outcome association [obtained from independent GWAS results (Ruderfer et al., 2018)] using a strategy known as 2-sample MR (2SMR) in the MR-Base platform (Hemani et al., 2018). To avoid bias in the MR estimates due to linkage disequilibrium (LD), LD clumping was applied using SNPsnap with the $R^2$ cutoff of 0.8 (Pers et al., 2015). The independent SNPs, which are available in both the SNP-exposure and SNP-outcome associations, were harmonized and then used as instruments to estimate the causal influence of the exposure on the outcome.

**Plasmid construction**

A miR-124-expressing plasmid was constructed with BLOCK-iT Pol II miR RNAi Expression Vector Kits (Invitrogen) following the manufacturer's instructions. The genomic sequence containing pre-miR-124 was synthesized and cloned to pcDNA6.2-GW/EmGFP-miR vector. The pcDNA6.2-GW/EmGFP-miR-control (EmGFP-miR-control) plasmid, which contains an insert not to target any known vertebrate genes, was used as the control vector. The EmGFP-miR-124 and EmGFP-miR-control were PCR amplified from pcDNA6.2-GW/EmGFP-miR-124 and pcDNA6.2-GW/EmGFP-miR-control vectors, respectively. Then, the EcoRI/HindIII-digested PCR products were cloned into the complementary EcoRI/HindIII-digested linearized backbone vector pAAV-6P-SEWB (Addgene, #31379) to generate pAAV-hSyn::EmGFP-miR-124 and pAAV-hSyn::EmGFP-miR-control. To generate pAAV-CaMKIIα::EmGFP-miR-124 and pAAV- CaMKIIα::EmGFP-miR-control, the KpnI/HindIII-digested EmGFP-miR-124 and EmGFP-miR-control PCR products were cloned into the complementary KpnI/HindIII-digested linearized backbone vector pAAV-CaMKIIα::hChR2(H134R)-EYFP (Addgene, #26969). To generate pAAV-CaMKIIα(0.4)::mCherry-GRIA2, the mCherry and GRIA2 were PCR amplified from pAAV-CaMKIIa(0.4)-mCherry-T2A-CreERT2 (Vector Biolabs, #VB1426) and tdTomato-GRIA2/pcDNA3.1 (Gu et al., 2016), respectively, and cloned into the backbone vector pAAV-CaMKIIa(0.4)-EGFP-WPRE (Addgene, #105541). To generate pAAV-CaMKIIα(0.4)::mCherry, the mCherry was PCR amplified from pAAV-CaMKIIa(0.4)-mCherry-T2A-CreERT2 (Vector Biolabs, #VB1426), and cloned into the same backbone vector pAAV-CaMKIIa(0.4)-EGFP-WPRE (Addgene, #105541).

**AAV production**

Production of adeno-associated viruses (AAVs) was performed as described previously with some modifications (Choi et al., 2007). Human embryonic kidney (HEK) 293-AAV cells (Stratagene) were co-transfected with AAV vectors and helper vectors (pAd helper vector and pAAV2/1 packaging vector) using the standard calcium phosphate precipitation method. Cells were harvested 72 hours post-transfection and disrupted by three times of freeze and thaw cycles. AAV viral particles were isolated and concentrated by a series of ammonium sulfate precipitations, applied to a discontinuous gradient of iodixanol (Optiprep™ density gradient medium, Sigma-Aldrich), and centrifuged at 60,000 rpm for 1.5 hours, 18 °C. An AAV-containing 40 % iodixanol fraction was carefully collected from an ultracentrifuge tube (Beckman), and concentrated using a centrifugal filter device (Amicon, Ultracel-100K). Stock viral titers were determined by qRT-PCR using the following AAV2 ITR primers: forward (5′-GGAACCCCTAGTGATGGAGTT-3′) and reverse (5′-CGGCCTCAGTGAGCGA-3′) primers. The AAVs whose titers were at least higher than $1 \times 10^{12}$ genomic copies (GC)/mL were utilized for experiments, although most of the AAVs had titers higher than $1 \times 10^{13}$ GC/mL.

**Fluorescence-Activated Cell Sorting (FACS)**

The mice injected with AAV-CaMKIIa::EmGFP-miR-124 or AAV-CaMKIIa::EmGFP-miR-control into the mPFC were subject to whole brain extraction. The mPFC tissues including EmGFP-positive neurons were carefully dissected using a stereomicroscope with a fluorescent flashlight (Nikon, #SMZ 745T). Cells were dissociated by utilizing papain dissociation kit (Worthington, LK003150) with a protocol optimized for isolating living neurons from the adult mouse brain (Saxena et al., 2012). Using the FACSAria cell sorter (BD Biosciences) at Johns Hopkins Bayview Immunomics Core, 50,000 EmGFP-positive neurons from each mouse were sorted and collected directly into QIAZol Lysis reagents in miRNeasy Mini kit (Qiagen) for the subsequent qRT-PCR.

**qRT-PCR for animal samples**

Total RNA was extracted with miRNeasy Mini kit (Qiagen) following the manufacturer's instructions. RNA integrity was evaluated by spectroscopic analysis using the Nanodrop 2000 (Thermo Scientific). Equivalent amounts of RNA (0.2 μg/sample) were used for reverse transcription PCR with miScript II RT Kit (Qiagen) according to the manufacturer's instructions. To profile miR-124 expression, qRT-PCR was performed in a QuantStudio 12K Flex Real-Time PCR System (Applied Biosystems) using miScript SYBR Green PCR kit (Qiagen) and the following miScript Primer Assays (Qiagen): Mm_miR-124 (MS00029211) and RNU6-2 (MS00033740). RNU6-2 served as a reference gene for normalization. To profile mRNA expression levels of *Gria1*, *Gria2*, and *Adarb1*, the following QuantiTect Primer Assays were used: Mm-Gria1 (QT01062544), Mm-Gria2 (QT00140000), Mm-Adarb1 (QT00164353), and Mm-Actb1 (QT00095242). *Actb* served as a reference gene for normalization. Relative expression was analyzed using the $2^{-\Delta\Delta C_t}$ method (Livak and Schmittgen, 2001).

#### Animal behavioral assays

##### *Social interaction*

The three-chamber social interaction test was performed to assess sociability (trial 1) and social novelty recognition (trial 2) at the Behavioral Core in the Johns Hopkins University School of Medicine. The testing arena consisted of three adjacent chambers separated by two clear plastic dividers and connected by two open doorways. Mice were habituated to the testing room and arena for 3 consecutive days (10 min free exploration of testing arena per day) before the testing day. Mice were transferred to a testing room and stayed for 1 hour before testing. The test consisted of 10 min habituation and two 10 min sessions with 10 min of inter-trial-interval. A subject mouse was kept in the center chamber during the 10 min habituation period. In the first 10 min session (trial 1, sociability session), the subject mouse freely investigated the three chambers, the novel object mouse (stranger 1) was placed under an inverted metal pencil cup in one of the side chambers, and another identical inverted empty cup was placed in the other side chamber. After 10 min of the trial 1 session, doors of dividers were closed and the subject mouse was kept in the center chamber during 10 min of the inter-trial-interval. In the second 10 min session (trial 2, social novelty recognition session), a novel object mouse (stranger 2) was put in the empty pencil cup, and the subject mouse freely investigated the three chambers. The subject mouse was restrained in the center chamber during the introduction of an object mouse. Age-matched C57BL/6J male mice were used as object mice in this study and housed under the same conditions as subject mice. Time spent sniffing each cup was scored by two independent investigators in a blind manner, and the scores were validated by comparing preference ratios measured by the two observers: for the trial 1, time spent for the chamber with a mouse / the chamber without a mouse; for the trial 2, time spent for the chamber with a novel mouse / the chamber with a familiar mouse.

*Amphetamine challenge*

To measure spontaneous activity, mice were placed in a transparent acrylic cage, and locomotion was measured every 5 min for 240 min by using digital counters with infrared sensors (Merquest, Scanet SV-10) at the Behavioral Core of the Johns Hopkins University School of Medicine. To measure saline and amphetamine-induced locomotion, mice received intraperitoneal injections of saline (1.5 mg/kg) and amphetamine (1.5 mg/kg) 60 min and 120 min after the start of habituation, respectively. In the pilot experiments alone (Extended Data Fig. 2), mice received subcutaneous injections of saline (1mg/kg) and amphetamine (1mg/kg) 60 min and 120 min after the start of habituation, respectively.

*Pre-pulse inhibition*

Mice were subjected to startle trials consisting of three conditions: (i) a 120 dB noise burst presented alone; (ii) a pre-pulse (74, 78, 82, 86, or 90 dB) followed by 120 dB noise burst or (iii) no stimulus (background noise alone), which was used to measure baseline movement in the startle chamber (San Diego Instruments, SR-Laboratory Systems). Percent prepulse inhibition of a startle response was calculated as: [1 – (startle response on prepulse + startle pulse) / (startle response on startle pulse)] X 100 (%).

*Forced swim test*

Mice were placed in a transparent glass cylinder (8 cm in diameter × 20 cm high), containing water at 22–23°C to a depth of 15 cm, and forced to swim for 6 min. Immobility was calculated as follows: immobility (%) = 100 x [360 s - swimming time (s)] / 360 s

**Immunohistochemistry**

Mice were euthanized and transcardially perfused with PBS, immediately followed by perfusion with 4% paraformaldehyde (PFA). Brains were post-fixed overnight in PFA at 4°C and coronally sectioned with a cryostat (Leica) at 40 µm. Brain cryosections were incubated for 16 h at 4°C with primary antibody: anti-GFP (1:1000; Abcam, ab13970) and anti-CaMKIIa (1:500; R&D systems, MAB5584) antibodies in 10 % normal goat serum (NGS) in PBST (PBS, 1% BSA, and 0.5% Triton X-100). The sections were then washed and incubated for 2 hours at room temperature with the secondary Alexa 488 goat anti-chicken (1:400; Invitrogen, A11039) and 568 goat anti-mouse (1:400, Invitrogen, A11031) antibodies in 1% NGS/PBST. Stained sections were washed and DAPI-stained (Invitrogen). All sections were mounted and imaged on either Zeiss AX10 Observer D1 or a Zeiss LSM800 confocal microscope at Johns Hopkins University Microscope Facility.

**Electrophysiology**

*Brain slice preparation for electrophysiological recordings*

After anesthetizing the mice with isoflurane, the brains were rapidly dissected and acute brain slices (300 µm) containing mPFC were prepared in ice-cold sucrose solution containing the following (in mM): 76 NaCl, 25 $NaHCO_3$, 25 D-(+)-glucose, 75 sucrose, 2.5 KCl, 1.25 $NaH_2PO_4$-$H_2O$, 0.5 $CaCl_2$-$2H_2O$, 7 $MgSO_4$-$7H_2O$, 2 pyruvic acid, 4 L-lactic acid, pH7.3, 310 mOsm. Coronal brain slices prepared using a vibratome (Leica VT 1200S) were recovered in warm (32–35°C) sucrose solution for 30 min and allowed to cool to room temperature in pre-warmed artificial CSF (aCSF) composed of (in mM): 125 NaCl, 26 $NaHCO_3$, 2.5 KCl, 1.25 $NaH_2PO_4$, 1 $MgSO_4$-7$H_2O$, 20 D-(+)-glucose, 2 $CaCl_2$-2$H_2O$, 0.4 ascorbic acid, 2 pyruvic acid, 4 L-lactic acid, pH 7.3, 315 mOsm. All solutions were continuously bubbled with 95% $O_2$/5% $CO_2$.

*Miniature excitatory postsynaptic current (mEPSC) recordings*

A single slice was transferred to the recording chamber and superfused with aCSF (3 ml/min) using a peristaltic pump (WPI). $GFP^+$ pyramidal neurons, which express AAV-CaMKIIa::EmGFP-miR124, in layer V of mPFC region were identified by their expression of GFP and typical morphology. The brain slice was visualized under an EMCCD camera (iXon3, ANDOR) using either transmitted light with infrared differential interference contrast optics or epifluorescence. Whole-cell patch-clamp recordings were made with a microglass electrodes (2.5–4 MΩ) filled with internal solution containing the following (in mM): 10 CsCl, 130 $CsMeSO_4$, 10 HEPES, 5 EGTA, 4 MgATP, 0.4 NaGTP, 10 phosphocreatine (Na), pH 7.3, 290 mOsm. mEPSCs were recorded for 5 min at $V_m$ = -70 mV in aCSF containing tetrodotoxin (1 μM) and picrotoxin (100 μM). Recordings were made using a Multiclamp 700B amplifier (Molecular Devices), and digitized by DigiDATA 1440A (Molecular Devices). Access resistance was monitored throughout the recording and a change less than 20% was deemed acceptable. All signals were low-pass filtered at 2 kHz and sampled at 10 kHz. The amplitude and frequency of mEPSCs were analyzed off-line with Mini Analysis software (Synaptosoft v6.0).

*mEPSC recordings with acute bath application of Naspm*

To assess the effect of Naspm, mPFC layer V pyramidal neurons expressing AAV-CaMKIIa::EmGFP-miR124 were patched and mEPSCs were recorded for 5 minutes at baseline following the above protocol. The recording chamber was then bath perfused with aCSF containing tetrodoxin (1 μM), picrotoxin (100 μM), and Naspm hydrochloride (100 μM). 5 minutes after the introduction of Naspm, mEPSCs were again recorded for 5 minutes. Each slice was recorded following only a single bath application of Naspm.

**MiR-124 target prediction analysis**

Downstream targets of miR-124 were predicted using four different types of miRNA target prediction databases: 1) sequence matching-based putative target database, for which we surveyed data from TargetScan (Agarwal et al., 2015), DIANA (Paraskevopoulou et al., 2013), and miRcode (Jeggari et al., 2012); 2) Ago-miR-124/Ago-mRNA interaction database, for which we surveyed data from Ago HITS-CLIP (Chi et al., 2009); 3) experimentally-validated target database, for which we used data from miRTarBase (Hsu et al., 2011); 4) Database for the genes downregulated in the cerebral cortex of both SZ and BP, for which we used recently published microarray and RNA sequence data (Gandal et al., 2018). We only considered predictive targets commonly implicated by these four types of datasets. We then further narrowed results to include only target genes that may directly affect AMPAR-mediated synaptic transmission.

**Surface labeling of synaptic AMPARs**

Cortical neurons were prepared from E 18 Sprague-Dawley rats. Neurons were dissociated and plated onto poly-L-lysine–coated coverslips in Neurobasal media (Invitrogen) supplemented with 2% B-27, 2 mM GlutaMax, 50 U/mL penicillin, 50 mg/mL streptomycin. Cells were infected with AAV at DIV4, and maintained in a humidified tissue culture incubator at 37 °C in a 95% air and 5% $CO_2$ mixture. On DIV 4 media was supplemented with 5 mM FDU (5-fluoro-2′-deoxyuridine; Sigma). Neurons were then maintained in glia-conditioned NM1 (Neurobasal media with 2% B-27, 2 mM GlutaMax, 50 U/mL penicillin, 50 mg/mL streptomycin, and 1% horse serum). For surface labeling, neurons were briefly rinsed in ACSF and were incubated with primary antibodies at 1:200 dilution at 10 °C for 20 min. Neurons were rinsed two times with ACSF and fixed with 4% PFA in PBS. After fixation neurons were again washed 2x with PBS followed by permeabilization with 0.2% Triton X100 for 10 min. After blocking with 3% BSA, neurons were incubated with secondary antibodies conjugated with Alexa dye for an hour. Cells were washed three times with PBS and mounted. Neurons were imaged with 63X objective at fixed exposure parameters for all experiments.

##### *Gria2* Q/R RNA editing assay

The RNA editing efficiency at the *Gria2* Q/R site was examined as previously described (Cueva Vargas et al., 2015). Briefly, total RNA extracted from FACS-sorted EmGFP-positive neurons was used for examining *Gria2* Q/R RNA editing. First-strand cDNA synthesis was performed using SuperScript III First-Strand Synthesis System (Life Technologies) and oligo $(dT)_{20}$ primers. To assess Q/R site editing, the *Gria2* open reading frame containing the Q/R site was amplified using the following primers: forward (5’- GAATGGTATGGTTGGAGAGC -3’) and reverse (5’-CACTT TCGATGGGAGACAC -3’) primers. The purified PCR products were then sequenced at the Johns Hopkins synthesis and sequencing facility.

##### Pharmacological and molecular intervention

For pharmacological intervention, following injection with either AAV-CaMKIIα::EmGFP-miR-124 or AAV- CaMKIIα::EmGFP-miR-control**,** a mouse was stereotaxically implanted with a 26 gauge guide cannula (PlasticsOne, C235G-1.0/SP) into the mPFC with the following coordinate from the Bregma: +1.94 mm AP, ± 0.5mm ML, -1.5 mm DV. Cannulas were held in place by dental acrylic. The intracranial microinjection was conducted using a microsyringe pump controller MICRO-4 (World Precision Instruments) connected with a Hamilton microsyringe (Plastic Ones, HSYR-1) through a tube (Plastics Ones, C230C/SPC) with the flow rate 300 nL/min. Saline or Naspm (1ug/0.3ul) was delivered bilaterally into the mPFC (0.3 μl/side) 30 mins before a three-chamber social interaction test or amphetamine injection in an amphetamine challenge test. For molecular intervention, AAV-CaMKIIα::EmGFP-miR-124 or AAV-CaMKIIα::EmGFP-miR-control was mixed in the 1:1 concentration ratio with AAV-CaMKIIα(0.4)::mCherry-GRIA2 or AAV-CaMKIIα(0.4)::mCherry. Then, the mixed AAVs (0.5 μL) were bilaterally injected into the mPFC using a syringe (Hamilton, 80383) with the following coordinate from the Bregma: +1.94 mm AP, ± 0.3mm ML, -1.5 mm DV.

#### QUANTIFICATION AND STATISTICAL ANALYSIS

Detailed statistical information on data for each figure is summarized in **Table S2**, in addition to the concise description in each figure legend. Briefly, no statistical methods were used to predetermine sample sizes, but most of our sample sizes are similar to those generally employed in the field. When performing parametric tests such as ANOVAs and *t*-tests, data distribution was assumed to be normal because such parametric tests are known to be robust against the normality assumption. When applying analysis of variance (ANOVA) with one factor, one-way ANOVA followed by *post hoc* Tukey's HSD test or Welch's ANOVA followed by *post hoc*

Games-Howell test was applied, depending on homogeneity of variance that was assessed by Levene's test. When applying ANOVA of two factors with one within-subjects factor and the other between-subjects factor, two-way mixed ANOVA was applied with the Greenhouse-Geisser correction for the violation of the sphericity assumption. Post-hoc two-tailed paired Student's *t*-test was used for within-group comparisons. For a comparison of independent two groups, two-tailed Student's or Welch's *t*-test was used, depending on homogeneity of variance assessed by Levene's test. To compare datasets with a small number of samples, non-parametric tests were applied: Kruskal-Wallis *H* test followed by *post hoc* with Dunn's test was used as the nonparametric alternative to one-way ANOVA.

### Figure S1

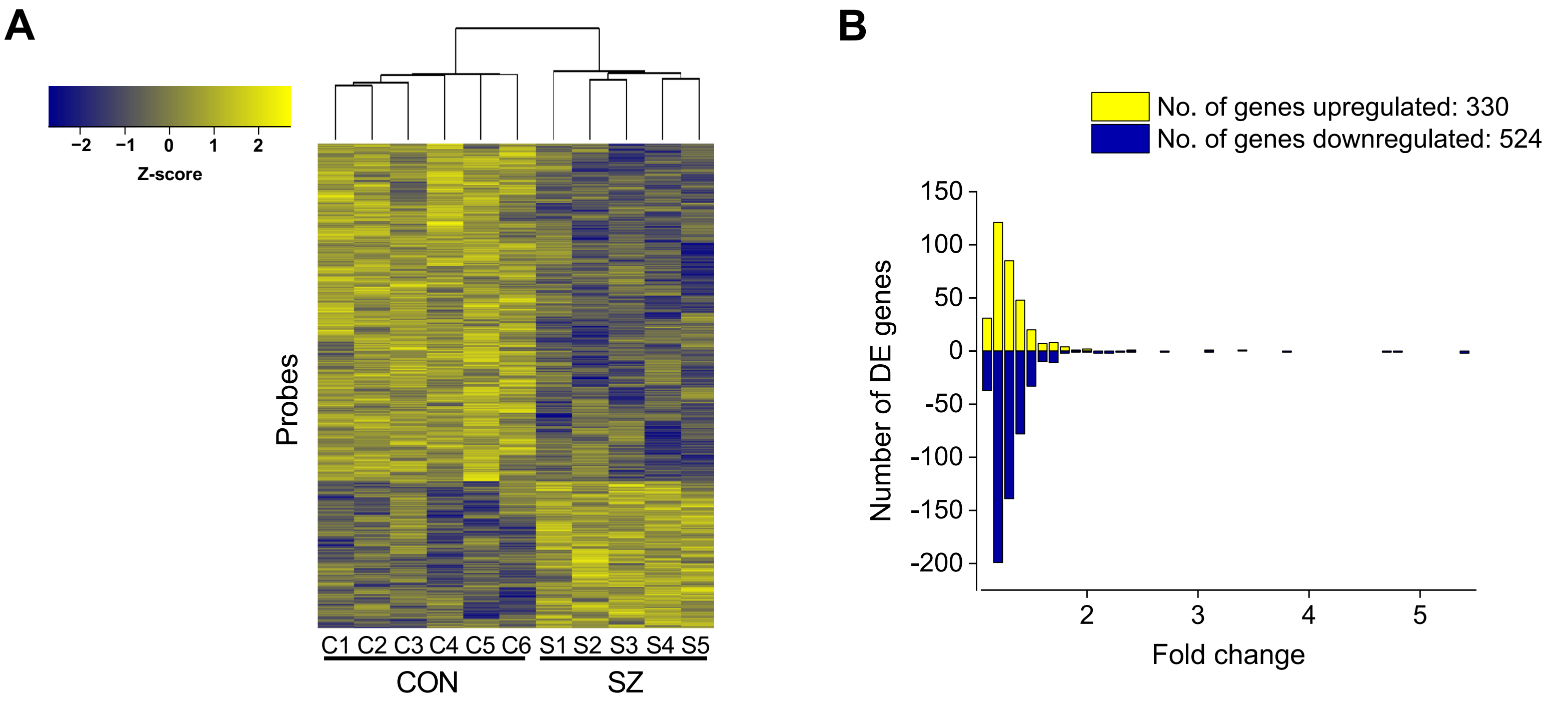


**Figure S1. Gene expression profiling of olfactory epithelium (OE)-derived neuronal cells (olfactory neuronal cells) identified differentially expressed genes in SZ compared with healthy CON. (A)** Microarray analysis of olfactory neuronal cells identified 854 dysregulated genes in SZ. For those genes, bivariate clustering of individuals (columns) on a heatmap depicts the control (CON) vs. schizophrenia (SZ) differences ($N_{CON}$=6, $N_{SZ}$=5). **(B)** Distribution of fold-change in differential expression. SZ:CON fold-changes for upregulated genes are plotted in yellow, and those for downregulated genes in blue.

### Figure S2

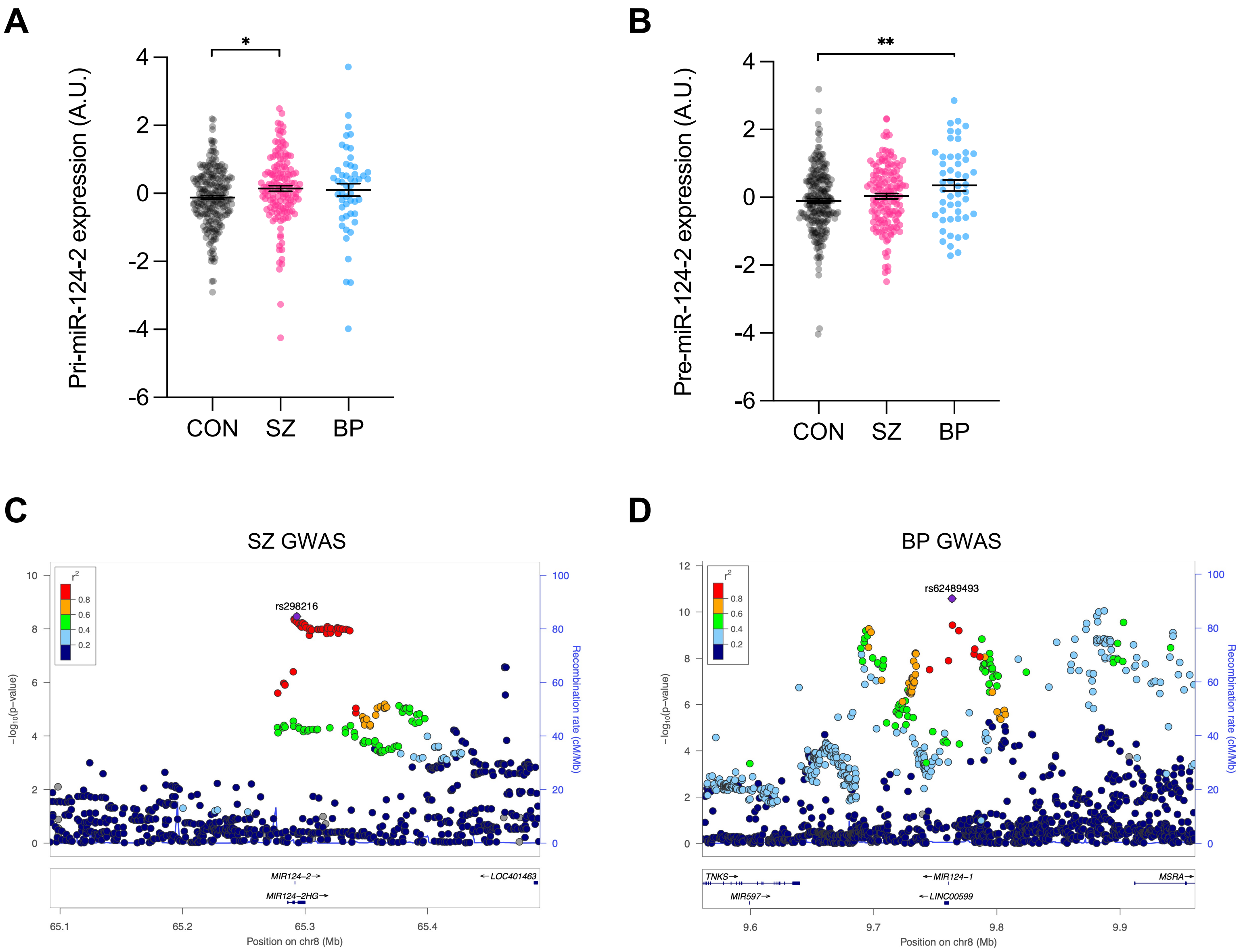

A
B
C
D
*
**
Pri-miR-124-2 expression (A.U.)
Pre-miR-124-2 expression (A.U.)
CON
SZ
BP
SZ GWAS
BP GWAS
rs298216
rs62489493
$r^2$
0.8
0.6
0.4
0.2
$-\log_{10}$(p-value)
Recombination rate (cM/Mb)
MIR124−2
MIR124−2HG
LOC401463
TNKS
MIR597
MIR124−1
LINC00599
MSRA
Position on chr8 (Mb)

**Figure S2. Upregulation of the primary miR-124-2 (pri-miR-124-2) and precursor miR-124-2 (pre-miR-124-2) in the postmortem DLPFC from SZ/BP, and SZ-/BP-associated SNPs near or within the pri-/pre-miR-124-2 and pri-/pre-miR-124-1. (A)** pri-miR-124-2 was found to be significantly upregulated in the postmortem DLPFC from SZ ($N_{CON}$=225, $N_{SZ}$=143, $N_{BP}$=50; one-way ANOVA, $F_{2,415}$=3.552, $p$=0.030; *post hoc* Tukey's HSD test). **(B)** pre-miR-124-2 was found to be significantly upregulated in the postmortem DLPFC from BP ($N_{CON}$=225, $N_{SZ}$=143, $N_{BP}$=50; one-way ANOVA, $F_{2,415}$=4.580, $p$=0.011; *post hoc* Tukey's HSD test). **(C)** Single nucleotide polymorphisms (SNPs) near or within the pri-/pre-miR-124-2 (MIR124-2HG/MIR124-2) gene loci have been shown to confer risk for SZ (Consortium et al., 2020). **(D)** SNPs near or within the pri-/pre-miR-124-1 (LINC00599/MIR124-1) gene loci have been shown to confer risk for BP (Mullins et al., 2021). Graph expressed as mean ± SEM. *$p < 0.05$, **$p < 0.01$. More detailed statistical information is in **Table S2.**

### Figure S3

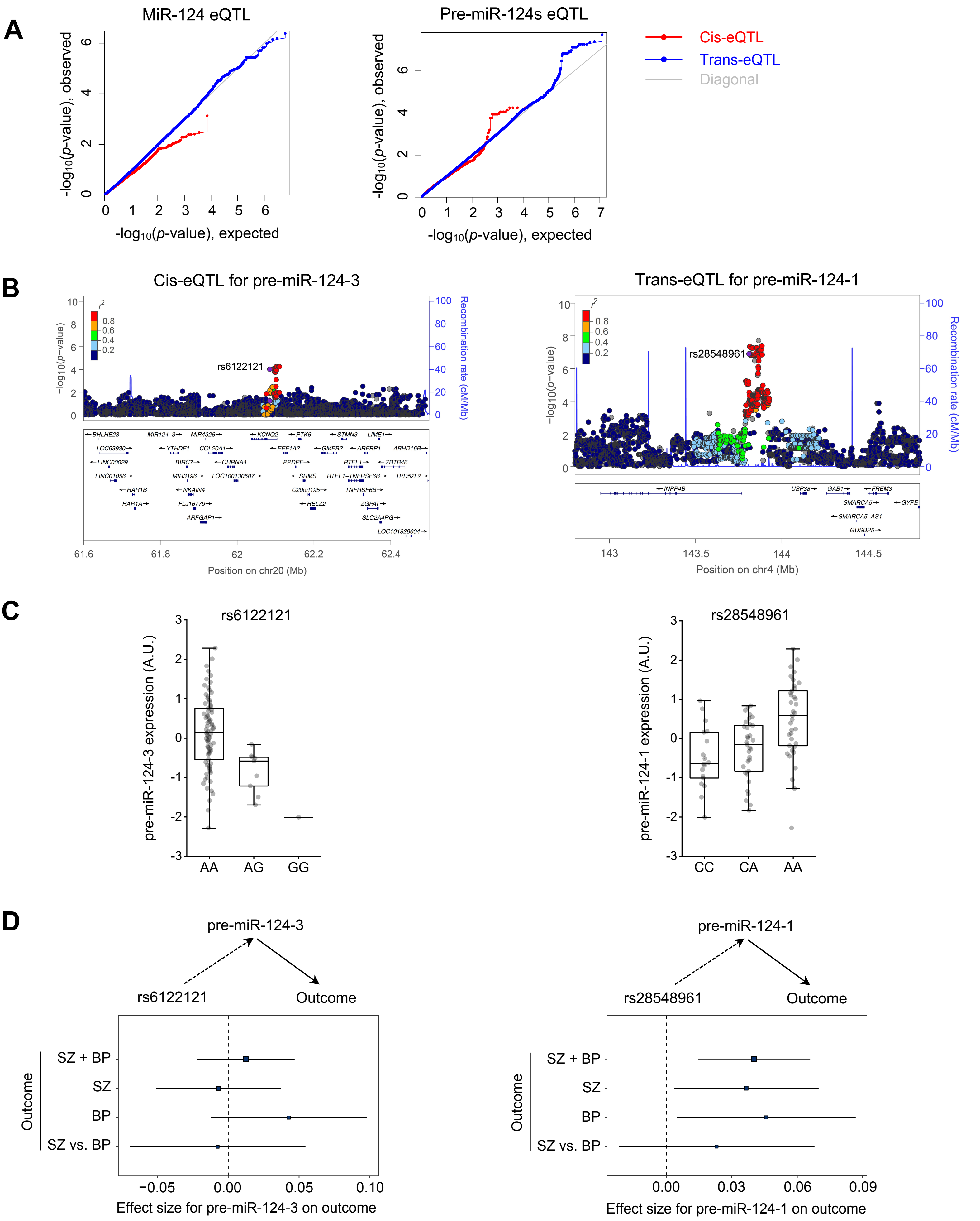
A
MiR-124 eQTL
Pre-miR-124s eQTL
-log10(p-value), observed
-log10(p-value), expected
Cis-eQTL
Trans-eQTL
Diagonal
B
Cis-eQTL for pre-miR-124-3
Trans-eQTL for pre-miR-124-1
r2
0.8
0.6
0.4
0.2
−log10(p−value)
Recombination rate (cM/Mb)
rs6122121
rs28548961
BHLHE23
MIR124−3
MIR4326
KCNQ2
PTK6
STMN3
LIME1
LOC63930
YTHDF1
COL20A1
EEF1A2
GMEB2
ARFRP1
ABHD16B
LINC00029
BIRC7
CHRNA4
PPDPF
RTEL1
ZBTB46
LINC01056
MIR3196
LOC100130587
SRMS
RTEL1−TNFRSF6B
TPD52L2
HAR1B
NKAIN4
C20orf195
TNFRSF6B
HAR1A
FLJ16779
HELZ2
ZGPAT
ARFGAP1
SLC2A4RG
LOC101928604
Position on chr20 (Mb)
INPP4B
USP38
GAB1
FREM3
SMARCA5
GYPE
SMARCA5−AS1
GUSBP5
Position on chr4 (Mb)
C
rs6122121
rs28548961
pre-miR-124-3 expression (A.U.)
pre-miR-124-1 expression (A.U.)
AA
AG
GG
CC
CA
AA
D
pre-miR-124-3
rs6122121
Outcome
pre-miR-124-1
rs28548961
Outcome
Outcome
SZ + BP
SZ
BP
SZ vs. BP
Effect size for pre-miR-124-3 on outcome
Effect size for pre-miR-124-1 on outcome

**Figure S3. Mendelian randomization (MR) showed the potential causal influence of pre-miR-124-1 on SZ or BP. (A)** Q-Q plots comparing observed distributions of association statistics against those expected under the fitted models for mature miR-124 (left) and precursor miR-124s (right). EQTL analysis identified 23 cis-eQTL for pre-miR-124-3 (red) and 37 trans-eQTL (blue) for pre-miR-124-1 at FDR<0.05. **(B)** Locus plots for the cis-eQTL on chromosome 20 (left) and trans-eQTL on 4 (right). LD clumping with the $R^2$ cutoff of 0.8 led to one independent cis- and the other independent trans-eQTL (rs6122121 and rs28548961, respectively). **(C)** Potential cis regulation of pre-miR-124-3 at rs6122121 (left) and trans regulation of pre-miR-124-1 at rs28548961 (right). rs6122121 (linear regression, $p$=9.54x$10^{-5}$, $\beta$=-1.058). rs28548961 (linear regression, $p$=1.25x$10^{-7}$, $\beta$=0.847). **(D)** Using rs28548961 as an instrument, MR showed a potential causal role of pre-miR-124-1 in SZ or BP (right), whereas a potential causal effect of pre-miR-124-3 on SZ or BP was not detected using rs6122121 as an instrument (left). For rs6122121, SZ+BP ($p$=0.476, $\beta$=0.012, se=0.017), SZ ($p$=0.761, $\beta$=-0.007, se=0.022), BP ($p$=0.128, $\beta$=0.043, se=0.028), SZ vs. BP ($p$=0.815, $\beta$=-0.007, se=0.032). For rs28548961, SZ+BP ($p$= 0.002, $\beta$= 0.04, se= 0.013), SZ ($p$= 0.03, $\beta$= 0.037, se= 0.017), BP ($p$= 0.029, $\beta$= 0.046, se= 0.021), SZ vs. BP ($p$= 0.315, $\beta$= 0.023, se= 0.023).

### Figure S4

| Rationale for homology | Humans | Rodents |
|---|---|---|
| Anatomy (Cytoarchitecture and connectivity) | BA 24 | Cg |
| | BA 32 | PrL |
| | BA 25 | IL |
| Functional features | BAs 9/46 | Cg |
| | | PrL |
| | BA 25 | IL |

**Figure S4. Discussion on cross-species homology.** While there is a consensus that the human Brodmann area 25 (BA 25) is homologous to the rodent infralimbic cortex (IL) of mPFC, the human counterparts of the mouse prelimbic cortex (PrL) and cingulate cortex (Cg) of mPFC are still under debate. The human DLPFC (BAs 9/46) shares certain functional features with the rodent PrL and Cg, whereas investigations on cytoarchitecture and connectivity have underscored the human BA 32 and BA24 as homologous with the rodent PrL and Cg, respectively. In either case, the mouse mPFC (IL, PrL, and Cg) includes brain regions homologous to many sub-regions of the human frontal lobe. This is the reason why we injected the miR-124-expressing AAV to a broad range of the mouse mPFC.

### Figure S5

**A**

| Behavioral tests | Associated dimensions |
|---|---|
| 3-chamber social interaction | Social processes |
| Psychostimulant sensitivity | Arousal / regulatory systems |
| Prepulse inhibition | Cognitive systems |
| Forced swim test | Negative valence systems |

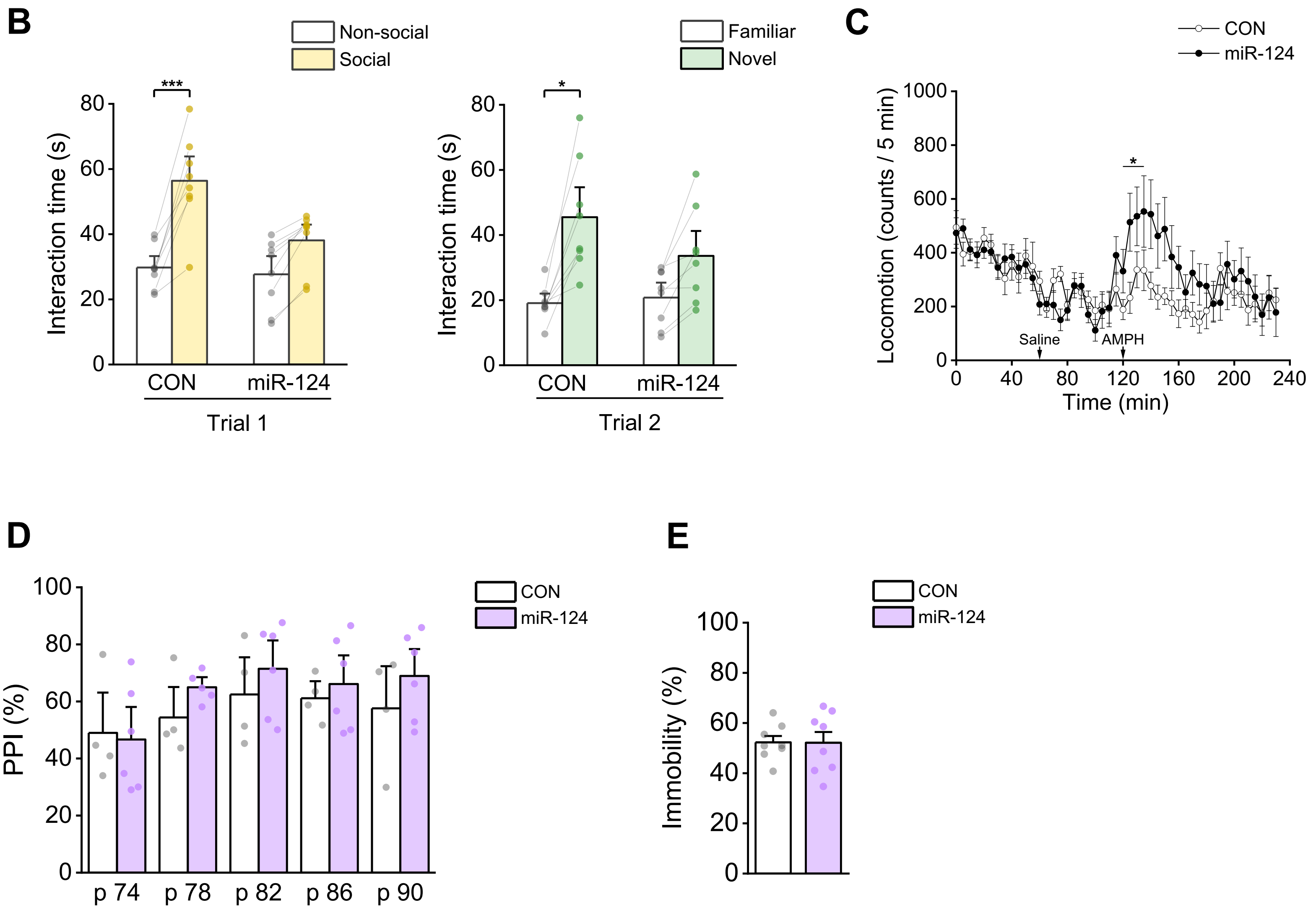

**Figure S5. Overexpression of miR-124 in mPFC neurons induced deficits in the behavioral dimensions relevant to both SZ and BP. (A)** Mice infected with miR-124 or miR-control (CON) AAV in mPFC neurons were subject to a comprehensive behavioral test battery composed of the dimensional constructs relevant to both SZ and BP. **(B)** Social interaction was assessed using three-chamber social interaction tests. The mice injected with miR-124 AAV into mPFC neurons showed behavioral deficits in both sociability (Trial 1, left) and social novelty recognition (Trial 2, right) compared to the mice injected with CON AAV. The sociability phase ($N_{CON}$=8, $N_{miR\text{-}124}$=8; two-way mixed ANOVA, AAV x Chamber, $F_{1,14}$=14.829, $p$=0.002; *post hoc* within-group comparisons using two-tailed paired Student's $t$-test). The social novelty recognition phase ($N_{CON}$=8, $N_{miR\text{-}124}$=8; two-way mixed ANOVA, AAV x Chamber, $F_{1,14}$=4.614, $p$=0.0497; *post hoc* within-group comparisons using two-tailed paired Student's $t$-test). **(C)** A psychostimulant (amphetamine, AMPH)-induced locomotion activity was measured. Upon a subcutaneous injection of AMPH, a significantly greater augmentation in the locomotion of miR-124 AAV-infected mice, in comparison to CON AAV-infected mice, was detected. The between-group contrast was most evident for the first 15 min after the injection of AMPH ($N_{CON}$=8, $N_{miR\text{-}124}$=8; two-way mixed ANOVA, AAV x Time, $F_{2,28}$=4.369, $p$=0.022; *post hoc* two-tailed paired $t$-test for each time point). **(D)** The pre-pulse inhibition of acoustic startle response was examined. The miR-124 AAV-infected mice did not differ from CON AAV-infected mice in startle response to test stimuli ($N_{CON}$=4, $N_{miR\text{-}124}$=5; pair-wise comparisons with two-tailed Welch's or Student's $t$-test, depending on homogeneity of variance). **(E)** In the forced swim test, no difference in immobility between miR-124 AAV-infected and CON AAV-infected mice was detected ($N_{CON}$=8, $N_{miR\text{-}124}$=8; two-tailed Welch's $t$-test). The bar graph expressed as mean ± SEM. *$p < 0.05$; ***$p < 0.001$. More detailed statistical information is in **Table S2.**

### Figure S6

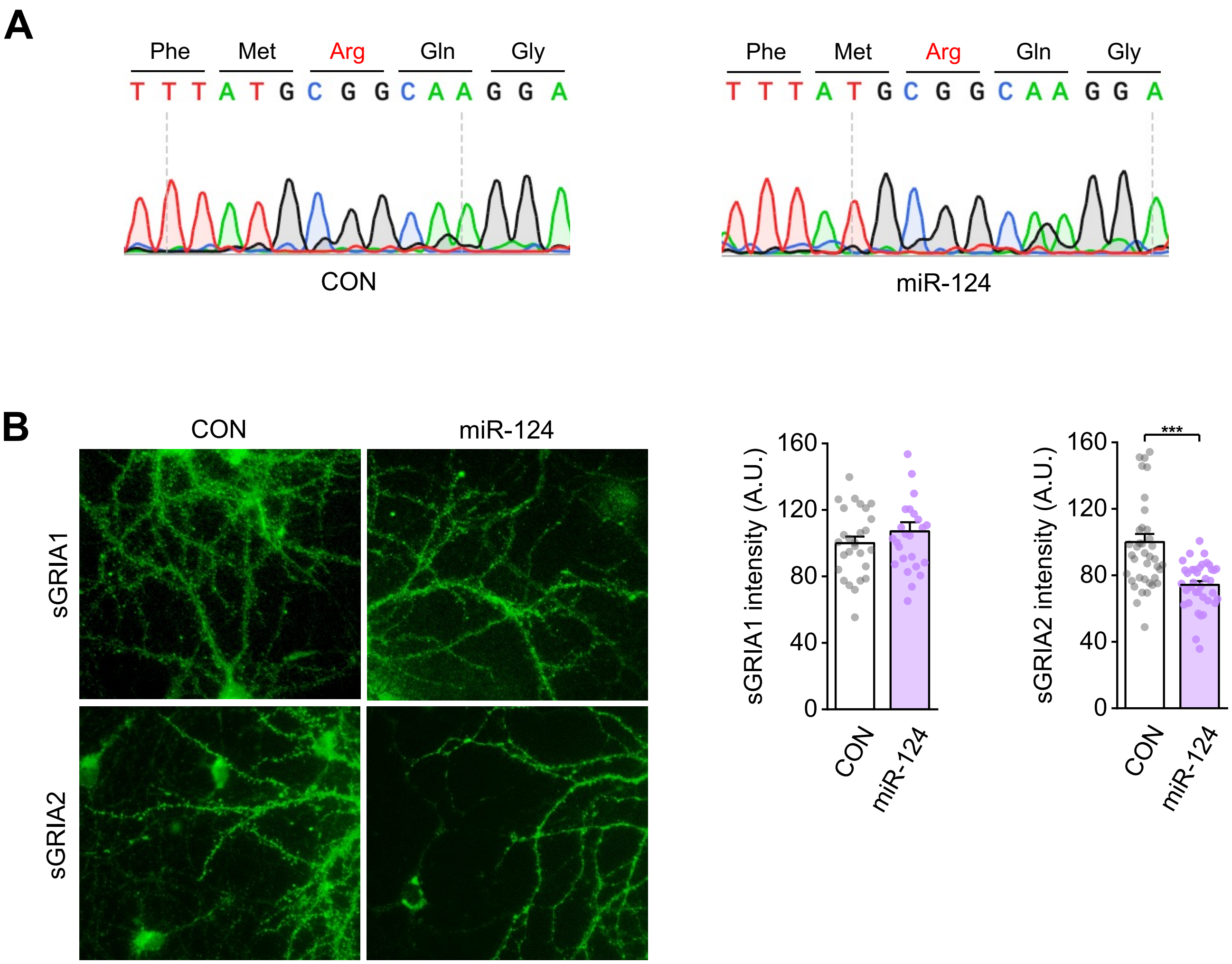
A
Phe Met Arg Gln Gly
T T T A T G C G G C A A G G A
CON
Phe Met Arg Gln Gly
T T T A T G C G G C A A G G A
miR-124
B
CON
miR-124
sGRIA1
sGRIA2
sGRIA1 intensity (A.U.)
0
40
80
120
160
CON
miR-124
sGRIA2 intensity (A.U.)
***
CON
miR-124

**Figure S6. Overexpression of miR-124 increases GRIA2-lacking calcium permeable AMPARs (CP-AMPARs). (A)** The sequencing results showed that *Gria2* transcripts from both miR-124 and miR-control (CON) AAVs-infected excitatory neurons encoded an arginine residue at the Q/R site, indicating that the modest downregulation of *Adarb1* did not affect Q/R RNA editing efficiency of *Gria2*. **(B)** Representative images of surface GRIA1 (sGRIA1) and GRIA2 (sGRIA2) in primary cortical neurons infected with CON or miR-124 AAV. sGRIA2, but not sGRIA1, was significantly decreased in cortical neurons infected with miR-124 AAV, implying the formation of GRIA2-lacking CP-AMPARs at mPFC excitatory synapses. sGRIA1 ($N_{CON}$=27, $N_{miR-124}$=25; two-tailed Student's *t*-test, $t_{50}$=-1.066, $p$=0.291). sGRIA2 ($N_{CON}$=40, $N_{miR-124}$=37; two-tailed Welch's *t*-test, $t_{55.262}$=4.774, $p=1.4\times10^{-5}$). Bar graph expressed as mean ± SEM. *$p < 0.05$, **$p < 0.01$, ***$p < 0.001$. More detailed statistical information is in **Table S2.**

**Table S1 | Pathway analysis with postmortem PFC RNA seq data**

| Rank | Database | Enriched pathway | Genes | Direction | *p*-value |
|---|---|---|---|---|---|
| 1 | Gene Ontology | single-organism process | 601 | down | 2.5E-108 |
| 2 | Gene Ontology | single-multicellular organism process | 446 | down | 1.4E-76 |
| 3 | Gene Ontology | single organism signaling | 396 | down | 1.2E-74 |
| 4 | Gene Ontology | cellular component organization or biogenesis | 340 | down | 1.1E-70 |
| 5 | Gene Ontology | cellular response to stimulus | 401 | down | 3.8E-65 |
| 6 | Gene Ontology | heterocyclic compound binding | 409 | down | 1.4E-62 |
| 7 | Gene Ontology | organic cyclic compound binding | 409 | down | 1.4E-62 |
| 8 | Gene Ontology | transport | 294 | down | 1.5E-61 |
| 9 | Gene Ontology | cellular component organization or biogenesis at cellular level | 284 | down | 8.8E-58 |
| 10 | Gene Ontology | cellular component organization at cellular level | 277 | down | 3.6E-57 |
| 11 | Broad MSigDB - Regulatory Motifs | TCF3 binding site geneset 5 | 245 | down | 1.2E-54 |
| 12 | Gene Ontology | cell periphery | 339 | down | 7.6E-53 |
| 13 | Broad MSigDB - Regulatory Motifs | SP1 binding site geneset 6 | 265 | down | 6.9E-52 |
| 14 | Gene Ontology | protein modification process | 223 | down | 5.4E-50 |
| 15 | Broad MSigDB - Regulatory Motifs | LEF1 binding site geneset 4 | 206 | down | 1.7E-49 |
| 16 | Broad MSigDB - Regulatory Motifs | Unknown transcription factor binding site geneset 3 | 198 | down | 7.9E-48 |
| 17 | Gene Ontology | regulation of nitrogen compound metabolic process | 299 | down | 9.5E-46 |
| 18 | Gene Ontology | small molecule binding | 230 | down | 9.6E-45 |
| 19 | Gene Ontology | cytosol | 219 | down | 1.8E-44 |
| 20 | Gene Ontology | nucleoside phosphate binding | 219 | down | 7.2E-44 |
| 21 | Gene Ontology | cellular aromatic compound metabolic process | 330 | down | 1.5E-42 |
| 22 | Gene Ontology | organic cyclic compound metabolic process | 337 | down | 3.9E-42 |
| 23 | Broad MSigDB - Regulatory Motifs | NFAT NFATC binding site geneset 3 | 188 | down | 5.4E-42 |
| 24 | Gene Ontology | purine ribonucleoside triphosphate binding | 183 | down | 1.1E-41 |
| 25 | Gene Ontology | heterocycle metabolic process | 327 | down | 2.1E-41 |
| 26 | Gene Ontology | cell differentiation | 209 | down | 7.9E-41 |
| 27 | Broad MSigDB - Regulatory Motifs | MLLT7 binding site geneset 3 | 192 | down | 5.1E-39 |
| 28 | Broad MSigDB - Regulatory Motifs | MAZ binding site geneset 2 | 199 | down | 3.1E-37 |
| 29 | Gene Ontology | regulation of response to stimulus | 202 | down | 7.3E-36 |
| 30 | Gene Ontology | response to stress | 215 | down | 1.3E-35 |
| 31 | Gene Ontology | regulation of biosynthetic process | 262 | down | 1.9E-34 |
| 32 | Gene Ontology | multicellular organismal signaling | 92 | down | 4.2E-32 |
| 33 | Gene Ontology | regulation of cellular macromolecule biosynthetic process | 244 | down | 4.6E-32 |
| 34 | Gene Ontology | anatomical structure formation involved in morphogenesis | 138 | down | 7.7E-32 |
| 35 | Gene Ontology | transmission of nerve impulse | 90 | down | 2.1E-31 |
| 36 | Gene Ontology | catabolic process | 172 | down | 7.7E-31 |
| 37 | Gene Ontology | neurogenesis | 119 | down | 1.3E-30 |
| 38 | Gene Ontology | ion transport | 106 | down | 7.1E-30 |
| 39 | Broad MSigDB - Regulatory Motifs | ESRRA binding site geneset 2 | 115 | down | 7.3E-30 |
| 40 | Broad MSigDB - Regulatory Motifs | REPIN1 binding site geneset 1 | 144 | down | 7.6E-30 |
| 41 | Broad MSigDB - Regulatory Motifs | PAX4 binding site geneset 5 | 129 | down | 1.9E-29 |
| 42 | Gene Ontology | regulation of RNA biosynthetic process | 224 | down | 1.5E-28 |
| 43 | Gene Ontology | transporter activity | 118 | down | 4.3E-28 |

| | | | | | |
|---|---|---|---|---|---|
| 44 | Gene Ontology | cytoskeleton | 154 | down | 4.7E-28 |
| 45 | Gene Ontology | regulation of transport | 110 | down | 7.9E-28 |
| 46 | Gene Ontology | synaptic transmission | 79 | down | 1.4E-27 |
| 47 | Gene Ontology | nucleolus | 139 | down | 1.7E-27 |
| 48 | Gene Ontology | neuron differentiation | 93 | down | 1.4E-26 |
| 49 | Gene Ontology | cell-cell signaling | 97 | down | 1.9E-26 |
| 50 | Gene Ontology | Gene Ontologylgi apparatus | 115 | down | 6.2E-26 |
| 51 | Broad MSigDB - Regulatory Motifs | Unknown transcription factor binding site geneset 70 | 104 | down | 6.2E-26 |
| 52 | Gene Ontology | neuron development | 82 | down | 8.2E-26 |
| 53 | Gene Ontology | cell death | 110 | down | 1E-25 |
| 54 | Gene Ontology | death | 110 | down | 1.2E-25 |
| 55 | Broad MSigDB - Regulatory Motifs | JUN binding site geneset 11 | 112 | down | 2.4E-25 |
| 56 | TargetScan miRNA targets DB | Predicted Gene Targets for miR-30 | 83 | down | 4E-25 |
| 57 | Gene Ontology | mitochondrion | 131 | down | 5.6E-25 |
| 58 | Gene Ontology | cell part morphogenesis | 76 | down | 1.4E-24 |
| 59 | Gene Ontology | regulation of cellular component organization | 113 | down | 1.6E-24 |
| 60 | Gene Ontology | cellular response to stress | 104 | down | 3.2E-24 |
| 61 | TargetScan miRNA targets DB | Predicted Gene Targets for miR-182 | 70 | down | 3.7E-24 |
| 62 | Gene Ontology | cell cycle | 110 | down | 5.4E-24 |
| 63 | Broad MSigDB - Canonical Pathways | Genes involved in Immune System | 97 | down | 5.5E-24 |
| 64 | Gene Ontology | cell morphogenesis involved in neuron differentiation | 67 | down | 5.8E-24 |
| 65 | Gene Ontology | neuron projection development | 72 | down | 2.1E-23 |
| 66 | Broad MSigDB - Regulatory Motifs | LEF1 binding site geneset 3 | 114 | down | 5E-23 |
| 67 | Broad MSigDB - Regulatory Motifs | FOXF2 binding site geneset 3 | 95 | down | 5.6E-23 |
| 68 | TargetScan miRNA targets DB | Predicted Gene Targets for miR-128 | 68 | down | 6.2E-23 |
| 69 | Gene Ontology | cell morphogenesis involved in differentiation | 72 | down | 7.2E-23 |
| 70 | InterPro | Armadillo-type fold | 51 | down | 9.8E-23 |
| 71 | Gene Ontology | nucleoplasm | 125 | down | 2.2E-22 |
| 72 | Gene Ontology | endoplasmic reticulum | 114 | down | 2.4E-22 |
| 73 | Gene Ontology | central nervous system development | 60 | up | 2.4E-22 |
| 74 | TargetScan miRNA targets DB | Predicted Gene Targets for miR-153 | 55 | down | 2.7E-22 |
| 75 | TargetScan miRNA targets DB | Predicted Gene Targets for miR-124u | 72 | down | 3.1E-22 |
| 76 | Broad MSigDB - Regulatory Motifs | Unknown transcription factor binding site geneset 64 | 84 | down | 4.3E-22 |
| 77 | Gene Ontology | response to organic substance | 102 | up | 4.8E-22 |
| 78 | Broad MSigDB - Regulatory Motifs | MYOD1 binding site geneset 4 | 94 | down | 5.7E-22 |
| 79 | Broad MSigDB - Regulatory Motifs | Targets of MicroRNA TGTTTAC,MIR-30A-5P,MIR-30C,MIR-30D,MIR-3 | 72 | down | 6.3E-22 |
| 80 | Broad MSigDB - Regulatory Motifs | Targets of MicroRNA TTTGCAC,MIR-19A,MIR-19B | 67 | down | 9.1E-22 |
| 81 | Broad MSigDB - Regulatory Motifs | MYC binding site geneset 6 | 100 | down | 9.5E-22 |
| 82 | TargetScan miRNA targets DB | Predicted Gene Targets for miR-96 | 69 | down | 1.1E-21 |
| 83 | TargetScan miRNA targets DB | Predicted Gene Targets for miR-26 | 64 | down | 1.2E-21 |
| **84** | **TargetScan miRNA targets DB** | **Predicted Gene Targets for miR-124a** | **90** | **down** | **1.3E-21** |
| 85 | Broad MSigDB - Regulatory Motifs | Targets of MicroRNA ACTGTGA,MIR-27A,MIR-27B | 64 | down | 1.6E-21 |
| 86 | TargetScan miRNA targets DB | Predicted Gene Targets for miR-9 | 76 | down | 1.6E-21 |
| 87 | TargetScan miRNA targets DB | Predicted Gene Targets for miR-25 | 60 | down | 1.8E-21 |
| 88 | Broad MSigDB - Regulatory Motifs | Unknown transcription factor binding site geneset 175 | 106 | down | 3E-21 |

| | | | | | |
|---|---|---|---|---|---|
| 89 | Gene Ontology | kinase activity | 82 | down | 4.5E-21 |
| 90 | Gene Ontology | enzyme binding | 99 | down | 9.2E-21 |
| 91 | Gene Ontology | cytoskeletal protein binding | 72 | down | 1.6E-20 |
| 92 | Broad MSigDB - Canonical Pathways | Genes involved in Adaptive Immune System | 65 | down | 2.9E-20 |
| 93 | Broad MSigDB - Regulatory Motifs | Unknown transcription factor binding site geneset 142 | 73 | up | 3.5E-20 |
| 94 | Broad MSigDB - Regulatory Motifs | TCF8 binding site geneset 5 | 82 | down | 6.7E-20 |
| 95 | TargetScan miRNA targets DB | Predicted Gene Targets for miR-448 | 52 | down | 1.2E-19 |
| 96 | TargetScan miRNA targets DB | Predicted Gene Targets for miR-496 | 61 | down | 2.8E-19 |
| 97 | Broad MSigDB - Regulatory Motifs | MEF2A binding site geneset 3 | 75 | down | 3.6E-19 |
| 98 | TargetScan miRNA targets DB | Predicted Gene Targets for miR-15 | 66 | down | 4E-19 |
| 99 | TargetScan miRNA targets DB | Predicted Gene Targets for miR-27 | 67 | down | 5.4E-19 |
| 100 | Gene Ontology | neuron projection | 68 | down | 5.7E-19 |
| 101 | Gene Ontology | regulation of cell differentiation | 71 | up | 7.5E-19 |
| 102 | Broad MSigDB - Regulatory Motifs | NF1 binding site geneset 4 | 76 | down | 8.3E-19 |
| 103 | TargetScan miRNA targets DB | Predicted Gene Targets for miR-19 | 63 | down | 1E-18 |
| 104 | Broad MSigDB - Regulatory Motifs | VSX1 binding site geneset 2 | 76 | down | 1.5E-18 |
| 105 | Broad MSigDB - Regulatory Motifs | Targets of MicroRNA TGCTGCT,MIR-15A,MIR-16,MIR-15B,MIR-195,M | 67 | down | 3.7E-18 |
| 106 | Gene Ontology | enzyme regulator activity | 88 | down | 5.1E-18 |
| 107 | Gene Ontology | ion channel activity | 49 | down | 5.4E-18 |
| 108 | TargetScan miRNA targets DB | Predicted Gene Targets for miR-381 | 60 | down | 6.3E-18 |
| 109 | Gene Ontology | passive transmembrane transporter activity | 50 | down | 8.2E-18 |
| 110 | Broad MSigDB - Regulatory Motifs | TAF TATA binding site geneset 3 | 83 | up | 9.4E-18 |
| 111 | Gene Ontology | locomotion | 87 | down | 1E-17 |
| 112 | Gene Ontology | cell junction | 73 | down | 1E-17 |
| 113 | TargetScan miRNA targets DB | Predicted Gene Targets for miR-218 | 54 | down | 1.1E-17 |
| 114 | Gene Ontology | brain development | 43 | up | 1.5E-17 |
| 115 | Broad MSigDB - Regulatory Motifs | POU1F1 binding site geneset 2 | 33 | up | 1.8E-17 |
| 116 | Broad MSigDB - Regulatory Motifs | Targets of MicroRNA TGGTGCT,MIR-29A,MIR-29B,MIR-29C | 49 | up | 2E-17 |
| 117 | Gene Ontology | protein kinase activity | 65 | down | 2.3E-17 |
| 118 | Broad MSigDB - Canonical Pathways | Genes involved in Antigen processing: Ubiquitination & Proteasome de | 37 | down | 2.4E-17 |
| 119 | Broad MSigDB - Regulatory Motifs | MEIS1 binding site geneset 4 | 79 | down | 2.5E-17 |
| 120 | Gene Ontology | cellular component assembly at cellular level | 83 | down | 3E-17 |
| 121 | TargetScan miRNA targets DB | Predicted Gene Targets for miR-495 | 46 | up | 4.1E-17 |
| 122 | Broad MSigDB - Regulatory Motifs | Targets of MicroRNA GTGCCTT,MIR-506 | 73 | down | 4.4E-17 |
| 123 | TargetScan miRNA targets DB | Predicted Gene Targets for miR-103 | 44 | down | 5E-17 |
| 124 | Broad MSigDB - Regulatory Motifs | ETS2 binding site geneset 2 | 73 | up | 7.1E-17 |
| 125 | Broad MSigDB - Regulatory Motifs | NRF1 binding site geneset 2 | 83 | down | 7.2E-17 |
| 126 | Broad MSigDB - Regulatory Motifs | ELK1 binding site geneset 3 | 99 | down | 7.6E-17 |
| 127 | Broad MSigDB - Regulatory Motifs | Unknown transcription factor binding site geneset 150 | 48 | up | 8.1E-17 |
| 128 | Broad MSigDB - Canonical Pathways | Genes involved in Neuronal System | 41 | down | 8.3E-17 |
| 129 | TargetScan miRNA targets DB | Predicted Gene Targets for miR-142-5p | 49 | down | 1.1E-16 |
| 130 | Gene Ontology | regulation of ion transport | 46 | down | 1.5E-16 |
| 131 | Broad MSigDB - Regulatory Motifs | E2F1 binding site geneset 7 | 39 | down | 2E-16 |
| 132 | Broad MSigDB - Regulatory Motifs | Targets of MicroRNA AAGCCAT,MIR-135A,MIR-135B | 46 | down | 2E-16 |
| 133 | Gene Ontology | envelope | 77 | down | 2.3E-16 |

| | | | | | |
|---|---|---|---|---|---|
| 134 | Broad MSigDB - Regulatory Motifs | Unknown transcription factor binding site geneset 149 | 65 | down | 2.5E-16 |
| 135 | Broad MSigDB - Regulatory Motifs | EGR4 binding site geneset 1 | 40 | down | 2.8E-16 |
| 136 | TargetScan miRNA targets DB | Predicted Gene Targets for miR-144 | 50 | down | 3.1E-16 |
| 137 | TargetScan miRNA targets DB | Predicted Gene Targets for miR-30-3p | 39 | down | 3.3E-16 |
| 138 | Broad MSigDB - Regulatory Motifs | FOXA1 binding site geneset 3 | 66 | down | 3.3E-16 |
| 139 | TargetScan miRNA targets DB | Predicted Gene Targets for miR-101 | 51 | down | 4E-16 |
| 140 | Broad MSigDB - Regulatory Motifs | Unknown transcription factor binding site geneset 176 | 39 | up | 4.3E-16 |
| 141 | Gene Ontology | purine-containing compound metabolic process | 73 | down | 4.4E-16 |
| 142 | Gene Ontology | ligase activity | 56 | down | 4.7E-16 |
| 143 | Gene Ontology | cell projection part | 63 | down | 5E-16 |
| 144 | Broad MSigDB - Regulatory Motifs | Targets of MicroRNA AAGCACA,MIR-218 | 50 | down | 6.7E-16 |
| 145 | Broad MSigDB - Canonical Pathways | Ubiquitin mediated proteolysis | 29 | down | 6.8E-16 |
| 146 | Gene Ontology | positive regulation of biosynthetic process | 99 | down | 7.9E-16 |
| 147 | Broad MSigDB - Regulatory Motifs | Targets of MicroRNA CACTGTG,MIR-128A,MIR-128B | 45 | down | 9E-16 |
| 148 | TargetScan miRNA targets DB | Predicted Gene Targets for miR-29 | 46 | up | 9.4E-16 |
| 149 | Gene Ontology | cytoskeleton organization | 64 | down | 9.7E-16 |
| 150 | Broad MSigDB - Canonical Pathways | Genes involved in Class I MHC mediated antigen processing & presen | 38 | down | 1.2E-15 |
| 151 | Gene Ontology | cellular homeostasis | 64 | down | 1.3E-15 |
| 152 | Gene Ontology | membrane-bounded vesicle | 78 | down | 1.5E-15 |
| 153 | Broad MSigDB - Regulatory Motifs | NFIL3 binding site geneset 2 | 39 | down | 1.7E-15 |
| 154 | Gene Ontology | regulation of protein modification process | 86 | down | 1.7E-15 |
| 155 | TargetScan miRNA targets DB | Predicted Gene Targets for miR-135 | 45 | down | 1.9E-15 |
| 156 | Gene Ontology | regulation of multicellular organismal development | 83 | down | 3.3E-15 |
| 157 | Gene Ontology | lipid binding | 70 | down | 3.5E-15 |
| 158 | Gene Ontology | extracellular matrix | 36 | down | 3.7E-15 |
| 159 | Gene Ontology | ligase activity, forming carbon-nitrogen bonds | 44 | down | 3.8E-15 |
| 160 | Gene Ontology | GTPase regulator activity | 51 | down | 3.8E-15 |
| 161 | Gene Ontology | cell division | 49 | down | 4.1E-15 |
| 162 | Gene Ontology | regulation of cellular component movement | 50 | down | 4.3E-15 |
| 163 | Broad MSigDB - Canonical Pathways | Genes involved in Axon guidance | 37 | down | 4.3E-15 |
| 164 | Gene Ontology | cytoplasmic membrane-bounded vesicle | 76 | down | 4.4E-15 |
| 165 | Gene Ontology | phospholipid binding | 55 | down | 5.5E-15 |
| 166 | Gene Ontology | telencephalon development | 24 | up | 5.9E-15 |
| 167 | Gene Ontology | forebrain development | 30 | up | 6.1E-15 |
| 168 | Gene Ontology | response to wounding | 76 | down | 7.1E-15 |
| 169 | Gene Ontology | proteinaceous extracellular matrix | 31 | down | 9.5E-15 |
| 170 | Gene Ontology | response to DNA damage stimulus | 58 | down | 1.1E-14 |
| 171 | Broad MSigDB - Regulatory Motifs | NR3C1 binding site geneset 5 | 32 | up | 1.1E-14 |
| 172 | Broad MSigDB - Regulatory Motifs | Targets of MicroRNA TTGCCAA,MIR-182 | 43 | down | 1.4E-14 |
| 173 | Gene Ontology | regulation of neurogenesis | 37 | up | 1.4E-14 |
| 174 | Broad MSigDB - Regulatory Motifs | POU3F2 binding site geneset 1 | 30 | up | 1.4E-14 |
| 175 | TargetScan miRNA targets DB | Predicted Gene Targets for let-7 | 53 | down | 1.6E-14 |
| 176 | Gene Ontology | regulation of phosphorus metabolic process | 63 | up | 1.6E-14 |
| 177 | TargetScan miRNA targets DB | Predicted Gene Targets for miR-203 | 43 | down | 1.7E-14 |
| 178 | Gene Ontology | circulatory system development | 60 | down | 2E-14 |

| | | | | | |
|---|---|---|---|---|---|
| 179 | Gene Ontology | cardiovascular system development | 60 | down | 2E-14 |
| 180 | Broad MSigDB - Regulatory Motifs | Targets of MicroRNA CTTTGTA,MIR-524 | 50 | down | 2.2E-14 |
| 181 | Gene Ontology | protein transport | 83 | down | 2.3E-14 |
| 182 | Gene Ontology | response to external stimulus | 86 | down | 2.4E-14 |
| 183 | Gene Ontology | negative regulation of biosynthetic process | 82 | down | 2.5E-14 |
| 184 | Gene Ontology | cytoplasmic vesicle | 78 | down | 2.7E-14 |
| 185 | Gene Ontology | chemotaxis | 56 | down | 2.8E-14 |
| 186 | Gene Ontology | taxis | 56 | down | 2.8E-14 |
| 187 | Gene Ontology | vesicle | 80 | down | 2.9E-14 |
| 188 | Gene Ontology | regulation of locomotion | 48 | down | 3.2E-14 |
| 189 | Gene Ontology | Gene Ontologylgi apparatus part | 63 | down | 3.8E-14 |
| 190 | Broad MSigDB - Regulatory Motifs | Targets of MicroRNA ATGCTGC,MIR-103,MIR-107 | 34 | down | 4.3E-14 |
| 191 | Gene Ontology | generation of precursor metabolites and energy | 46 | down | 4.5E-14 |
| 192 | InterPro | WD40/YVTN repeat-like | 42 | down | 5E-14 |
| 193 | Gene Ontology | pallium development | 19 | up | 5.2E-14 |
| 194 | Gene Ontology | identical protein binding | 72 | down | 5.2E-14 |
| 195 | Gene Ontology | acid-amino acid ligase activity | 40 | down | 5.3E-14 |
| 196 | TargetScan miRNA targets DB | Predicted Gene Targets for miR-323 | 46 | down | 5.3E-14 |
| 197 | Broad MSigDB - Regulatory Motifs | Targets of MicroRNA GCACTTT,MIR-17-5P,MIR-20A,MIR-106A,MIR-1 | 59 | down | 5.4E-14 |
| 198 | Broad MSigDB - Regulatory Motifs | Unknown transcription factor binding site geneset 82 | 29 | up | 5.6E-14 |
| 199 | Gene Ontology | microtubule organizing center | 52 | down | 6.3E-14 |
| 200 | Gene Ontology | cellular component movement | 56 | up | 6.5E-14 |

### Table S2 | Statistics summary

#### Figure 1

| Figure | Test | Measurement | Group | Sub-group | Sample size | Average | Standard deviation | Statistical test | Statistical result | *post hoc* test | Result from the *post hoc* tests |
|---|---|---|---|---|---|---|---|---|---|---|---|
| 1B | qRT-PCR for miR-124 in OE | Relative expression (A.U.) | CON | - | 38 | 1.000 | 0.507 | Welch's ANOVA | $F(2,52.335)=14.100$, $p=0.000013$ | Games-Howell (CON vs. SZ) | $p=0.004586$ |
| | | | SZ | | 37 | 1.614 | 1.007 | | | Games-Howell (CON vs. BP) | $p=0.000073$ |
| | | | BP | | 24 | 1.811 | 0.721 | | | Games-Howell (SZ vs. BP) | $p=0.650228$ |
| 1C | qRT-PCR for miR-17 | Relative expression (A.U.) | CON | - | 39 | 1.0000 | 0.71674 | One-way ANOVA | $F(2,98)=3.246$, $p=0.043152$ | Tukey's HSD (CON vs. SZ) | $p=0.038438$ |
| | | | SZ | | 37 | 1.4457 | 0.82436 | | | Tukey's HSD (CON vs. BP) | $p=0.257029$ |
| | | | BP | | 25 | 1.3171 | 0.81118 | | | Tukey's HSD (SZ vs. BP) | $p=0.800740$ |
| | RT-qPCR for miR-107 | Relative expression (A.U.) | CON | - | 38 | 1.0000 | 0.49561 | One-way ANOVA | $F(2,98)=3.393$, $p=0.037614$ | Tukey's HSD (CON vs. SZ) | $p=0.036222$ |
| | | | SZ | | 38 | 1.4586 | 0.91749 | | | Tukey's HSD (CON vs. BP) | $p=0.195089$ |
| | | | BP | | 25 | 1.3572 | 0.95589 | | | Tukey's HSD (SZ vs. BP) | $p=0.874009$ |
| | RT-qPCR for miR-132 | Relative expression (A.U.) | CON | - | 39 | 1 | 0.364192 | Welch's ANOVA | $F(2,55.228)=4.552$, $p=0.014797$ | Games-Howell (CON vs. SZ) | $p=0.015387$ |
| | | | SZ | | 38 | 1.350925 | 0.660894 | | | Games-Howell (CON vs. BP) | $p=0.242975$ |
| | | | BP | | 25 | 1.1896 | 0.50111 | | | Games-Howell (SZ vs. BP) | $p=0.518220$ |
| | RT-qPCR for miR-134 | Relative expression (A.U.) | CON | - | 39 | 1 | 0.473222 | Welch's ANOVA | $F(2,59.049)=5.428$, $p=0.006854$ | Games-Howell (CON vs. SZ) | $p=0.039894$ |
| | | | SZ | | 38 | 1.380893 | 0.815285 | | | Games-Howell (CON vs. BP) | $p=0.019064$ |
| | | | BP | | 25 | 1.361035 | 0.517764 | | | Games-Howell (SZ vs. BP) | $p=0.992325$ |
| | RT-qPCR for miR-137 | Relative expression (A.U.) | CON | - | 38 | 1 | 0.65484 | Welch's ANOVA | $F(2,49.415)=3.811$, $p=0.028896$ | Games-Howell (CON vs. SZ) | $p=0.032729$ |
| | | | SZ | | 37 | 1.55340 | 1.13236 | | | Games-Howell (CON vs. BP) | $p=0.280200$ |
| | | | BP | | 23 | 1.38510 | 1.07404 | | | Games-Howell (SZ vs. BP) | $p=0.832442$ |
| | RT-qPCR for miR-652 | Relative expression (A.U.) | CON | - | 39 | 1 | 0.72139 | One-way ANOVA | $F(2,99)=3.117$, $p=0.048685$ | Tukey's HSD (CON vs. SZ) | $p=0.038592$ |
| | | | SZ | | 38 | 1.43710 | 0.92344 | | | Tukey's HSD (CON vs. BP) | $p=0.660325$ |
| | | | BP | | 25 | 1.17210 | 0.56424 | | | Tukey's HSD (SZ vs. BP) | $p=0.380153$ |
| 1D | qRT-PCR for miR-124 in PFC | Relative expression (A.U.) | CON | - | 15 | 1.0000 | 0.34448 | One-way ANOVA | $F(3,58)=4.695$, $p=0.005$ | Tukey's HSD | CON vs. SZ: $p=0.048512$<br>CON vs. BP: $p=0.014913$<br>CON vs. MDD: $p=0.999587$<br>SZ vs. BP: $p=0.858130$<br>SZ vs. MDD: $p=0.101950$<br>BP vs. MDD: $p=0.032423$ |
| | | | SZ | | 15 | 1.4286 | 0.43325 | | | | |
| | | | BP | | 14 | 1.5228 | 0.63008 | | | | |
| | | | MDD | | 15 | 1.0183 | 0.48232 | | | | |
| | qRT-PCR for miR-124 in LB | Relative expression (A.U.) | CON | - | 22 | 1 | 0.74794 | - | - | Two-tailed Student's t-test (CON vs. SZ) | $t(43)=-0.415$, $p=0.754$ |
| | | | SZ | | 23 | 1.0873 | 0.66201 | | | | |

#### Figure 2

| Figure | Test | Measurement | Group | Sub-group | Sample size | Average | Standard deviation | Statistical test | Statistical result | *post hoc* test | Result from the *post hoc* tests |
|---|---|---|---|---|---|---|---|---|---|---|---|
| 2B | Association analysis between PRSs and psychosis-control status | Nagelkerke's $R^2$ | SZ+BP PRS | $p_T$<0.0001 | CON (45)<br>Case (63) = SZ (38) + BP(25) | - | - | Logistic regression without covariates | Nagelkerke's $R^2$(%)=8.99, $p$=6.57E-3 | Logistic regression with covariates (age, gender, and PCs) | Nagelkerke's $R^2$(PRS) (%)=5.441, $p$(PRS)=0.0238 |
| | | | | $p_T$<0.001 | | | | | Nagelkerke's $R^2$(%)=10.382, $p$=3.40E-3 | | Nagelkerke's $R^2$(PRS) (%)=7.816, $p$(PRS)=7.37E-3 |
| | | | | $p_T$<0.01 | | | | | Nagelkerke's $R^2$(%)=16.659, $p$=1.72E-4 | | Nagelkerke's $R^2$(PRS) (%)=13.460, $p$(PRS)=8.03E-4 |
| | | | | $p_T$<0.05 | | | | | Nagelkerke's $R^2$(%)=19.337, $p$=4.69E-5 | | Nagelkerke's $R^2$(PRS) (%)=13.838, $p$(PRS)=7.35E-4 |
| | | | | $p_T$<0.1 | | | | | Nagelkerke's $R^2$(%)=19.202, $p$=5.01E-5 | | Nagelkerke's $R^2$(PRS) (%)=14.008, $p$(PRS)=6.45E-4 |
| | | | | $p_T$<0.2 | | | | | Nagelkerke's $R^2$(%)=17.983, $p$=9.06E-5 | | Nagelkerke's $R^2$(PRS) (%)=12.472, $p$(PRS)=1.17E-3 |
| | | | | $p_T$<0.3 | | | | | Nagelkerke's $R^2$(%)=17.155, $p$=1.35E-4 | | Nagelkerke's $R^2$(PRS) (%)=12.134, $p$(PRS)=1.37E-3 |
| | | | | $p_T$<0.4 | | | | | Nagelkerke's $R^2$(%)=17.461, $p$=1.17E-4 | | Nagelkerke's $R^2$(PRS) (%)=12.341, $p$(PRS)=1.25E-3 |
| | | | | $p_T$<0.5 | | | | | Nagelkerke's $R^2$(%)=18.188, $p$=8.20E-5 | | Nagelkerke's $R^2$(PRS) (%)=12.656, $p$(PRS)=1.16E-3 |
| | | | | $p_T$<1 | | | | | Nagelkerke's $R^2$(%)=19.169, $p$=5.09E-5 | | Nagelkerke's $R^2$(PRS) (%)=13.092, $p$(PRS)=9.48E-4 |
| | | | SZ PRS | $p_T$<0.0001 | CON (45)<br>Case (63) = SZ (38) + BP(25) | - | - | Logistic regression without covariates | Nagelkerke's $R^2$(%)=7.467 , $p$=0.0135 | Logistic regression with covariates (age, gender, and PCs) | Nagelkerke's $R^2$(PRS) (%)=4.747, $p$(PRS)=0.033 |
| | | | | $p_T$<0.001 | | | | | Nagelkerke's $R^2$(%)=5.844 , $p$=0.029 | | Nagelkerke's $R^2$PRS (%)=4.527 , $p$(PRS)=0.036 |
| | | | | $p_T$<0.01 | | | | | Nagelkerke's $R^2$(%)=2.614 , $p$=0.148 | | Nagelkerke's $R^2$(PRS) (%)=4.687 , $p$(PRS)=0.034 |
| | | | | $p_T$<0.05 | | | | | Nagelkerke's $R^2$(%)=6.896 , $p$=0.018 | | Nagelkerke's $R^2$(PRS) (%)=7.205 , $p$(PRS)=0.011 |
| | | | | $p_T$<0.1 | | | | | Nagelkerke's $R^2$(%)=8.028 , $p$=0.010 | | Nagelkerke's $R^2$(PRS) (%)=8.123 , $p$(PRS)=6.79E-3 |
| | | | | $p_T$<0.2 | | | | | Nagelkerke's $R^2$(%)=14.705 , $p$=4.38E-4 | | Nagelkerke's $R^2$(PRS) (%)=12.203 , $p$(PRS)=1.23E-3 |
| | | | | $p_T$<0.3 | | | | | Nagelkerke's $R^2$(%)=13.264 , $p$=8.7E-4 | | Nagelkerke's $R^2$(PRS) (%)=12.097 , $p$(PRS)=1.26E-3 |
| | | | | $p_T$<0.4 | | | | | Nagelkerke's $R^2$(%)=13.771, $p$=6.84E-4 | | Nagelkerke's $R^2$(PRS) (%)=13.311, $p$(PRS)=7.89E-4 |
| | | | | $p_T$<0.5 | | | | | Nagelkerke's $R^2$(%)=14.722 , $p$=4.35E-4 | | Nagelkerke's $R^2$(PRS) (%)=13.616 , $p$(PRS)=6.98E-4 |
| | | | | $p_T$<1 | | | | | Nagelkerke's $R^2$(%)=13.031 , $p$=9.72E-4 | | Nagelkerke's $R^2$(PRS) (%)=11.893 , $p$(PRS)=1.34E-3 |
| | | | BP PRS | $p_T$<0.0001 | CON (45)<br>Case (63) = SZ (38) + BP(25) | - | - | Logistic regression without covariates | Nagelkerke's $R^2$(%)=9.376 , $p$=5.48E-3 | Logistic regression with covariates (age, gender, and PCs) | Nagelkerke's $R^2$(PRS) (%)=5.863 , $p$(PRS)= 0.020 |
| | | | | $p_T$<0.001 | | | | | Nagelkerke's $R^2$(%)=7.153 , $p$=0.016 | | Nagelkerke's $R^2$(PRS) (%)=4.692 , $p$(PRS)=0.037 |
| | | | | $p_T$<0.01 | | | | | Nagelkerke's $R^2$(%)=10.565 , $p$=3.12E-3 | | Nagelkerke's $R^2$(PRS) (%)=6.317 , $p$(PRS)=0.016 |
| | | | | $p_T$<0.05 | | | | | Nagelkerke's $R^2$(%)=9.508 , $p$=5.15E-3 | | Nagelkerke's $R^2$(PRS) (%)=6.620 , $p$(PRS)=0.014 |
| | | | | $p_T$<0.1 | | | | | Nagelkerke's $R^2$(%)=11.051 , $p$=2.48E-3 | | Nagelkerke's $R^2$(PRS) (%)=7.040 , $p$(PRS)=0.011 |
| | | | | $p_T$<0.2 | | | | | Nagelkerke's $R^2$(%)=13.345 , $p$=8.38E-4 | | Nagelkerke's $R^2$(PRS) (%)=9.248 , $p$(PRS)=4.25E-3 |
| | | | | $p_T$<0.3 | | | | | Nagelkerke's $R^2$(%)=12.403 , $p$=1.39E-3 | | Nagelkerke's $R^2$(PRS) (%)=7.418 , $p$(PRS)=9.81E-3 |
| | | | | $p_T$<0.4 | | | | | Nagelkerke's $R^2$(%)=12.259 , $p$=1.40E-3 | | Nagelkerke's $R^2$(PRS) (%)=7.364 , $p$(PRS)=9.84E-3 |
| | | | | $p_T$<0.5 | | | | | Nagelkerke's $R^2$(%)=10.905 , $p$=2.66E-3 | | Nagelkerke's $R^2$(PRS) (%)=6.049 , $p$(PRS)=0.019 |
| | | | | $p_T$<1 | | | | | Nagelkerke's $R^2$(%)=11.633 , $p$=1.86E-3 | | Nagelkerke's $R^2$(PRS) (%)=6.019 , $p$(PRS)=0.019 |
| | | | SZ vs. BP PRS | $p_T$<0.0001 | CON (45)<br>Case (63) = SZ (38) + BP(25) | - | - | Logistic regression without covariates | Nagelkerke's $R^2$(%)=0.184 , $p$=0.702 | Logistic regression with covariates (age, gender, and PCs) | Nagelkerke's $R^2$(PRS) (%)=0.579 , $p$(PRS)=0.448 |
| | | | | $p_T$<0.001 | | | | | Nagelkerke's $R^2$(%)=0.422 , $p$=0.562 | | Nagelkerke's $R^2$(PRS) (%)=0.231 , $p$(PRS)=0.630 |
| | | | | $p_T$<0.01 | | | | | Nagelkerke's $R^2$(%)=0.004, $p$=0.957 | | Nagelkerke's $R^2$(PRS) (%)=0.0001, $p$(PRS)=0.992 |
| | | | | $p_T$<0.05 | | | | | Nagelkerke's $R^2$(%)=0.008 , $p$=0.936 | | Nagelkerke's $R^2$(PRS) (%)=0.111 , $p$(PRS)=0.738 |
| | | | | $p_T$<0.1 | | | | | Nagelkerke's $R^2$(%)=0.092 , $p$=0.787 | | Nagelkerke's $R^2$(PRS) (%)=0.006 , $p$(PRS)=0.937 |
| | | | | $p_T$<0.2 | | | | | Nagelkerke's $R^2$(%)=0.206 , $p$=0.686 | | Nagelkerke's $R^2$(PRS) (%)=0.007 , $p$(PRS)=0.935 |
| | | | | $p_T$<0.3 | | | | | Nagelkerke's $R^2$(%)=0.046 , $p$=0.849 | | Nagelkerke's $R^2$(PRS) (%)=0.053 , $p$(PRS)=0.818 |
| | | | | $p_T$<0.4 | | | | | Nagelkerke's $R^2$(%)=0.016 , $p$=0.909 | | Nagelkerke's $R^2$(PRS) (%)=0.038 , $p$(PRS)=0.845 |
| | | | | $p_T$<0.5 | | | | | Nagelkerke's $R^2$(%)=0.055 , $p$=0.835 | | Nagelkerke's $R^2$(PRS) (%)=0.004 , $p$(PRS)=0.949 |
| | | | | $p_T$<1 | | | | | Nagelkerke's $R^2$(%)=0.043 , $p$=0.854 | | Nagelkerke's $R^2$(PRS) (%)=0.003 , $p$(PRS)=0.955 |
| 2C | Association analysis between miR-124 and PRSs (at the $p$-value thresholds where the PRSs best account for case-control status) | $R^2$ | SZ+BP PRS | | CON (25)<br>Case (32) = SZ (16) + BP(16) | - | - | Linear regression without covariates | $R^2$(%)=17.4 , $p$=0.0012 | Linear regression with covariates (age, gender, PCs, and diagnosis) | $\beta_{standardized}$ (PRS)=0.291 , $p$(PRS)=0.037 |
| | | | SZ PRS | | | | | | $R^2$(%)=8.3 , $p$=0.02948 | | $\beta_{standardized}$ (PRS)=0.162 , $p$(PRS)=0.245 |
| | | | BP PRS | | | | | | $R^2$(%)=3.5 , $p$=0.16194 | | $\beta_{standardized}$ (PRS)=0.082238 , $p$(PRS)=0.555997 |
| | | | SZ vs. BP PRS | | | | | | $R^2$(%)=0 , $p$=0.98938 | | $\beta_{standardized}$ (PRS)=-0.001 , $p$(PRS)=0.991058 |
| 2D | Mediation analysis | Proportion mediated | SZ+BP PRS | - | CON (25)<br>Case (32) = SZ (16) + BP(16) | - | - | Mediation analysis with the covariates of | A bootstrapping method with 5,000 iterations, Proportion mediated (%) = 30.681, 95% CI [0.00960, 1.44], $p$=0.042 | - | - |
| | | Proportion mediated | SZ PRS | | | | | Mediation analysis | A bootstrapping method with 5,000 iterations, Proportion mediated (%) = 27.41, 95% CI [-0.960, 2.38], $p$=0.210 | - | - |
| | | Proportion mediated | BP PRS | | | | | Mediation analysis | A bootstrapping method with 5,000 iterations, Proportion mediated (%) = 0.0986, 95% CI [-0.5025, 0.79], $p$=0.7120 | - | - |
| | | Proportion mediated | SZ vs. BP PRS | | | | | Mediation analysis | A bootstrapping method with 5,000 iterations, Proportion mediated (%) = 1.149, 95% CI [-5.31, 6.01], $p$=0.88 | - | - |

#### Figure 3

| Figure | Test | Measurement | Group | Sub-group | Sample size | Average | Standard deviation | Statistical test | Statistical result | *post hoc* test | Result from the *post hoc* tests |
|---|---|---|---|---|---|---|---|---|---|---|---|
| 3B | qRT-PCR for miR-124 in the mPFC | Relative expression (A.U.) | CON | - | 7 | 1 | 0.394578 | - | - | Two-tailed Student's t-test | $t(12)$=-4.095273, $p$=0.001485 |
| | | | miR-124 | | 7 | 2.418784 | 0.827329 | | | | |
| 3C | Three-chmaber social interaction test (trial1): sociability test | Investigation time (s) | CON AAV | Non-social | 11 | 34.545 | 14.376 | Two-way Mixed ANOVA<br>Between-subjects factor: AAV (CON v.s miR-14)<br>Within-subjects factor: Chamber (non-social vs. social) | AAV: $F(1,19)$=0.939518, $p$=0.344580<br>Chamber: $F(1,19)$=45.602986, $p$=0.000002<br>AAV*Chamber: $F(1,19)$=21.518772, $p$=0.000104 | Two-tailed paried Student's t-test | $t(10)$=-7.950, $p$=0.000012 |
| | | | | Social | 11 | 91.636 | 29.090 | | | | |
| | | | miR-124 AAV | Non-social | 10 | 50.800 | 16.498 | | | Two-tailed paried Student's t-test | $t(9)$=-1.394, $p$=0.196646 |
| | | | | Social | 10 | 60.000 | 21.980 | | | | |
| | Three-chmaber social interaction test (trial2): social recognition test | Investigation time (s) | CON AAV | Familiar | 11 | 27.000 | 8.752 | Two-way Mixed ANOVA<br>between-subjects factor: AAV (CON v.s miR-14)<br>within-subjects factor: Chamber (familiar vs. novel) | AAV: $F(1,19)$=2.161648, $p$=0.157859<br>Chamber: $F(1,42)$=22.142131, $p$=0.000154<br>AAV*Chamber: $F(1,42)$=6.991824, $p$=0.015999 | Two-tailed paried Student's t-test | $t(10)$=5.251, $p$=0.000373 |
| | | | | Novel | 11 | 54.455 | 18.886 | | | | |
| | | | miR-124 AAV | Familiar | 10 | 45.000 | 15.741 | | | Two-tailed paried Student's t-test | $t(9)$=1.447, $p$=0.181781 |
| | | | | Novel | 10 | 52.700 | 15.952 | | | | |
| 3D | Amphetamine challenge | Locomotion activity (counts / 5 min)<br>- left | Habituation | CON_1 | 14 | 994.143 | 183.180 | Two-way Mixed ANOVA<br>Between-subjects factor: AAV (CON vs. miR-124)<br>Within-subjects factor: Time (multiple time points) | AAV: F(1,26)=0.281, p=0.601<br>Time: F(5.551,286)=80.016, p=6.878E-43<br>AAV*Time: F(5.551,286)=0.718, p=0.625 | Two-tailed Student's t-test | $t(26)$=0.231, $p$=0.819492 |
| | | | | miR-124_1 | 14 | 981.571 | 89.901 | | | | |
| | | | | CON_2 | 14 | 876.786 | 142.009 | | | Two-tailed Student's t-test | $t(26)$=0.547, $p$=0.589125 |
| | | | | miR-124_2 | 14 | 848.786 | 128.581 | | | | |
| | | | | CON_3 | 14 | 715.786 | 105.569 | | | Two-tailed Student's t-test | $t(26)$=-0.200, $p$=0.843377 |
| | | | | miR-124_3 | 14 | 724.500 | 124.703 | | | | |
| | | | | CON_4 | 14 | 718.357 | 101.411 | | | Two-tailed Student's t-test | $t(26)$=0.557, $p$=0.582327 |
| | | | | miR-124_4 | 14 | 688.000 | 176.944 | | | | |
| | | | | CON_5 | 14 | 644.286 | 117.786 | | | Two-tailed Student's t-test | $t(26)$=-0.095, $p$=0.924650 |
| | | | | miR-124_5 | 14 | 648.214 | 99.087 | | | | |
| | | | | CON_6 | 14 | 602.857 | 123.622 | | | Two-tailed Student's t-test | $t(26)$=-0.604, $p$=0.551209 |
| | | | | miR-124_6 | 14 | 632.714 | 137.659 | | | | |
| | | | | CON_7 | 14 | 518.143 | 201.558 | | | Two-tailed Student's t-test | $t(26)$=-0.729, $p$=0.472483 |
| | | | | miR-124_7 | 14 | 568.071 | 158.223 | | | | |
| | | | | CON_8 | 14 | 450.286 | 159.635 | | | Two-tailed Student's t-test | $t(26)$=-1.000, $p$=0.326675 |
| | | | | miR-124_8 | 14 | 505.357 | 130.396 | | | | |
| | | | | CON_9 | 14 | 409.786 | 205.144 | | | Two-tailed Student's t-test | $t(26)$=-1.023, $p$=0.315823 |
| | | | | miR-124_9 | 14 | 486.357 | 190.738 | | | | |
| | | | | CON_10 | 14 | 423.500 | 162.295 | | | Two-tailed Student's t-test | $t(26)$=-0.304, $p$=0.763542 |
| | | | | miR-124_10 | 14 | 443.357 | 182.730 | | | | |
| | | | | CON_11 | 14 | 392.286 | 179.039 | | | Two-tailed Student's t-test | $t(26)$=-1.112, $p$=0.276471 |
| | | | | miR-124_11 | 14 | 463.429 | 159.018 | | | | |
| | | | | CON_12 | 14 | 361.857 | 196.690 | | | Two-tailed Student's t-test | $t(26)$=-0.292, $p$=0.772718 |
| | | | | miR-124_12 | 14 | 380.500 | 135.792 | | | | |
| | | | Saline | CON_1 | 14 | 382.714 | 205.590 | Two-way Mixed ANOVA<br>Between-subjects factor: AAV (CON vs. miR-124)<br>Within-subjects factor: Time (multiple time points) | AAV: $F(1,26)$=1.240, $p$=0.276<br>Time: $F(6.626,286)$=4.672, $p$=1.128E-04<br>AAV*Time: $F(6.626,286)$=0.999, $p$=0.432 | Two-tailed Student's t-test | $t(26)$=-0.592, $p$=0.558816 |
| | | | | miR-124_1 | 14 | 423.786 | 158.326 | | | | |
| | | | | CON_2 | 14 | 267.143 | 107.599 | | | Two-tailed Student's t-test | $t(26)$=-1.410, $p$=0.170348 |
| | | | | miR-124_2 | 14 | 344.500 | 174.795 | | | | |
| | | | | CON_3 | 14 | 185.214 | 85.474 | | | Two-tailed Student's t-test | $t(26)$=-2.749, $p$=0.010729 |
| | | | | miR-124_3 | 14 | 303.500 | 136.446 | | | | |
| | | | | CON_4 | 14 | 250.071 | 121.153 | | | Two-tailed Student's t-test | $t(26)$=1.038, $p$=0.308819 |
| | | | | miR-124_4 | 14 | 204.357 | 111.692 | | | | |
| | | | | CON_5 | 14 | 260.143 | 121.454 | | | Two-tailed Student's t-test | $t(26)$=0.611, $p$=0.546528 |
| | | | | miR-124_5 | 14 | 226.357 | 167.513 | | | | |
| | | | | CON_6 | 14 | 215.643 | 135.977 | | | Two-tailed Student's t-test | $t(26)$=-0.972, $p$=0.340126 |
| | | | | miR-124_6 | 14 | 279.500 | 204.854 | | | | |
| | | | | CON_7 | 14 | 267.000 | 121.840 | | | Two-tailed Welch's t-test | $t(20.847)$=-0.705, $p$=0.488607 |
| | | | | miR-124_7 | 14 | 312.786 | 210.243 | | | | |
| | | | | CON_8 | 14 | 265.571 | 126.014 | | | Two-tailed Student's t-test | $t(26)$=-0.892, $p$=0.380418 |
| | | | | miR-124_8 | 14 | 316.429 | 172.046 | | | | |
| | | | | CON_9 | 14 | 209.357 | 89.828 | | | Two-tailed Student's t-test | $t(26)$=-0.565, $p$=0.576717 |
| | | | | miR-124_9 | 14 | 236.071 | 152.303 | | | | |
| | | | | CON_10 | 14 | 254.857 | 129.396 | | | Two-tailed Student's t-test | $t(26)$=-0.702, $p$=0.489218 |
| | | | | miR-124_10 | 14 | 301.071 | 209.801 | | | | |
| | | | | CON_11 | 14 | 251.929 | 121.731 | | | Two-tailed Student's t-test | $t(26)$=-0.916, $p$=0.368024 |
| | | | | miR-124_11 | 14 | 303.357 | 171.176 | | | | |
| | | | | CON_12 | 14 | 225.071 | 157.920 | | | Two-tailed Student's t-test | $t(26)$=-1.162, $p$=0.255942 |
| | | | | miR-124_12 | 14 | 293.357 | 153.103 | | | | |
| | | | AMPH | CON_1 | 14 | 486.857 | 237.405 | Two-way Mixed ANOVA with the Greenhouse-Geisser correction<br>Between-subjects factor: AAV (CON vs. miR-124)<br>Within-subjects factor: Time (multiple time points) | For the first 60 min,<br>AAV: $F(1,26)$=22.416, $p$=6.772E-05<br>Time: $F(2.545,66.164)$=16.127, $p$=2.626E-7<br>AAV*Time: $F(2.545,66.164)$=3.870, $p$=0.018<br><br>For the first 30 min,<br>AAV: $F(1,26)$=21.885, $p$=7.9E-8<br>Time: $F(2.226,57.866)$=23.612, $p$=1.086E-8<br>AAV*Time: $F(2.226,57.866)$=9.065, $p$=2.34E-4 | Two-tailed Student's t-test | $t(26)$=-2.137, $p$=0.042171 |
| | | | | miR-124_1 | 14 | 713.929 | 318.907 | | | | |
| | | | | CON_2 | 14 | 499.071 | 145.360 | | | Two-tailed Welch's t-test | $t(15.927)$=-4.577, $p$=0.000314 |
| | | | | miR-124_2 | 14 | 1055.071 | 430.667 | | | | |
| | | | | CON_3 | 14 | 570.929 | 176.891 | | | Two-tailed Welch's t-test | $t(16.579)$=-4.716, $p$=0.000213 |
| | | | | miR-124_3 | 14 | 1206.429 | 472.136 | | | | |
| | | | | CON_4 | 14 | 652.286 | 214.525 | | | Two-tailed Welch's t-test | $t(16.984)$=-3.843, $p$=0.001306 |
| | | | | miR-124_4 | 14 | 1250.357 | 541.401 | | | | |
| | | | | CON_5 | 14 | 673.500 | 221.443 | | | Two-tailed Welch's t-test | $t(18.538)$=-4.638, $p$=0.000190 |
| | | | | miR-124_5 | 14 | 1315.500 | 468.222 | | | | |
| | | | | CON_6 | 14 | 553.786 | 170.899 | | | Two-tailed Welch's t-test | $t(16.064)$=-5.438, $p$=0.000054 |
| | | | | miR-124_6 | 14 | 1314.000 | 494.341 | | | | |
| | | | | CON_7 | 14 | 517.571 | 166.176 | | | Two-tailed Welch's t-test | $t(15.685)$=-4.859, $p$=0.000184 |
| | | | | miR-124_7 | 14 | 1219.429 | 514.286 | | | | |
| | | | | CON_8 | 14 | 485.571 | 113.713 | | | Two-tailed Welch's t-test | $t(14.498)$=-4.626, $p$=0.000359 |
| | | | | miR-124_8 | 14 | 1087.286 | 473.163 | | | | |
| | | | | CON_9 | 14 | 455.500 | 152.541 | | | Two-tailed Welch's t-test | $t(15.201)$=-3.360, $p$=0.004231 |
| | | | | miR-124_9 | 14 | 944.143 | 522.367 | | | | |
| | | | | CON_10 | 14 | 393.500 | 188.082 | | | Two-tailed Welch's t-test | $t(16.186)$=-2.911, $p$=0.010104 |
| | | | | miR-124_10 | 14 | 833.429 | 533.187 | | | | |
| | | | | CON_11 | 14 | 385.286 | 175.292 | | | Two-tailed Welch's t-test | $t(16.017)$=-2.381, $p$=0.029993 |
| | | | | miR-124_11 | 14 | 729.143 | 511.036 | | | | |
| | | | | CON_12 | 14 | 344.500 | 174.904 | | | Two-tailed Welch's t-test | $t(17.401)$=-2.708, $p$=0.014703 |
| | | | | miR-124_12 | 14 | 673.000 | 418.811 | | | | |
| 3E | mEPSC recordings | mEPSC amplitdue (pA) | CON | - | 11 | 9.718 | 1.0643 | - | - | Two-tailed Student's t-test | $t(19)$=-3.590, $p$=0.002 |
| | | | miR-124 | | 10 | 11.956 | 1.7431 | | | | |
| | | mEPSC frequency (Hz) | CON | - | 11 | 1.734 | 0.3974 | - | - | Two-tailed Student's t-test | $t(19)$=-0.787, $p$=0.441 |
| | | | miR-124 | | 10 | 1.869 | 0.3895 | | | | |

#### Figure 4

| Figure | Test | Measurement | Group | Sub-group | Sample size | Average | Standard deviation | Statistical test | Statistical result | *post hoc* test | Result from the *post hoc* tests |
|---|---|---|---|---|---|---|---|---|---|---|---|
| 4A | mEPSC recordings with or without Naspm | mESPC amplitude | CON AAV | without Naspm | 10 | 10.582 | 1.473 | Two-way Mixed ANOVA<br>Between-subjects factor: AAV<br>Within-subjects factor: Naspm (without Naspm vs. with Naspm) | AAV: $F(1,18)=2.646, p=0.121$<br>Naspm: $F(1,18)=15.182, p=0.001$<br>AAV*Naspm: $F(1,18)=8.454, p=0.009$ | Two-tailed paried Student's t-test | $t(9)=-0.840, p=0.422645$ |
| | | | | with Naspm | 10 | 10.230 | 1.160 | | | | |
| | | | miR-124 AAV | without Naspm | 10 | 12.634 | 2.379 | | | Two-tailed paried Student's t-test | $t(9)=-4.208, p=0.002280$ |
| | | | | with Naspm | 10 | 10.212 | 1.080 | | | | |
| | | mESPC frequency | CON AAV | without Naspm | 10 | 0.807 | 0.475 | Two-way Mixed ANOVA<br>Between-subjects factor: AAV<br>Within-subjects factor: Naspm (without Naspm vs. with Naspm) | AAV: $F(1,18)=0.017, p=0.896$<br>Naspm: $F(1,18)=0.400, p=0.535$<br>AAV*Naspm: $F(1,18)=0.593, p=0.451$ | Two-tailed paried Student's t-test | $t(9)=-0.099, p=0.923338$ |
| | | | | with Naspm | 10 | 0.802 | 0.454 | | | | |
| | | | miR-124 AAV | without Naspm | 10 | 0.751 | 0.380 | | | Two-tailed paried Student's t-test | $t(9)=0.975, p=0.354930$ |
| | | | | with Naspm | 10 | 0.806 | 0.455 | | | | |
| 4B | Three-chmaber social interaction test (trial1): sociability test | Investigation time (s) | CON AAV + Saline | Non-social | 9 | 33.667 | 6.652 | Two-way Mixed ANOVA<br>Between-subjects factor: AAV (CON vs. miR-124)<br>Within-subjects factor: Chamber (Non-social vs. Social) | AAV: $F(1,17)=1.469, p=1.065E-04$<br>Chamber: $F(1,17)=25.138, p=0.535$<br>AAV*Chamber: $F(1,17)=14.684, p=0.001$ | Two-tailed paried Student's t-test | $t(8)=-4.668, p=0.001606$ |
| | | | | Social | 9 | 87.556 | 31.679 | | | | |
| | | | miR-124 AAV +Saline | Non-social | 10 | 48.200 | 16.772 | | | Two-tailed paried Student's t-test | $t(9)=-1.406, p=0.193251$ |
| | | | | Social | 10 | 55.400 | 19.929 | | | | |
| | | Investigation time (s) | miR-124 AAV +Saline | Non-social | 10 | 48.200 | 16.772 | Two-way Mixed ANOVA<br>Between-subjects factor: Intervention (Saline vs. Naspm)<br>Within-subjects factor: Chamber (Non-social vs. Social) | Intervention: $F(1,18)=0.207, p=0.655$<br>Chamber: $F(1,18)=13.310, p=0.002$<br>Intervention*Chamber: $F(1,18)=6.725, p=0.018$ | Two-tailed paried Student's t-test | $t(9)=-1.406, p=0.193251$ |
| | | | | Social | 10 | 55.400 | 19.929 | | | | |
| | | | miR-124 AAV + Naspm | Non-social | 10 | 26.600 | 11.815 | | | Two-tailed paried Student's t-test | $t(9)=-3.367, p=0.008302$ |
| | | | | Social | 10 | 69.200 | 39.805 | | | | |
| | Three-chmaber social interaction test (trial2): social novelty recognition | Investigation time (s) | CON AAV + Saline | Familiar | 9 | 29.111 | 17.230 | Two-way Mixed ANOVA<br>Between-subjects factor: AAV (CON vs. miR-124)<br>Within-subjects factor: Chamber (Familiar vs. Novel) | AAV: $F(1,17)=2.952E-05, p=0.996$<br>Chamber: $F(1,17)=21.377, p=2.426E-04$<br>AAV*Chamber: $F(1,17)=15.515, p=0.001$ | Two-tailed paried Student's t-test | $t(8)=4.705, p=0.001532$ |
| | | | | Novel | 9 | 72.889 | 27.634 | | | | |
| | | | miR-124 AAV +Saline | Familiar | 10 | 49.200 | 23.203 | | | Two-tailed paried Student's t-test | $t(9)=0.710, p=0.495633$ |
| | | | | Novel | 10 | 52.700 | 22.416 | | | | |
| | | Investigation time (s) | miR-124 AAV +Saline | Familiar | 10 | 49.200 | 23.203 | Two-way Mixed ANOVA<br>Between-subjects factor: Intervention (Saline vs. Naspm)<br>Within-subjects factor: Chamber (Familiar vs. Novel) | Intervention: $F(1,18)=0.680, p=0.420$<br>Chamber: $F(1,18)=16.490, p=0.001$<br>Intervention*Chamber: $F(1,18)=9.276, p=0.007$ | Two-tailed paried Student's t-test | $t(9)=0.710, p=0.495633$ |
| | | | | Novel | 10 | 52.700 | 22.416 | | | | |
| | | | miR-124 AAV + Naspm | Familiar | 10 | 31.300 | 17.017 | | | Two-tailed paried Student's t-test | $t(9)=5.081, p=0.000662$ |
| | | | | Novel | 10 | 55.800 | 22.739 | | | | |
| 4C | Amphetamine challenge | Locomotion activity (counts / 5 min) - left | Habituation:<br>CON AAV vs. miR-124 AAV<br>(no Saline or Naspm injections yet) | CON_1 | 12 | 1009.583 | 191.767 | Two-way Mixed ANOVA<br>Between-subjects factor: AAV (CON vs. miR-124)<br>Within-subjects factor: Time (multiple time points) | AAV: $F(1,33)=0.270, p=0.607$<br>Time: $F(5.450,363)=75.283, p=3.229E-44$<br>AAV*Time: $F(5.450,363)=0.432, p=0.841$ | Two-tailed Student's t-test | $t(33)=-0.191, p=0.849831$ |
| | | | | miR-124_1 | 23 | 1023.522 | 211.473 | | | | |
| | | | | CON_2 | 12 | 894.583 | 170.754 | | | Two-tailed Student's t-test | $t(33)=0.299, p=0.766900$ |
| | | | | miR-124_2 | 23 | 876.522 | 169.163 | | | | |
| | | | | CON_3 | 12 | 798.833 | 180.241 | | | Two-tailed Student's t-test | $t(33)=0.530, p=0.599980$ |
| | | | | miR-124_3 | 23 | 767.348 | 159.921 | | | | |
| | | | | CON_4 | 12 | 781.333 | 181.600 | | | Two-tailed Student's t-test | $t(33)=0.932, p=0.358299$ |
| | | | | miR-124_4 | 23 | 722.174 | 176.660 | | | | |
| | | | | CON_5 | 12 | 688.750 | 141.504 | | | Two-tailed Student's t-test | $t(33)=-0.044, p=0.965082$ |
| | | | | miR-124_5 | 23 | 691.217 | 164.313 | | | | |
| | | | | CON_6 | 12 | 727.500 | 157.761 | | | Two-tailed Student's t-test | $t(33)=-1.431, p=0.161845$ |
| | | | | miR-124_6 | 23 | 645.261 | 163.172 | | | | |
| | | | | CON_7 | 12 | 585.917 | 246.593 | | | Two-tailed Student's t-test | $t(33)=0.668, p=0.508677$ |
| | | | | miR-124_7 | 23 | 535.565 | 191.751 | | | | |
| | | | | CON_8 | 12 | 568.917 | 223.097 | | | Two-tailed Student's t-test | $t(33)=0.548, p=0.587696$ |
| | | | | miR-124_8 | 23 | 529.652 | 189.585 | | | | |
| | | | | CON_9 | 12 | 536.750 | 249.022 | | | Two-tailed Student's t-test | $t(33)=0.182, p=0.856764$ |
| | | | | miR-124_9 | 23 | 521.565 | 226.742 | | | | |
| | | | | CON_10 | 12 | 497.833 | 199.172 | | | Two-tailed Student's t-test | $t(33)=0.077, p=0.938984$ |
| | | | | miR-124_10 | 23 | 492.174 | 209.393 | | | | |
| | | | | CON_11 | 12 | 473.250 | 185.994 | | | Two-tailed Student's t-test | $t(33)=0.298, p=0.767663$ |
| | | | | miR-124_11 | 23 | 452.957 | 193.913 | | | | |
| | | | | CON_12 | 12 | 460.333 | 209.109 | | | Two-tailed Student's t-test | $t(33)=0.570, p=0.572348$ |
| | | | | miR-124_12 | 23 | 421.870 | 178.734 | | | | |
| | | | Saline:<br>CON AAV vs. miR-124 AAV<br>(no Saline or Naspm injections yet) | CON_1 | 12 | 409.583 | 143.893 | Two-way Mixed ANOVA<br>Between-subjects factor: AAV (CON vs. miR-124)<br>Within-subjects factor: Time (multiple time points) | AAV: $F(1,33)=0.880, p=0.355$<br>Time: $F(6.988,363)=4.246, p=1.998E-04$<br>AAV*Time: $F(6.988,363)=0.584, p=0.769$ | Two-tailed Student's t-test | $t(33)=-0.623, p=0.537290$ |
| | | | | miR-124_1 | 23 | 452.957 | 216.567 | | | | |
| | | | | CON_2 | 12 | 314.500 | 76.068 | | | Two-tailed Welch's t-test | $t(31.234)=-1.447, p=0.157901$ |
| | | | | miR-124_2 | 23 | 382.043 | 197.570 | | | | |
| | | | | CON_3 | 12 | 309.917 | 113.582 | | | Two-tailed Student's t-test | $t(33)=0.971, p=0.338545$ |
| | | | | miR-124_3 | 23 | 263.304 | 144.222 | | | | |
| | | | | CON_4 | 12 | 275.583 | 82.502 | | | Two-tailed Welch's t-test | $t(32.515)=-0.675, p=0.504141$ |
| | | | | miR-124_4 | 23 | 306.217 | 185.090 | | | | |
| | | | | CON_5 | 12 | 282.250 | 132.924 | | | Two-tailed Welch's t-test | $t(30.691)=0.041, p=0.967835$ |
| | | | | miR-124_5 | 23 | 279.957 | 198.332 | | | | |
| | | | | CON_6 | 12 | 252.583 | 124.260 | | | Two-tailed Welch's t-test | $t(32.567)=-1.027, p=0.312069$ |
| | | | | miR-124_6 | 23 | 311.739 | 216.197 | | | | |
| | | | | CON_7 | 12 | 271.167 | 156.883 | | | Two-tailed Student's t-test | $t(33)=-0.399, p=0.692744$ |
| | | | | miR-124_7 | 23 | 298.652 | 209.600 | | | | |
| | | | | CON_8 | 12 | 267.917 | 152.005 | | | Two-tailed Student's t-test | $t(33)=-1.040, p=0.306055$ |
| | | | | miR-124_8 | 23 | 333.478 | 188.375 | | | | |
| | | | | CON_9 | 12 | 224.833 | 57.157 | | | Two-tailed Welch's t-test | $t(28.709)=-1.165, p=0.253423$ |
| | | | | miR-124_9 | 23 | 274.304 | 187.562 | | | | |
| | | | | CON_10 | 12 | 267.750 | 157.997 | | | Two-tailed Student's t-test | $t(33)=-0.777, p=0.442716$ |
| | | | | miR-124_10 | 23 | 319.348 | 199.212 | | | | |
| | | | | CON_11 | 12 | 255.083 | 147.150 | | | Two-tailed Student's t-test | $t(33)=-0.636, p=0.529255$ |
| | | | | miR-124_11 | 23 | 289.304 | 153.083 | | | | |
| | | | | CON_12 | 12 | 219.000 | 131.015 | | | Two-tailed Student's t-test | $t(33)=-1.604, p=0.118147$ |
| | | | | miR-124_12 | 23 | 307.913 | 166.564 | | | | |
| | | | AMPH<br>CON AAV + Saline<br>vs. miR-124 AAV +Saline<br>(Saline or Naspm injected) | CON_1 | 12 | 497.167 | 182.496 | Two-way Mixed ANOVA<br>Between-subjects factor: AAV (CON vs. miR-124)<br>Within-subjects factor: Time (multiple time points) | AAV: $F(1,23)=15.361, p=0.001$<br>Time: $F(2.583,253)=6.054, p=0.002$<br>AAV*Time: $F(2.583,253)=3.396, p=0.029$ | Two-tailed Welch's t-test | $t(18.378)=-1.559, p=0.136114$ |
| | | | | miR-124_1 | 13 | 669.231 | 349.757 | | | | |
| | | | | CON_2 | 12 | 446.750 | 255.970 | | | Two-tailed Student's t-test | $t(23)=-3.918, p=0.000689$ |
| | | | | miR-124_2 | 13 | 991.846 | 414.026 | | | | |
| | | | | CON_3 | 12 | 486.750 | 310.776 | | | Two-tailed Student's t-test | $t(23)=-4.667, p=0.000107$ |
| | | | | miR-124_3 | 13 | 1149.615 | 390.874 | | | | |
| | | | | CON_4 | 12 | 604.500 | 301.384 | | | Two-tailed Student's t-test | $t(23)=-3.707, p=0.001160$ |
| | | | | miR-124_4 | 13 | 1154.077 | 423.748 | | | | |
| | | | | CON_5 | 12 | 542.250 | 333.947 | | | Two-tailed Student's t-test | $t(23)=-4.214, p=0.000330$ |
| | | | | miR-124_5 | 13 | 1166.923 | 400.733 | | | | |
| | | | | CON_6 | 12 | 524.000 | 327.969 | | | Two-tailed Student's t-test | $t(23)=-4.311, p=0.000259$ |
| | | | | miR-124_6 | 13 | 1158.692 | 400.822 | | | | |
| | | | | CON_7 | 12 | 506.167 | 316.325 | | | Two-tailed Student's t-test | $t(23)=-3.453, p=0.002159$ |
| | | | | miR-124_7 | 13 | 1040.308 | 440.896 | | | | |
| | | | | CON_8 | 12 | 482.083 | 284.678 | | | Two-tailed Student's t-test | $t(23)=-3.390, p=0.002518$ |
| | | | | miR-124_8 | 13 | 968.154 | 414.216 | | | | |
| | | | | CON_9 | 12 | 503.500 | 335.109 | | | Two-tailed Student's t-test | $t(23)=-2.220, p=0.036539$ |
| | | | | miR-124_9 | 13 | 853.769 | 441.291 | | | | |
| | | | | CON_10 | 12 | 472.583 | 295.770 | | | Two-tailed Student's t-test | $t(23)=-2.240, p=0.035068$ |
| | | | | miR-124_10 | 13 | 827.000 | 468.286 | | | | |
| | | | | CON_11 | 12 | 449.417 | 305.212 | | | Two-tailed Student's t-test | $t(23)=-1.631, p=0.116452$ |
| | | | | miR-124_11 | 13 | 693.231 | 426.362 | | | | |
| | | | | CON_12 | 12 | 439.750 | 273.855 | | | Two-tailed Student's t-test | $t(23)=-1.443, p=0.162374$ |
| | | | | miR-124_12 | 13 | 645.154 | 416.454 | | | | |

| | | | | | | | | | | | |
|---|---|---|---|---|---|---|---|---|---|---|---|
| | | | AMPH<br>miR-124 AAV +Saline<br>vs. miR-124 AAV + Naspm<br>(Saline or Naspm injected) | Saline1 | 10 | 693.600 | 398.569 | Two-way Mixed ANOVA with the Greenhouse-Geisser correction<br>Between-subjects factor: Intervention (Saline vs. Naspm)<br>Within-subjects factor: Time (multiple time points) | For the first 60 min,<br>Intervention: $F(1,21)=7.733, p=0.011$<br>Time: $F(2.414,50.696)=6.770, p=0.001$<br>Intervention*Time: $F(2.414,50.696)=1.775, p=0.059$<br><br>For the first 15 min,<br>Intervention: $F(1,21)=2.570, p=0.124$<br>Time: $F(1.542,32.391)=13.275, p=1.92E-04$<br>Intervention*Time: $F(1.542,32.391)=9.671, p=0.001$ | Two-tailed Student's t-test | $t(21)=-0.156, p=0.877548$ |
| | | | | Naspm1 | 13 | 669.231 | 349.757 | | | | |
| | | | | Saline2 | 10 | 724.000 | 242.967 | | | Two-tailed Student's t-test | $t(21)=1.814, p=0.084021$ |
| | | | | Naspm2 | 13 | 991.846 | 414.026 | | | | |
| | | | | Saline3 | 10 | 730.300 | 277.123 | | | Two-tailed Student's t-test | $t(21)=2.875, p=0.009059$ |
| | | | | Naspm3 | 13 | 1149.615 | 390.874 | | | | |
| | | | | Saline4 | 10 | 728.000 | 370.258 | | | Two-tailed Student's t-test | $t(21)=2.522, p=0.019824$ |
| | | | | Naspm4 | 13 | 1154.077 | 423.748 | | | | |
| | | | | Saline5 | 10 | 799.900 | 371.089 | | | Two-tailed Student's t-test | $t(21)=2.247, p=0.035508$ |
| | | | | Naspm5 | 13 | 1166.923 | 400.733 | | | | |
| | | | | Saline6 | 10 | 685.900 | 214.822 | | | Two-tailed Welch's t-test | $t(19.087)=-3.629, p=0.001776$ |
| | | | | Naspm6 | 13 | 1158.692 | 400.822 | | | | |
| | | | | Saline7 | 10 | 630.500 | 269.123 | | | Two-tailed Welch's t-test | $t(20.140)=2.751, p=0.012276$ |
| | | | | Naspm7 | 13 | 1040.308 | 440.896 | | | | |
| | | | | Saline8 | 10 | 549.900 | 220.292 | | | Two-tailed Welch's t-test | $t(19.019)=3.113, p=0.005721$ |
| | | | | Naspm8 | 13 | 968.154 | 414.216 | | | | |
| | | | | Saline9 | 10 | 542.300 | 232.973 | | | Two-tailed Welch's t-test | $t(18.954)=2.751, p=0.042045$ |
| | | | | Naspm9 | 13 | 853.769 | 441.291 | | | | |
| | | | | Saline10 | 10 | 518.700 | 214.264 | | | Two-tailed Welch's t-test | $t(17.675)=2.105, p=0.049914$ |
| | | | | Naspm10 | 13 | 827.000 | 468.286 | | | | |
| | | | | Saline11 | 10 | 514.700 | 184.842 | | | Two-tailed Welch's t-test | $t(17.211)=1.353, p=0.193427$ |
| | | | | Naspm11 | 13 | 693.231 | 426.362 | | | | |
| | | | | Saline12 | 10 | 499.400 | 207.125 | | | Two-tailed Welch's t-test | $t(18.419)=1.098, p=0.286483$ |
| | | | | Naspm12 | 13 | 645.154 | 416.454 | | | | |
| 4D | Three-chmaber<br>social interaction test<br>(trial1): sociability test | Investigation time (s) | CON AAV + mCherry AAV | Non-social | 10 | 30.900 | 12.827 | Two-way Mixed ANOVA<br>Between-subjects factor: AAV (CON vs. miR-124)<br>Within-subjects factor: Chamber (Non-social vs. Social) | AAV: $F(1,19)=1.294, p=0.269$<br>Chamber: $F(1,19)=23.325, p=1.165E-04$<br>AAV*Chamber: $F(1,17)=15.679, p=0.001$ | Two-tailed paried Student's t-test | $t(9)=-5.100, p=0.000645$ |
| | | | | Social | 10 | 91.700 | 30.667 | | | | |
| | | | miR-124 AAV + mCherry AAV | Non-social | 11 | 51.909 | 9.439 | | | Two-tailed paried Student's t-test | $t(10)=-0.796, p=0.444452$ |
| | | | | Social | 11 | 57.909 | 22.282 | | | | |
| | | Investigation time (s) | miR-124 AAV + mCherry AAV | Non-social | 11 | 51.909 | 9.439 | Two-way Mixed ANOVA<br>Between-subjects factor: Intervention (mCherry vs. GluA2)<br>Within-subjects factor: Chamber (Non-social vs. Social) | Intervention: $F(1,19)=1.288, p=0.271$<br>Chamber: $F(1,19)=10.055, p=0.005$<br>Intervention*Chamber: $F(1,19)=5.741, p=0.027$ | Two-tailed paried Student's t-test | $t(10)=-0.796, p=0.444452$ |
| | | | | Social | 11 | 57.909 | 22.282 | | | | |
| | | | miR-124 AAV + GluA2 AAV | Non-social | 10 | 40.900 | 20.366 | | | Two-tailed paried Student's t-test | $t(9)=-3.081, p=0.013124$ |
| | | | | Social | 10 | 84.000 | 35.169 | | | | |
| | Three-chmaber<br>social interaction test<br>(trial2): social novelty recognition | Investigation time (s) | CON AAV + mCherry | Familiar | 10 | 36.300 | 19.517 | Two-way Mixed ANOVA<br>Between-subjects factor: AAV (CON vs. miR-124)<br>Within-subjects factor: Chamber (Familiar vs. Novel) | AAV: $F(1,19)=0.782, p=0.388$<br>Chamber: $F(1,19)=14.314, p=0.001$<br>AAV*Chamber: $F(1,19)=4.662, p=0.044$ | Two-tailed paried Student's t-test | $t(9)=3.553, p=0.006187$ |
| | | | | Novel | 10 | 81.200 | 39.471 | | | | |
| | | | miR-124 AAV + mCherry | Familiar | 11 | 44.546 | 22.487 | | | Two-tailed paried Student's t-test | $t(10)=1.408, p=0.189374$ |
| | | | | Novel | 11 | 56.818 | 23.297 | | | | |
| | | Investigation time (s) | miR-124 AAV + mCherry | Familiar | 11 | 44.546 | 22.487 | Two-way Mixed ANOVA<br>Between-subjects factor: Intervention (mCherry vs. GluA2)<br>Within-subjects factor: Chamber (Familiar vs. Novel) | Intervention: $F(1,19)=0.117, p=0.736$<br>Chamber: $F(1,19)=10.016, p=0.005$<br>Intervention*Chamber: $F(1,19)=1.876, p=0.187$ | Two-tailed paried Student's t-test | $t(10)=1.408, p=0.189374$ |
| | | | | Novel | 11 | 56.818 | 23.297 | | | | |
| | | | miR-124 AAV + GluA2 | Familiar | 10 | 32.500 | 17.871 | | | Two-tailed paried Student's t-test | $t(9)=2.905, p=0.017466$ |
| | | | | Novel | 10 | 63.500 | 30.123 | | | | |
| | | | Habituation:<br>CON AAV + mCherry AAV vs.<br>miR-124 AAV + mCherry AAV | CON_1 | 11 | 648.818 | 390.379 | Two-way Mixed ANOVA<br>Between-subjects factor: AAV (CON vs. miR-124)<br>Within-subjects factor: Time (multiple time points) | AAV: $F(1,19)=0.557, p=0.465$<br>Time: $F(4.691,209)=31.475, p=9.508E-18$<br>AAV*Time: $F(4.691,209)=0.903, p=0.478$ | Two-tailed Student's t-test | $t(19)=-0.667, p=0.512725$ |
| | | | | miR-124_1 | 10 | 754.200 | 326.524 | | | | |
| | | | | CON_2 | 11 | 578.455 | 296.191 | | | Two-tailed Student's t-test | $t(19)=-0.477, p=0.639071$ |
| | | | | miR-124_2 | 10 | 636.700 | 260.134 | | | | |
| | | | | CON_3 | 11 | 453.727 | 251.679 | | | Two-tailed Student's t-test | $t(19)=-1.234, p=0.232367$ |
| | | | | miR-124_3 | 10 | 591.500 | 259.871 | | | | |
| | | | | CON_4 | 11 | 413.182 | 193.363 | | | Two-tailed Student's t-test | $t(19)=-1.176, p=0.254261$ |
| | | | | miR-124_4 | 10 | 522.500 | 232.530 | | | | |
| | | | | CON_5 | 11 | 348.909 | 204.781 | | | Two-tailed Student's t-test | $t(19)=-1.104, p=0.283308$ |
| | | | | miR-124_5 | 10 | 446.700 | 200.359 | | | | |
| | | | | CON_6 | 11 | 359.091 | 232.290 | | | Two-tailed Student's t-test | $t(19)=-0.531, p=0.601272$ |
| | | | | miR-124_6 | 10 | 414.500 | 245.468 | | | | |
| | | | | CON_7 | 11 | 328.636 | 251.561 | | | Two-tailed Student's t-test | $t(19)=-0.089, p=0.929750$ |
| | | | | miR-124_7 | 10 | 337.100 | 170.119 | | | | |
| | | | | CON_8 | 11 | 333.636 | 234.380 | | | Two-tailed Student's t-test | $t(19)=-0.671, p=0.510412$ |
| | | | | miR-124_8 | 10 | 396.700 | 191.559 | | | | |
| | | | | CON_9 | 11 | 332.182 | 223.515 | | | Two-tailed Student's t-test | $t(19)=0.164, p=0.871081$ |
| | | | | miR-124_9 | 10 | 317.900 | 166.883 | | | | |
| | | | | CON_10 | 11 | 237.364 | 209.479 | | | Two-tailed Student's t-test | $t(19)=-1.243, p=0.228829$ |
| | | | | miR-124_10 | 10 | 359.800 | 241.779 | | | | |
| | | | | CON_11 | 11 | 256.182 | 179.503 | | | Two-tailed Student's t-test | $t(19)=-0.600, p=0.555450$ |
| | | | | miR-124_11 | 10 | 306.300 | 203.219 | | | | |
| | | | | CON_12 | 11 | 236.727 | 206.249 | | | Two-tailed Student's t-test | $t(19)=-0.383, p=0.705880$ |
| | | | | miR-124_12 | 10 | 268.400 | 168.254 | | | | |
| | | | Habituation:<br>miR-124 AAV + mCherry AAV<br>vs. miR-124 AAV + GluA2 AAV | GluA2_1 | 11 | 691.909 | 241.086 | Two-way Mixed ANOVA<br>Between-subjects factor: Intervention (mCherry vs. GluA2)<br>Within-subjects factor: Time (multiple time points) | Intervention: $F(1,19)=0.128, p=0.725$<br>Time: $F(5.819,209)=32.917, p=3.807E-22$<br>Intervention*Time: $F(5.819,209)=1.471, p=0.197$ | Two-tailed Student's t-test | $t(19)=-0.501, p=0.622381$ |
| | | | | mCherry_1 | 10 | 754.200 | 326.524 | | | | |
| | | | | GluA2_2 | 11 | 634.636 | 213.667 | | | Two-tailed Student's t-test | $t(19)=0.020, p=0.984296$ |
| | | | | mCherry_2 | 10 | 636.700 | 260.134 | | | | |
| | | | | GluA2_3 | 11 | 557.364 | 218.418 | | | Two-tailed Student's t-test | $t(19)=0.327, p=0.747273$ |
| | | | | mCherry_3 | 10 | 591.500 | 259.871 | | | | |
| | | | | GluA2_4 | 11 | 508.818 | 174.420 | | | Two-tailed Student's t-test | $t(19)=0.153, p=0.879636$ |
| | | | | mCherry_4 | 10 | 522.500 | 232.530 | | | | |
| | | | | GluA2_5 | 11 | 511.364 | 208.623 | | | Two-tailed Student's t-test | $t(19)=-0.723, p=0.478602$ |
| | | | | mCherry_5 | 10 | 446.700 | 200.359 | | | | |
| | | | | GluA2_6 | 11 | 474.091 | 155.830 | | | Two-tailed Student's t-test | $t(19)=-0.671, p=0.510342$ |
| | | | | mCherry_6 | 10 | 414.500 | 245.468 | | | | |
| | | | | GluA2_7 | 11 | 472.182 | 175.382 | | | Two-tailed Student's t-test | $t(19)=-1.788, p=0.089737$ |
| | | | | mCherry_7 | 10 | 337.100 | 170.119 | | | | |
| | | | | GluA2_8 | 11 | 421.455 | 193.360 | | | Two-tailed Student's t-test | $t(19)=-0.294, p=0.771719$ |
| | | | | mCherry_8 | 10 | 396.700 | 191.559 | | | | |
| | | | | GluA2_9 | 11 | 380.727 | 178.283 | | | Two-tailed Student's t-test | $t(19)=-0.831, p=0.416141$ |
| | | | | mCherry_9 | 10 | 317.900 | 166.883 | | | | |
| | | | | GluA2_10 | 11 | 359.273 | 170.485 | | | Two-tailed Student's t-test | $t(19)=0.006, p=0.995417$ |
| | | | | mCherry_10 | 10 | 359.800 | 241.779 | | | | |
| | | | | GluA2_11 | 11 | 393.000 | 135.943 | | | Two-tailed Student's t-test | $t(19)=-1.159, p=0.260633$ |
| | | | | mCherry_11 | 10 | 306.300 | 203.219 | | | | |
| | | | | GluA2_12 | 11 | 285.000 | 133.195 | | | Two-tailed Student's t-test | $t(19)=-0.252, p=0.803821$ |
| | | | | mCherry_12 | 10 | 268.400 | 168.254 | | | | |

| | | | | | | | | | | | |
|---|---|---|---|---|---|---|---|---|---|---|---|
| 4E | Amphetamine challenge | Locomotion activity (counts / 5 min) - left | Saline:<br>CON AAV + mCherry AAV vs. miR-124 AAV + mCherry AAV | CON_1 | 11 | 346.273 | 277.140 | Two-way Mixed ANOVA<br>Between-subjects factor: AAV (CON vs. miR-124)<br>Within-subjects factor: Time (multiple time points) | AAV: $F(1,19)=0.493, p=0.491$<br>Time: $F(4.725,209)=2.921, p=0.019$<br>AAV*Time: $F(4.725,209)=0.452, p=0.801$ | Two-tailed Student's t-test | $t(19)=0.657, p=0.519199$ |
| | | | | miR-124_1 | 10 | 278.300 | 181.945 | | | | |
| | | | | CON_2 | 11 | 217.000 | 123.354 | | | Two-tailed Student's t-test | $t(19)=-0.350, p=0.729865$ |
| | | | | miR-124_2 | 10 | 238.200 | 153.503 | | | | |
| | | | | CON_3 | 11 | 205.818 | 170.172 | | | Two-tailed Student's t-test | $t(19)=0.483, p=0.634901$ |
| | | | | miR-124_3 | 10 | 174.400 | 121.217 | | | | |
| | | | | CON_4 | 11 | 217.545 | 149.349 | | | Two-tailed Student's t-test | $t(19)=1.193, p=0.247415$ |
| | | | | miR-124_4 | 10 | 144.900 | 127.259 | | | | |
| | | | | CON_5 | 11 | 205.818 | 181.989 | | | Two-tailed Student's t-test | $t(19)=0.159, p=0.875679$ |
| | | | | miR-124_5 | 10 | 193.700 | 166.675 | | | | |
| | | | | CON_6 | 11 | 223.091 | 178.445 | | | Two-tailed Student's t-test | $t(19)=0.216, p=0.830998$ |
| | | | | miR-124_6 | 10 | 208.600 | 119.225 | | | | |
| | | | | CON_7 | 11 | 227.000 | 201.826 | | | Two-tailed Student's t-test | $t(19)=1.029, p=0.316487$ |
| | | | | miR-124_7 | 10 | 153.800 | 103.539 | | | | |
| | | | | CON_8 | 11 | 231.000 | 182.914 | | | Two-tailed Student's t-test | $t(19)=0.749, p=0.462935$ |
| | | | | miR-124_8 | 10 | 177.400 | 139.395 | | | | |
| | | | | CON_9 | 11 | 189.273 | 147.816 | | | Two-tailed Student's t-test | $t(19)=0.956, p=0.351021$ |
| | | | | miR-124_9 | 10 | 131.400 | 127.425 | | | | |
| | | | | CON_10 | 11 | 227.000 | 177.810 | | | Two-tailed Student's t-test | $t(19)=0.692, p=0.497098$ |
| | | | | miR-124_10 | 10 | 180.100 | 124.961 | | | | |
| | | | | CON_11 | 11 | 251.818 | 188.683 | | | Two-tailed Welch's t-test | $t(16.135)=0.807, p=0.431155$ |
| | | | | miR-124_11 | 10 | 198.300 | 107.551 | | | | |
| | | | | CON_12 | 11 | 169.091 | 157.142 | | | Two-tailed Student's t-test | $t(19)=0.015$ $p=0.988499$ |
| | | | | miR-124_12 | 10 | 168.200 | 117.079 | | | | |
| | | | Saline:<br>miR-124 AAV + mCherry AAV vs. miR-124 AAV + GluA2 AAV | GluA2_1 | 11 | 259.364 | 150.571 | Two-way Mixed ANOVA<br>Between-subjects factor: Intervention (mCherry vs. GluA2)<br>Within-subjects factor: Time (multiple time points) | Intervention: $F(1,19)=0.007, p=0.933$<br>Time: $F(5.760,209)=3.406, p=0.005$<br>Intervention*Time: $F(5.760,209)=0.841, p=0.537$ | Two-tailed Student's t-test | $t(19)=0.261$ $p=0.797045$ |
| | | | | mCherry_1 | 10 | 278.300 | 181.945 | | | | |
| | | | | GluA2_2 | 11 | 242.818 | 133.405 | | | Two-tailed Student's t-test | $t(19)=-0.074$ $p=0.941964$ |
| | | | | mCherry_2 | 10 | 238.200 | 153.503 | | | | |
| | | | | GluA2_3 | 11 | 234.273 | 161.935 | | | Two-tailed Student's t-test | $t(19)=-0.951$ $p=0.353542$ |
| | | | | mCherry_3 | 10 | 174.400 | 121.217 | | | | |
| | | | | GluA2_4 | 11 | 210.364 | 184.555 | | | Two-tailed Student's t-test | $t(19)=-0.936$ $p=0.360792$ |
| | | | | mCherry_4 | 10 | 144.900 | 127.259 | | | | |
| | | | | GluA2_5 | 11 | 171.727 | 109.143 | | | Two-tailed Welch's t-test | $t(15.293)=0.354, p=0.728452$ |
| | | | | mCherry_5 | 10 | 193.700 | 166.675 | | | | |
| | | | | GluA2_6 | 11 | 178.273 | 148.690 | | | Two-tailed Student's t-test | $t(19)=0.512, p=0.614468$ |
| | | | | mCherry_6 | 10 | 208.600 | 119.225 | | | | |
| | | | | GluA2_7 | 11 | 158.909 | 128.932 | | | Two-tailed Student's t-test | $t(19)=-0.099, p=0.921831$ |
| | | | | mCherry_7 | 10 | 153.800 | 103.539 | | | | |
| | | | | GluA2_8 | 11 | 162.455 | 129.383 | | | Two-tailed Student's t-test | $t(19)=0.255, p=0.801577$ |
| | | | | mCherry_8 | 10 | 177.400 | 139.395 | | | | |
| | | | | GluA2_9 | 11 | 177.818 | 114.422 | | | Two-tailed Student's t-test | $t(19)=-0.880, p=0.389979$ |
| | | | | mCherry_9 | 10 | 131.400 | 127.425 | | | | |
| | | | | GluA2_10 | 11 | 154.000 | 137.722 | | | Two-tailed Student's t-test | $t(19)=0.453, p=0.655598$ |
| | | | | mCherry_10 | 10 | 180.100 | 124.961 | | | | |
| | | | | GluA2_11 | 11 | 214.636 | 153.831 | | | Two-tailed Student's t-test | $t(19)=-0.279, p=0.783113$ |
| | | | | mCherry_11 | 10 | 198.300 | 107.551 | | | | |
| | | | | GluA2_12 | 11 | 130.455 | 87.581 | | | Two-tailed Student's t-test | $t(19)=0.842, p=0.410344$ |
| | | | | mCherry_12 | 10 | 168.200 | 117.079 | | | | |
| | | | AMPH:<br>CON AAV + mCherry AAV vs. miR-124 AAV + mCherry AAV | CON_1 | 11 | 452.818 | 250.517 | Two-way Mixed ANOVA<br>Between-subjects factor: AAV (CON vs. miR-124)<br>Within-subjects factor: Time (multiple time points) | AAV: $F(1,19)=8.990, p=0.007$<br>Time: $F(2.270,209)=26.845, p=8.53E-9$<br>AAV*Time: $F(2.270,209)=3.920, p=0.023$ | Two-tailed Welch's t-test | $t(12.778)=-3.423, p=0.004638$ |
| | | | | miR-124_1 | 10 | 1065.900 | 513.478 | | | | |
| | | | | CON_2 | 11 | 487.545 | 297.772 | | | Two-tailed Welch's t-test | $t(13.029)=-3.525, p=0.003722$ |
| | | | | miR-124_2 | 10 | 1216.800 | 589.475 | | | | |
| | | | | CON_3 | 11 | 529.727 | 346.878 | | | Two-tailed Student's t-test | $t(19)=-3.403, p=0.002986$ |
| | | | | miR-124_3 | 10 | 1232.300 | 581.152 | | | | |
| | | | | CON_4 | 11 | 579.818 | 417.701 | | | Two-tailed Student's t-test | $t(19)=-2.707, p=0.013966$ |
| | | | | miR-124_4 | 10 | 1186.900 | 601.799 | | | | |
| | | | | CON_5 | 11 | 553.545 | 407.442 | | | Two-tailed Student's t-test | $t(19)=-2.798, p=0.011479$ |
| | | | | miR-124_5 | 10 | 1184.800 | 615.212 | | | | |
| | | | | CON_6 | 11 | 534.909 | 356.515 | | | Two-tailed Student's t-test | $t(19)=-2.622, p=0.016765$ |
| | | | | miR-124_6 | 10 | 1095.100 | 602.829 | | | | |
| | | | | CON_7 | 11 | 444.727 | 333.129 | | | Two-tailed Student's t-test | $t(19)=-2.172, p=0.042736$ |
| | | | | miR-124_7 | 10 | 902.000 | 605.732 | | | | |
| | | | | CON_8 | 11 | 306.455 | 206.465 | | | Two-tailed Welch's t-test | $t(10.888)=-2.206, p=0.049826$ |
| | | | | miR-124_8 | 10 | 751.200 | 606.457 | | | | |
| | | | | CON_9 | 11 | 224.273 | 161.053 | | | Two-tailed Welch's t-test | $t(10.338)=-2.531, p=0.029123$ |
| | | | | miR-124_9 | 10 | 691.200 | 562.767 | | | | |
| | | | | CON_10 | 11 | 188.182 | 99.585 | | | Two-tailed Welch's t-test | $t(9.723)=-2.621, p=0.026135$ |
| | | | | miR-124_10 | 10 | 588.700 | 473.895 | | | | |
| | | | | CON_11 | 11 | 171.909 | 119.705 | | | Two-tailed Student's t-test | $t(19)=-1.652, p=0.115062$ |
| | | | | miR-124_11 | 10 | 413.100 | 468.965 | | | | |
| | | | | CON_12 | 11 | 215.818 | 114.806 | | | Two-tailed Student's t-test | $t(19)=-1.368, p=0.187329$ |
| | | | | miR-124_12 | 10 | 339.500 | 275.262 | | | | |
| | | | AMPH:<br>miR-124 AAV + mCherry AAV vs. miR-124 AAV + GluA2 AAV | GluA2_1 | 11 | 400.364 | 192.066 | Two-way Mixed ANOVA with the Greenhouse-Geisser correction<br>Between-subjects factor: Intervention (mCherry vs. GluA2)<br>Within-subjects factor: Time (multiple time points) | Intervention: $F(1,19)=1.485, p=0.238$<br>Time: $F(2.593,49.262)=18.865, p=5.772E-9$<br>Intervention*Time: $F(2.593,49.262)=6.301, p=0.002$ | Two-tailed Welch's t-test | $t(11.271)=3.861, p=0.002536$ |
| | | | | mCherry_1 | 10 | 1065.900 | 513.478 | | | | |
| | | | | GluA2_2 | 11 | 624.818 | 336.766 | | | Two-tailed Welch's t-test | $t(14.022)=2.789, p=0.014478$ |
| | | | | mCherry_2 | 10 | 1216.800 | 589.475 | | | | |
| | | | | GluA2_3 | 11 | 745.273 | 437.220 | | | Two-tailed Student's t-test | $t(19)=2.184, p=0.041742$ |
| | | | | mCherry_3 | 10 | 1232.300 | 581.152 | | | | |
| | | | | GluA2_4 | 11 | 809.909 | 446.323 | | | Two-tailed Student's t-test | $t(19)=1.641, p=0.117213$ |
| | | | | mCherry_4 | 10 | 1186.900 | 601.799 | | | | |
| | | | | GluA2_5 | 11 | 845.091 | 483.247 | | | Two-tailed Student's t-test | $t(19)=1.414, p=0.173442$ |
| | | | | mCherry_5 | 10 | 1184.800 | 615.212 | | | | |
| | | | | GluA2_6 | 11 | 844.455 | 523.628 | | | Two-tailed Student's t-test | $t(19)=1.020, p=0.320654$ |
| | | | | mCherry_6 | 10 | 1095.100 | 602.829 | | | | |
| | | | | GluA2_7 | 11 | 797.727 | 507.484 | | | Two-tailed Student's t-test | $t(19)=0.429, p=0.672694$ |
| | | | | mCherry_7 | 10 | 902.000 | 605.732 | | | | |
| | | | | GluA2_8 | 11 | 724.182 | 424.247 | | | Two-tailed Student's t-test | $t(19)=0.119, p=0.906339$ |
| | | | | mCherry_8 | 10 | 751.200 | 606.457 | | | | |
| | | | | GluA2_9 | 11 | 613.273 | 468.852 | | | Two-tailed Student's t-test | $t(19)=0.346, p=0.733149$ |
| | | | | mCherry_9 | 10 | 691.200 | 562.767 | | | | |
| | | | | GluA2_10 | 11 | 596.909 | 430.061 | | | Two-tailed Student's t-test | $t(19)=-0.042, p=0.967231$ |
| | | | | mCherry_10 | 10 | 588.700 | 473.895 | | | | |
| | | | | GluA2_11 | 11 | 501.636 | 416.399 | | | Two-tailed Student's t-test | $t(19)=-0.458, p=0.651892$ |
| | | | | mCherry_11 | 10 | 413.100 | 468.965 | | | | |
| | | | | GluA2_12 | 11 | 373.273 | 417.000 | | | Two-tailed Student's t-test | $t(19)=-0.217, p=0.830871$ |
| | | | | mCherry_12 | 10 | 339.500 | 275.262 | | | | |

#### Figure S2

| Figure | Test | Measurement | Group | Sub-group | Sample size | Average | Standard deviation | Statistical test | Statistical result | *post hoc* test | Result from the *post hoc* tests |
|---|---|---|---|---|---|---|---|---|---|---|---|
| S2A | Expression analysis for pri-miR-124-2 | Relative expression (A.U.) | CON | - | 225 | -0.117 | 0.85503 | One-way ANOVA | $F(2,415)=3.552, p=0.030$ | Tukey's HSD (CON vs. SZ) | $p=0.030$ |
| | | | SZ | | 143 | 0.148 | 1.01757 | | | Tukey's HSD (CON vs. BP) | $p=0.325$ |
| | | | BP | | 50 | 0.101 | 1.28960 | | | Tukey's HSD (SZ vs. BP) | $p=0.953$ |
| S2B | Expression analysis for pre-miR-124-2 | Relative expression (A.U.) | CON | - | 225 | -0.101 | 0.951 | One-way ANOVA | $F(2,415)=4.580, p=0.011$ | Tukey's HSD (CON vs. SZ) | $p=0.038$ |
| | | | SZ | | 143 | 0.0365 | 0.93745 | | | Tukey's HSD (CON vs. BP) | $p=0.009$ |
| | | | BP | | 50 | 0.350 | 1.133 | | | Tukey's HSD (SZ vs. BP) | $p=0.382$ |

#### Figure S5

| Figure | Test | Measurement | Group | Sub-group | Sample size | Average | Standard deviation | Statistical test | Statistical result | *post hoc* test | Result from the *post hoc* tests |
|---|---|---|---|---|---|---|---|---|---|---|---|
| S5B | Three-chmaber social interaction test (trial1): sociability test | Investigation time (s) | CON AAV | Non-social | 8 | 29.763 | 6.648 | Two-way Mixed ANOVA<br>Between-subjects factor: AAV (CON v.s miR-14)<br>Within-subjects factor: Chamber (non-social vs. social) | AAV: $F(1,14)=4.527$, $p=0.052$<br>Chamber: $F(1,14)=76.664$, $p=4.716E-07$<br>AAV*Chamber: $F(1,14)=14.829$, $p=0.002$ | Two-tailed paired Student's t-test | $t(7)=6.864$, $p=0.000239$ |
| | | | | Social | 8 | 56.525 | 14.301 | | | | |
| | | | miR-124 AAV | Non-social | 8 | 27.688 | 10.548 | | | Two-tailed paired Student's t-test | $t(7)=6.195$, $p=0.000447$ |
| | | | | Social | 8 | 38.100 | 9.138 | | | | |
| | Three-chmaber social interaction test (trial2): social recognition test | Investigation time (s) | CON AAV | Familiar | 8 | 19.088 | 5.498 | Two-way Mixed ANOVA<br>between-subjects factor: AAV (CON v.s miR-14)<br>within-subjects factor: Chamber (familiar vs. novel) | AAV: $F(1,14)=0.920$, $p=0.354$<br>Chamber: $F(1,14)=38.336$, $p=0.000023$<br>AAV*Chamber: $F(1,14)=4.614$, $p=0.049693$ | Two-tailed paired Student's t-test | $t(7)=-5.014$, $p=0.001541$ |
| | | | | Novel | 8 | 45.488 | 17.306 | | | | |
| | | | miR-124 AAV | Familiar | 8 | 20.788 | 8.643 | | | Two-tailed paired Student's t-test | $t(7)=-3.641$, $p=0.008281$ |
| | | | | Novel | 8 | 33.588 | 14.436 | | | | |
| S5C | Amphetamine challenge | Locomotion activity (counts / 5 min) - left | Habituation | CON_1 | 8 | 608.000 | 237.230 | Two-way Mixed ANOVA<br>Between-subjects factor: AAV (CON vs. miR-124)<br>Within-subjects factor: Time (multiple time points) | AAV: $F(1,14)=0.079$, $p=0.783$<br>Time: $F(11,154)=10.429$, $p=3.909E-14$<br>AAV*Time: $F(11,154)=0.757$, $p=0.682$ | Two-tailed Student's t-test | $t(14)=-0.747$, $p=0.467287$ |
| | | | | miR-124_1 | 8 | 685.500 | 172.561 | | | | |
| | | | | CON_2 | 8 | 494.000 | 188.724 | | | Two-tailed Student's t-test | $t(14)=0.232$, $p=0.820218$ |
| | | | | miR-124_2 | 8 | 473.125 | 171.435 | | | | |
| | | | | CON_3 | 8 | 394.875 | 133.336 | | | Two-tailed Student's t-test | $t(14)=-1.582$, $p=0.135920$ |
| | | | | miR-124_3 | 8 | 490.000 | 105.535 | | | | |
| | | | | CON_4 | 8 | 404.500 | 104.973 | | | Two-tailed Student's t-test | $t(14)=-0.136$, $p=0.893538$ |
| | | | | miR-124_4 | 8 | 412.125 | 118.421 | | | | |
| | | | | CON_5 | 8 | 392.125 | 145.467 | | | Two-tailed Student's t-test | $t(14)=0.008$, $p=0.993506$ |
| | | | | miR-124_5 | 8 | 391.625 | 89.267 | | | | |
| | | | | CON_6 | 8 | 454.500 | 117.648 | | | Two-tailed Student's t-test | $t(14)=0.819$, $p=0.426522$ |
| | | | | miR-124_6 | 8 | 410.625 | 95.496 | | | | |
| | | | | CON_7 | 8 | 429.125 | 119.028 | | | Two-tailed Student's t-test | $t(14)=0.423$, $p=0.679048$ |
| | | | | miR-124_7 | 8 | 403.125 | 126.974 | | | | |
| | | | | CON_8 | 8 | 350.000 | 131.848 | | | Two-tailed Student's t-test | $t(14)=0.079$, $p=0.938006$ |
| | | | | miR-124_8 | 8 | 344.750 | 133.348 | | | | |
| | | | | CON_9 | 8 | 303.750 | 181.138 | | | Two-tailed Student's t-test | $t(14)=-0.861$, $p=0.403956$ |
| | | | | miR-124_9 | 8 | 378.125 | 164.132 | | | | |
| | | | | CON_10 | 8 | 354.500 | 171.054 | | | Two-tailed Student's t-test | $t(14)=-0.381$, $p=0.708615$ |
| | | | | miR-124_10 | 8 | 384.125 | 137.844 | | | | |
| | | | | CON_11 | 8 | 310.375 | 182.456 | | | Two-tailed Student's t-test | $t(14)=-0.387$, $p=0.704568$ |
| | | | | miR-124_11 | 8 | 344.000 | 164.622 | | | | |
| | | | | CON_12 | 8 | 388.125 | 195.787 | | | Two-tailed Student's t-test | $t(14)=0.356$, $p=0.727414$ |
| | | | | miR-124_12 | 8 | 356.375 | 159.467 | | | | |
| | | | Saline | CON_1 | 8 | 349.125 | 272.460 | Two-way Mixed ANOVA<br>Between-subjects factor: AAV (CON vs. miR-124)<br>Within-subjects factor: Time (multiple time points) | AAV: $F(1,14)=0.785$, $p=0.391$<br>Time: $F(11,154)=3.268$, $p=4.832E-04$<br>AAV*Time: $F(11,154)=1.076$, $p=0.384$ | Two-tailed Welch's t-test | $t(7.886)=0.439$, $p=0.672349$ |
| | | | | miR-124_1 | 8 | 305.500 | 68.692 | | | | |
| | | | | CON_2 | 8 | 293.875 | 112.680 | | | Two-tailed Student's t-test | $t(14)=1.489$, $p=0.158613$ |
| | | | | miR-124_2 | 8 | 207.750 | 118.577 | | | | |
| | | | | CON_3 | 8 | 189.625 | 90.974 | | | Two-tailed Student's t-test | $t(14)=-0.472$, $p=0.644129$ |
| | | | | miR-124_3 | 8 | 209.375 | 75.661 | | | | |
| | | | | CON_4 | 8 | 296.750 | 132.220 | | | Two-tailed Student's t-test | $t(14)=1.234$, $p=0.237369$ |
| | | | | miR-124_4 | 8 | 205.500 | 161.967 | | | | |
| | | | | CON_5 | 8 | 320.875 | 85.519 | | | Two-tailed Student's t-test | $t(14)=3.244$, $p=0.005879$ |
| | | | | miR-124_5 | 8 | 150.375 | 121.580 | | | | |
| | | | | CON_6 | 8 | 205.375 | 173.555 | | | Two-tailed Welch's t-test | $t(0.285)=11.084$, $p=0.780708$ |
| | | | | miR-124_6 | 8 | 185.250 | 98.478 | | | | |
| | | | | CON_7 | 8 | 278.875 | 141.613 | | | Two-tailed Student's t-test | $t(14)=0.043$, $p=0.966292$ |
| | | | | miR-124_7 | 8 | 276.375 | 83.428 | | | | |
| | | | | CON_8 | 8 | 252.375 | 168.358 | | | Two-tailed Student's t-test | $t(14)=-0.299$, $p=0.769466$ |
| | | | | miR-124_8 | 8 | 275.750 | 143.554 | | | | |
| | | | | CON_9 | 8 | 225.625 | 97.115 | | | Two-tailed Student's t-test | $t(14)=1.181$, $p=0.257306$ |
| | | | | miR-124_9 | 8 | 169.875 | 91.640 | | | | |
| | | | | CON_10 | 8 | 184.750 | 154.476 | | | Two-tailed Student's t-test | $t(14)=1.053$, $p=0.310025$ |
| | | | | miR-124_10 | 8 | 111.500 | 121.755 | | | | |
| | | | | CON_11 | 8 | 205.375 | 149.902 | | | Two-tailed Student's t-test | $t(14)=0.269$, $p=0.791750$ |
| | | | | miR-124_11 | 8 | 182.250 | 191.288 | | | | |
| | | | | CON_12 | 8 | 190.875 | 199.597 | | | Two-tailed Student's t-test | $t(14)=-0.044$, $p=0.965338$ |
| | | | | miR-124_12 | 8 | 195.000 | 172.378 | | | | |
| | | | AMPH | CON_1 | 8 | 264.500 | 189.299 | Two-way Mixed ANOVA<br>Between-subjects factor: AAV (CON vs. miR-124)<br>Within-subjects factor: Time (multiple time points) | For the first 60 min,<br>AAV: $F(1,14)=12.924$, $p=0.003$<br>Time: $F(11,154)=4.461$, $p=8E-6$<br>AAV*Time: $F(11,154)=1.138$, $p=0.336$<br><br>For the first 15 min,<br>AAV: $F(1,14)=9.425$, $p=0.008$<br>Time: $F(2,28)=7.404$, $p=0.003$<br>AAV*Time: $F(2,28)=4.369$, $p=0.022$ | Two-tailed Student's t-test | $t(14)=-2.438$, $p=0.029$ |
| | | | | miR-124_1 | 8 | 460.625 | 126.198 | | | | |
| | | | | CON_2 | 8 | 188.750 | 112.988 | | | Two-tailed Student's t-test | $t(14)=-2.322$, $p=0.036$ |
| | | | | miR-124_2 | 8 | 381.125 | 205.251 | | | | |
| | | | | CON_3 | 8 | 232.500 | 174.930 | | | Two-tailed Student's t-test | $t(14)=-3.392$, $p=0.004$ |
| | | | | miR-124_3 | 8 | 606.375 | 258.064 | | | | |
| | | | | CON_4 | 8 | 336.500 | 202.399 | | | Two-tailed Student's t-test | $t(14)=-2.536$, $p=0.024$ |
| | | | | miR-124_4 | 8 | 629.375 | 256.419 | | | | |
| | | | | CON_5 | 8 | 334.625 | 228.070 | | | Two-tailed Student's t-test | $t(14)=-2.281$, $p=0.039$ |
| | | | | miR-124_5 | 8 | 676.250 | 357.074 | | | | |
| | | | | CON_6 | 8 | 277.250 | 153.090 | | | Two-tailed Student's t-test | $t(14)=-2.896$, $p=0.012$ |
| | | | | miR-124_6 | 8 | 666.625 | 348.128 | | | | |
| | | | | CON_7 | 8 | 234.875 | 134.829 | | | Two-tailed Welch's t-test | $t(9.156)=-2.621$, $p=0.027$ |
| | | | | miR-124_7 | 8 | 573.250 | 339.377 | | | | |
| | | | | CON_8 | 8 | 238.500 | 93.680 | | | Two-tailed Welch's t-test | $t(8.280)=-3.055$, $p=0.015$ |
| | | | | miR-124_8 | 8 | 586.750 | 308.554 | | | | |
| | | | | CON_9 | 8 | 212.750 | 139.128 | | | Two-tailed Welch's t-test | $t(9.292)=-2.071$, $p=0.067$ |
| | | | | miR-124_9 | 8 | 481.125 | 339.045 | | | | |
| | | | | CON_10 | 8 | 173.125 | 155.570 | | | Two-tailed Student's t-test | $t(14)=-2.708$, $p=0.017$ |
| | | | | miR-124_10 | 8 | 415.750 | 200.002 | | | | |
| | | | | CON_11 | 8 | 191.250 | 105.795 | | | Two-tailed Student's t-test | $t(14)=-1.270$, $p=0.225$ |
| | | | | miR-124_11 | 8 | 285.000 | 180.054 | | | | |
| | | | | CON_12 | 8 | 170.625 | 134.808 | | | Two-tailed Student's t-test | $t(14)=-2.365$, $p=0.033$ |
| | | | | miR-124_12 | 8 | 381.000 | 212.395 | | | | |
| S5D | Prepulse inhibition | p74 - PPI (%) | CON | - | 4 | 48.988 | 18.8305 | Two-way Mixed ANOVA<br>Between-subjects factor: AAV<br>Within-subjects factor: Sound (p74,p78, p82, p86, p90) | AAV: $F(1,7)=1.266$, $p=0.298$<br>Sound: $F(4,28)=5.640$, $p=0.002$<br>AAV*Sound: $F(4,28)=0.705$, $p=0.595$ | Two-tailed Student's t-test | $t(8)=0.194$, $p=0.851307$ |
| | | | miR-124 | | 5 | 49.972 | 18.7339 | | | | |
| | | p78 - PPI (%) | CON | - | 4 | 54.398 | 14.1930 | | | Two-tailed Student's t-test | $t(7)=-1.559$, $p=0.163060$ |
| | | | miR-124 | | 5 | 64.964 | 5.2598 | | | | |
| | | p82 - PPI (%) | CON | - | 4 | 62.453 | 17.3506 | | | Two-tailed Student's t-test | $t(8)=-0.838$, $p=0.426093$ |
| | | | miR-124 | | 5 | 75.738 | 13.8271 | | | | |
| | | p86 - PPI (%) | CON | - | 4 | 61.123 | 7.9608 | | | Two-tailed Welch's t-test | $t(7.583)=-0.641$, $p=0.540419$ |
| | | | miR-124 | | 5 | 69.318 | 16.0767 | | | | |
| | | p90 - PPI (%) | CON | - | 4 | 57.593 | 19.6765 | | | Two-tailed Student's t-test | $t(8)=-1.026$, $p=0.335033$ |
| | | | miR-124 | | 5 | 72.154 | 14.7837 | | | | |
| S5E | Forced swim test | Immobility (%) | CON | - | 8 | 52.307 | 7.110 | - | - | Two-tailed Welch's t-test | $t(11.342)=0.0.031$, $p=0.976$ |
| | | | miR-124 | | 8 | 52.152 | 12.060 | | | | |

#### Figure S6

| Figure | Test | Measurement | Group | Sub-group | Sample size | Average | Standard deviation | Statistical test | Statistical result | *post hoc* test | Result from the *post hoc* tests |
|---|---|---|---|---|---|---|---|---|---|---|---|
| S6B | Surface AMPARs (sGluA1 and sGluA2) intensities | sGluA1 | CON | - | 27 | 100.000 | 20.411 | - | - | Two-tailed Student's t-test | $t(50)=-1.066, p=0.291$ |
| | | | miR-124 | | 25 | 107.100 | 27.336 | | | | |
| | | sGluA2 | CON | - | 40 | 100.000 | 30.867 | - | - | Two-tailed Welch's t-test | $t(55.262)=4.774, p=0.000014$ |
| | | | miR-124 | | 37 | 74.246 | 13.976 | | | | |
